# A transgene-free, human peri-gastrulation embryo model with trilaminar embryonic disc-, amnion- and yolk sac-like structures

**DOI:** 10.1101/2024.08.05.606556

**Authors:** Shiyu Sun, Yi Zheng, Yung Su Kim, Zheng Zhong, Norio Kobayashi, Xufeng Xue, Yue Liu, Zhuowei Zhou, Yanhong Xu, Jinglei Zhai, Hongmei Wang, Jianping Fu

## Abstract

The ultimate outcome of the gastrulation in mammalian development is a recognizable trilaminar disc structure containing organized cell lineages with spatially defined identities in an emerging coordinate system^1–4^. Despite its importance in human development, gastrulation remains difficult to study. Stem cell-based embryo models, including those that recapitulate different aspects of pre- and peri-gastrulation human development^5–15^, are emerging as promising tools for studying human embryogenesis^16–18^. However, it remains unclear whether existing human embryo models are capable of modeling the development of the trilaminar embryonic disc structure, a hallmark of human gastrulation. Here we report a transgene-free human embryo model derived solely from primed human pluripotent stem cells (hPSCs), which recapitulates various aspects of peri-gastrulation human development, including formation of trilaminar embryonic layers situated between dorsal amnion and ventral definitive yolk sac and primary hematopoiesis. We term this model the peri-gastrulation trilaminar embryonic disc (PTED) embryoid. The development of PTED embryoid does not follow natural developmental sequences of cell lineage diversification or spatial organization. Instead, it exploits both extrinsic control of tissue boundaries and intrinsic self-organizing properties and embryonic plasticity of the diverse peri-gastrulation-stage cell lineages, leading to the emergence of *in vivo*-like tissue organization and function at a global scale. Our lineage tracing study reveals that in PTED embryoids, embryonic and extraembryonic mesoderm cells, as well as embryonic and extraembryonic endoderm cells, share common progenitors emerging during peri-gastrulation development. Active hematopoiesis and blood cell generation are evident in the yolk sac-like structure of PTED embryoids. Together, PTED embryoids provide a promising and ethically less challenging model for studying self-organizing properties of peri-gastrulation human development.

---

Gastrulation is considered as one of the most crucial milestones in mammalian development^1–4^. Through gastrulation, a homogeneous population of pluripotent epiblast cells in the bilaminar disc of human embryo self-organizes to form the trilaminar embryonic disc, consisting of embryonic ectoderm, mesoderm, and endoderm along the dorsal (*D*)-ventral (*V*) axis, flanked by dorsal amnion and ventral definitive yolk sac^19^ (**Fig. 1a**, **Extended Data Fig. 1a**). Gastrulation thus leads to the formation of the basic body plan in tandem with a dramatic change in morphology. The final outcome of gastrulation is a distinct structure with organized cell lineages that have spatially defined identities, establishing the foundation for organogenesis. Despite its importance, direct study of human gastrulation remains challenging both technically and ethically. Thus, it remains unclear how the specification and organization of the embryonic germ layers are choreographed during human gastrulation. It also remains an open question how embryonic and extraembryonic lineages are segregated and allocated to form various embryonic and extraembryonic structures during human gastrulation. As a unique feature of primate embryogenesis, the definitive yolk sac emerges during human gastrulation. Despite its role as a multifunctional hub for hematopoiesis and nutritional supply, the origin of extraembryonic lineages comprising the definitive yolk sac remains elusive^20–22^.

**Figure 1.**
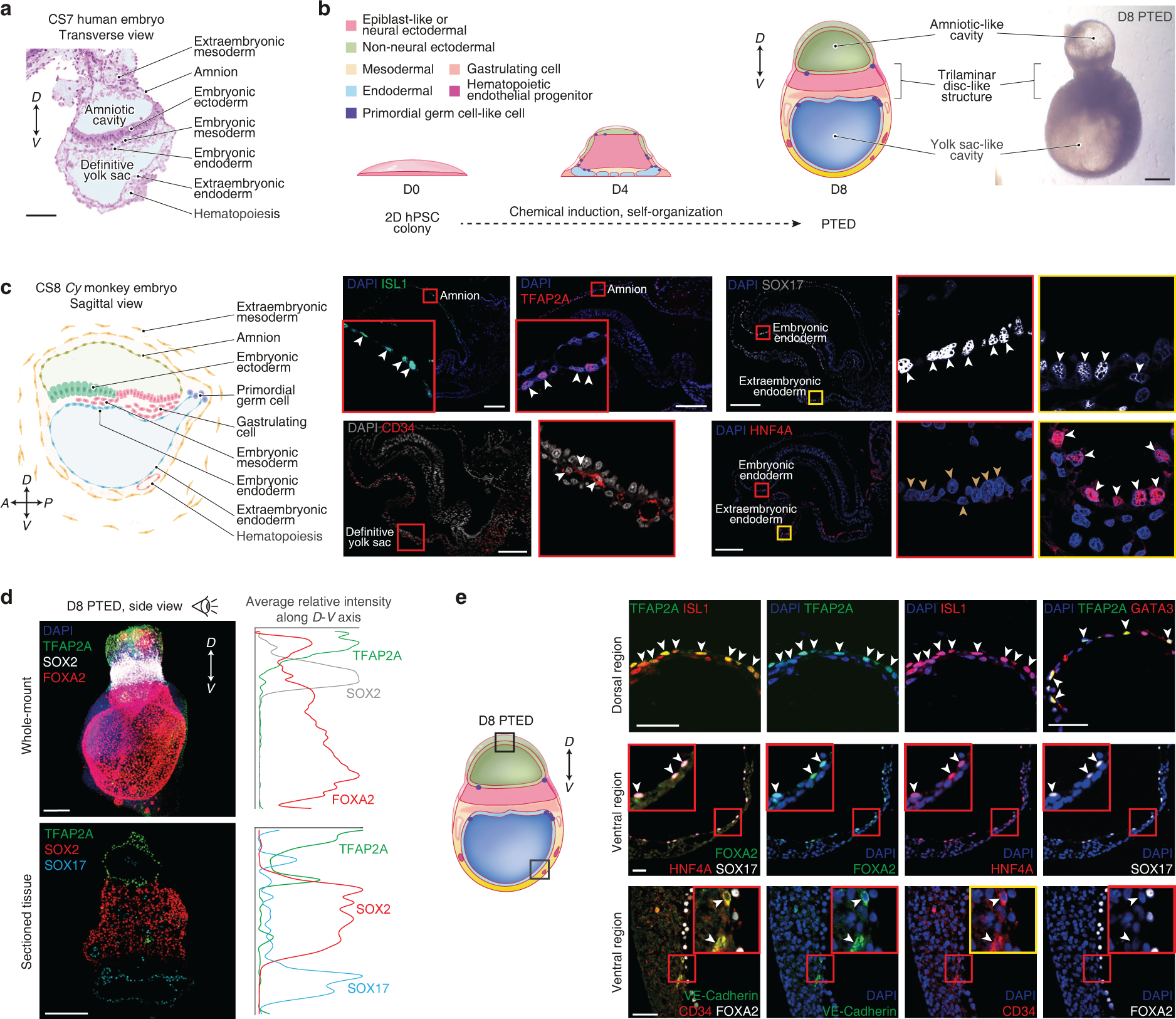
A peri-gastrulation trilaminar embryonic disc (PTED) embryoid with amnion- and yolk sac-like structures. **a.** H&E staining of a Carnegie stage (CS) 7 human embryo (https://www.ehd.org/virtual-human-embryo/). *D*, dorsal; *V*, ventral. Scale bar, 100 µm. **b.** Left: Cartoons of PTED embryoids on different days (D). Different colored regions in PTED embryoids mark distinct cellular compartments as indicated. See **Methods** for PTED protocol. Right: Representative bright-field micrograph showing a D8 PTED embryoid. Scale bar, 200 µm. **c.** Left: Schematic of CS8 *Cynomolgus* (*Cy)* monkey embryo. Right: Representative confocal micrographs of a *Cy* monkey embryo on embryonic day 22 (E22 or CS8) stained for indicated lineage markers. Magnified views are included for regions highlighted by colored boxes. White arrowheads mark ISL1^+^, TFAP2A^+^, HNF4A^+^, SOX17^+^ or CD34^+^ cells, respectively. Brown arrowheads label HNF4A^-^ cells. DAPI counterstains cell nuclei. *A*, anterior; *P*, posterior. Scale bars, 200 µm. **d.** Top: Representative light-sheet micrograph of whole-mount D8 PTED embryoid stained for indicated lineage markers. Bottom: Representative confocal micrograph of cross-sectioned D8 PTED embryoid stained for indicated lineage markers. Right: Plots showing average relative intensity of indicated marker expression along the *D*-*V* axis. DAPI counterstains cell nuclei. Scale bars, 200 µm. **e.** Left: Cartoon of D8 PTED embryoid, with boxes marking dorsal and ventral regions, where magnified views are provided on the right. Right: Representative confocal micrographs showing dorsal and ventral regions of D8 PTED embryoids stained for indicated markers. Further magnified views are included for regions highlighted by red boxes. White arrowheads mark TFAP2A^+^ISL1^+^ and TFAP2A^+^GATA3^+^ cells (*top*), FOXA2^+^HNF4A^+^SOX17^+^ cells (*middle*), and VE-Cadherin^+^CD34^+^ cells (*bottom*), respectively. DAPI counterstains cell nuclei. Scale bars, 50 µm.

Stem cell-based human embryo models, or embryoids, are emerging as promising tools to promote fundamental understanding of human development and advance reproductive and regenerative medicine^16–18^. Early human embryoid studies aim to model germ layer lineage specification and organization based on patterned two-dimensional (2D) colonies of primed human pluripotent stem cells (hPSCs)^5^. Three-dimensional (3D) human embryoids were also reported based on 3D cultures of primed hPSCs to recapitulate post-implantation amnion-epiblast patterning, primordial germ cell (PGC) development, and the onset of gastrulation at the posterior epiblast compartment^6^, or anterior (*A*)-posterior (*P*) symmetry breaking of the epiblast^7^. Following these studies, assembloid approaches were utilized to assemble primed hPSCs and extraembryonic-like cells to examine the role of embryonic-extraembryonic interactions in *A*-*P* symmetry breaking of the epiblast^23^. There are other peri-gastrulation human embryo models, *e.g.* gastruloids^8^ and trunk-like structures^24–27^, that can model axial patterning and caudal tissue development. Recently, 3D human embryoids were reported using hPSCs with naïve, extended or primed pluripotency states, sometimes combined with extraembryonic-like cells obtained through transcription factor overexpression, to recapitulate important aspects of human pre- and peri-gastrulation development^9–15^, including bilaminar disk formation, *A*-*P* symmetry breaking of the epiblast, yolk sac formation, and primary hematopoiesis. However, it remains unclear whether these human embryoids are capable of modeling the stereotypic organization of the trilaminar embryonic disc structure, a hallmark of gastrulation in vertebrate species^1–4^.

In this study, we report a transgene-free human embryoid derived solely from primed hPSCs that recapitulates different aspects of the peri-gastrulation human development, including formations of trilaminar disc structure flanked by dorsal amnion and ventral definitive yolk sac (**Fig. 1b**). We term this model the peri-gastrulation trilaminar embryonic disc (PTED) embryoid. The development of PTED embryoid does not follow natural developmental sequences of cell lineage diversification or spatial organization. Instead, it exploits extrinsic bioengineering control of tissue boundaries to establish an effective morphogenetic field to promote cellular organization during embryoid development, as well as intrinsic self-organizing properties and embryonic plasticity of the diverse peri-gastrulation-stage cell lineages, resulting in the emergence of *in vivo*-like tissue organization and function at a global scale. Specifically, our data show that even without external signals, the local cellular interactions among the diverse peri-gastrulation embryonic and extraembryonic lineages could lead to the formation of patterned structures resembling the trilaminar disc, amnion, and yolk sac in PTED embryoids. Cell lineage identities in PTED embryoids are validated through immunostaining for lineage markers and benchmarking against *in vivo* human and monkey gastrula at the transcriptomic level. Our data support that embryonic and extraembryonic mesoderm cells, as well as embryonic and extraembryonic endoderm cells, might share common progenitors that emerge during peri-gastrulation development. This notion suggests gastrulating epiblast as an alternative origin of these extraembryonic lineages. Furthermore, primary hematopoiesis and blood cell generation are evident in the definitive yolk sac-like structure of PTED embryoids. Together, the PTED embryoid provides a promising, non-canonical model for studying self-organizing properties of peri-gastrulation human development.

## RESULTS

### Development of PTED embryoid

To ascertain cell lineages potentially present in PTED embryoids, we first examined lineage markers associated with different embryonic and extraembryonic tissues in peri-gastrulation primate embryos (**Fig. 1c**, **Extended Data Fig. 1b-f**). In *in vivo*, peri-gastrulation *Cynomolgus* (*Cy*) monkey embryo (embryonic day 22 or E22, or Carnegie stage 8 or CS8), non-neural ectoderm (NNE, including amnion) is marked by TFAP2A, ISL1 and GATA3^28^, embryonic ectoderm retains OCT4 and SOX2 expression, gastrulating cells exhibit marked expression of BRACHYURY (or BRA), embryonic endoderm (EmEndo) upregulates SOX17 but is negative for HNF4A, and extraembryonic endoderm (ExEndo) upregulates HNF4A and exhibit a low level of SOX17 expression^29,30^ (**Fig. 1c**, **Extended Data Fig. 1b-f**). Thus, HNF4A serves as a marker useful for distinguishing peri-gastrulation EmEndo and ExEndo. CD34^+^VE-Cadherin^+^ hematopoietic endothelial progenitors (HEPs) are evident in the extraembryonic mesoderm (ExM) surrounding the definitive yolk sac^20^ (**Fig. 1c**, **Extended Data Fig. 1b-f**).

Studies of mammalian gastrulation, based on both mouse embryos^2,31–33^ and human embryoids^5,34,35^, show that a signaling cascade involving the Bone Morphogenic Protein (BMP), WNT and NODAL pathways is integral for initiating the gastrulation. Particularly, BMP signaling in extraembryonic tissues adjacent to the pre-gastrulation epiblast initiates the gastrulation signaling cascade, activating WNT followed by NODAL in the epiblast to trigger the onset of gastrulation^32,33^. To develop PTED embryoids, we thus applied exogenous BMP4 to treat primed hPSCs, which reside in a developmental state similar to post-implantation epiblast^36,37^ and have been successfully utilized for modelling peri-gastrulation human development^5–7,34,35,38^. Specifically, H9 human embryonic stem cells (hESCs) are first seeded onto circular Geltrex adhesive islands with a diameter of 800 µm printed onto glass coverslips using microcontact printing (see **Methods**; **Extended Data Fig. 2a&b**). Such geometric boundary confinement provides an effective morphogenetic field to promote cellular organization and tissue-tissue interaction during embryoid development^5,34,35,39,40^ and is utilized in 2D human embryoids to study germ layer differentiation and organization^5,35,41^. Under a cell seeding density of 5,000 cells mm^-2^, about 1,800 hESCs are attached to each adhesive island. Cells are cultured in mTeSR1 medium for the first two days before BMP4 is supplemented for the following two days to initiate hESC differentiation (**Extended Data Fig. 2a**). The day on which BMP4 is added into mTeSR1 is designated as Day 0 (**Extended Data Fig. 2a**).

Notably, BMP4 stimulations transform hESC colonies from a 2D monolayer to a 3D structure, even evident on Day 1 (**Extended Data Fig. 2c-i**). Two regions of TFAP2A^+^ NNE cells emerge on Day 1, with one located at the boundary and the other at the top central region of PTED embryoids (**Extended Data Fig. 2e**, **Supplementary Video 1**). On Day 2, BRA^+^ gastrulating-like cells emerge and form small clusters embedded inside PTED embryoids at discrete, peripheral regions (**Extended Data Fig. 2f**). The development of BRA^+^ gastrulating-like cells does not appear to involve the formation of a primitive streak-like structure. These cells co-express SNAIL, MIXL1, EOMES, or N-Cadherin (N-CAD), known gastrulation-related markers (**Extended Data Fig. 2g**). Even with exogeneous BMP4 being removed from culture medium from Day 3 onwards, hESC colonies continue to develop (**Extended Data Fig. 2h&i**, **Supplementary Video 2**). On Day 3, FOXA2^+^SOX17^+^ endodermal cells emerge in a concentric ring region at bottom surfaces of PTED embryoids (**Extended Data Fig. 2h**). Concomitantly, BRA^+^ gastrulating-like cells become organized in a similar way (**Extended Data Fig. 2h**). From Day 3 onwards, TFAP2A^+^ NNE cells become accumulated at top central regions of PTED embryoids (**Extended Data Fig. 2h&i**).

On Day 5, large cavities become evident in PTED embryoids, and some embryoids start to detach from underlying glass coverslips (**Extended Data Fig. 3a**). Immunofluorescence staining of Day 5 (D5) PTED embryoids reveals three distinct regions along their lengths, with one pole accumulated with TFAP2A^+^ISL1^+^ amnion cells (‘the dorsal pole’), the opposite pole containing ISL1^+^TFAP2A^-^ or ISL1^+^SNAIL^+^HAND2^+^ mesodermal and SOX17^+^FOXA2^+^ endodermal cells (‘the ventral pole’), and regions in-between occupied by OCT4^+^ and SOX2^+^ epiblast-like cells (**Extended Data Fig. 3b-d**). Ventral mesodermal regions often contain cavities (**Extended Data Fig. 3d**). In adjacent areas, smaller openings are often evident, and they are demarcated by a single layer of FOXA2^+^SOX17^+^ endodermal cells (**Extended Data Fig. 3e**). Notably, these cells show upregulated HNF4A, suggesting their ExEndo-like identity (**Extended Data Fig. 3e**). In contrast, FOXA2^+^SOX17^+^ endodermal cells near the central epiblast-like region are often HNF4A^-^ (**Extended Data Fig. 3e**). Together, around Day 5, the endodermal cell lineage in PTED embryoids bifurcates into embryonic- or extraembryonic-like fates, depending on their locations^29,30^. From Day 5 onwards, FOXA2^+^SOX17^+^HNF4A^+^ ExEndo-like cells form yolk sac-like cavities at the ventral pole of PTED embryoids (**Extended Data Fig. 3e**).

Between Day 5 and Day 8, most PTED embryoids become free-floating (**Extended Data Fig. 4a&b**, **Supplementary Video 3**). PTED embryoids grow in size and become elongated over time (**Extended Data Fig. 4c**). They exhibit a distinct tissue architecture characterized by the presence of two cavities at two ends flanking a central region containing densely packed cells (**Fig. 1b&d**, **Extended Data Fig. 4a-d**). Immunofluorescence staining of D8 PTED embryoids reveals TFAP2A^+^ NNE cells and SOX17^+^ or FOXA2^+^ endodermal cells lining the two cavities, respectively, and most of cells in-between retaining SOX2 or NANOG expression (**Fig. 1d**, **Extended Data Fig. 4d**, **Supplementary Video 4**). Further analysis shows that cavities at one pole of are surrounded by TFAP2A^+^ISL1^+^ or GATA3^+^ISL1^+^ amnion cells, supporting this structure as an amniotic-like cavity (**Fig. 1e**, **Extended Data Fig. 4e**). Thus, this pole is designated as the dorsal pole and the opposite pole ascribed as the ventral pole. Within dorsal amniotic-like cavities, ISL1^+^TFAP2A^-^ mesodermal cells are sometimes detectable (**Extended Data Fig. 4e**). Cavities at the ventral pole are lined with a single layer of FOXA2^+^SOX17^+^HNF4A^+^ ExEndo-like cells (**Fig. 1e**, **Extended Data Fig. 4f&g**). These cells also express GATA4/6 but not TFAP2A (**Extended Data Fig. 4f&g**). Thus, the ventral cavities in D8 PTED embryoids are considered hereafter as definitive yolk sac-like structures.

Notably, there are FOXA2^+^SOX17^+^ endodermal cells located in the central ‘embryonic’ region of D8 PTED embryoids (**Extended Data Fig. 4f&g**). Some of these cells do not line any cavities, nor express HNF4A, supporting their EmEndo-like fate (**Extended Data Fig. 4f&g**). The nuclear shape also differs between these endodermal cells *vs.* those in the ventral yolk sac-like structures. Specifically, FOXA2^+^SOX17^+^ endodermal cells in the ventral yolk sac-like structures show spindle-shaped nuclei, whereas those in central ‘embryonic’ region exhibit cuboid-shaped nuclei, regardless of HNF4A expression (**Extended Data Fig. 4f&g**). This observation is consistent with the nuclear morphological difference between parietal endoderm and visceral/definitive endoderm *in vivo*^42–44^ (**Fig. 1a**, **Extended Data Fig. 1**).

We next compared lineage marker expression between mesodermal cells in the ‘embryonic’ region *vs.* those surrounding yolk sac-like structures with a putative ExM-like fate (**Extended Data Fig. 5a-d**). Mesodermal cells in both regions express mesoderm markers N-CAD, FOXF1, or SNAIL (**Extended Data Fig. 5b&c**). BRA^+^ gastrulating cells, however, are only evident in the ‘embryonic’ region (**Extended Data Fig. 5b**). ISL1^+^NKX2.5^+^ and ISL1^+^HAND2^+^ cardiac mesoderm-like cells are also detected in the ‘embryonic’ region^45–47^ (**Extended Data Fig. 5c**). Mesodermal cells surrounding yolk sac-like structures are ACT2^+^E-CAD^-/LOW^, and LAMININ and DECORIN (DCN) are detected in these areas, suggesting active ECM secretion by these cells (**Extended Data Fig. 5d**). Importantly, in the putative ExM-like compartment surrounding yolk sac-like structures, CD34^+^VE-Cadherin^+^ HEP cells are evident, supporting primary hematopoiesis in these structures^20^ (**Fig. 1e**, **Fig. S4I**). We also observed periodic collapse and re-expansion of the cavity in the ventral yolk sac-like structure during PTED embryoid culture (**Supplementary Video 3**), consistent with observations from other human embryoids that develop yolk sac-like structures^9,12,20^.

To examine the developmental potential of yolk sac-like structures, they are dissected from D8 PTED embryoids and are transferred to culture plates for prolonged culture with Essential 6 (**Extended Data Fig. 5e**). On Day 14, FOXA2^+^HNF4A^+^AFP^+^APOA4^+^ ExEndo-like cells are evident, and these cells are negative or show low expression of SOX17 (**Extended Data Fig. 5e**); this observation is consistent with *in vivo* yolk sac endoderm marker expression^20,29,46,48^. HNF4A^-^AFP^-^APOA4^-^FOXA2^+^ cells are sometimes evident, albeit with a very low frequency (**Extended Data Fig. 5e**). Interestingly, colonies of PU.1^+^RUNX1^+^ cells with a rounded cell shape are sometimes evident near the yolk sac-like tissues, suggesting hematopoiesis and blood progenitor formation in these tissues (**Extended Data Fig. 5e**).

Immunofluorescence staining of D8 PTED embryoids for lineage markers associated with embryonic germ layers (ectoderm: SOX2, OCT4, and NANOG; mesoderm: BRA; endoderm: FOXA2) reveal a distinct layered organization of ectodermal, mesodermal and endodermal cells in the center ‘embryonic’ region along the *D*-*V* axis (**Fig. 2a&b**, **Extended Data Fig. 6a**). In some D8 PTED embryoids (7 out of 12 embryoids), BRA+ gastrulating-like cells appear localized on one side of the trilaminar embryonic disc-like structure, a prominent feature marking the *A*-*P* axis formation in CS8 *Cy* monkey embryos (**Fig. 2c-e**, **Extended Data Fig. 1f**, **Extended Data Fig. 6b**). Clusters of SOX2^+^OCT4^+^NANOG^-^ cells are occasionally detected in dorsal regions of the trilaminar embryonic disc-like structure, indicative of neural ectoderm development (**Extended Data Fig. 6c**).

**Figure 2.**
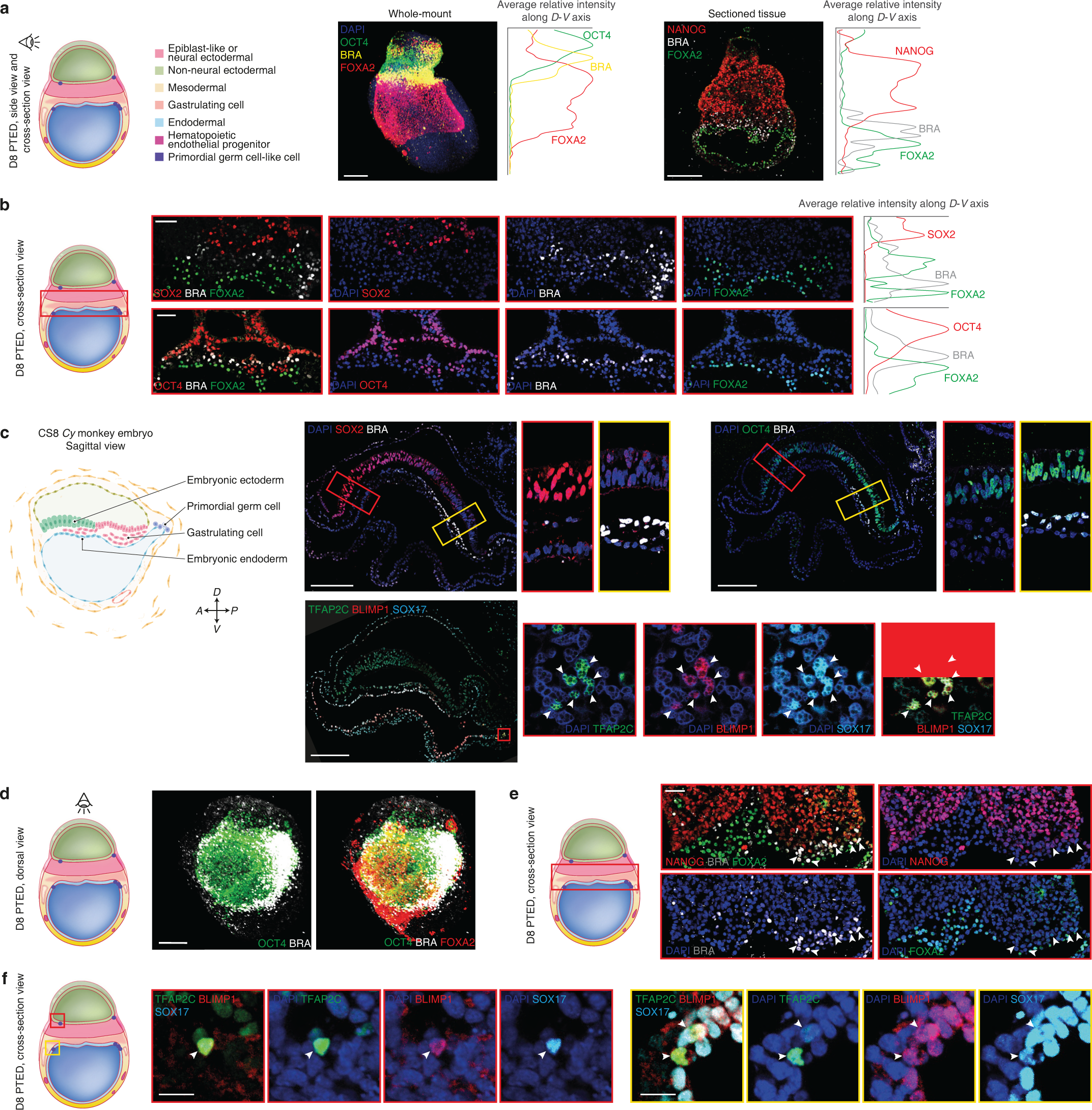
PTED embryoids with a trilaminar embryonic disc-like structure and primordial germ cell-like cells (PGCLCs). **a.** Left: Cartoon of D8 PTED embryoids. Right: Representative light-sheet micrograph of whole-mount D8 PTED embryoid and confocal micrograph of cross-sectioned D8 PTED embryoid stained for indicated lineage markers. DAPI counterstains cell nuclei. Plots show average relative intensity of indicated marker expression along the *D*-*V* axis. BRA, BRACHYURY. Scale bars, 200 µm. **b.** Left: Cartoon of D8 PTED embryoid, with its trilaminar embryonic disc-like region marked by a red box. Middle: Representative confocal micrographs showing lineage marker expression pattern in the trilaminar embryonic disc-like structure. DAPI counterstains cell nuclei. Right: Plots show average relative intensity of indicated marker expression along the *D*-*V* axis. Scale bars, 50 µm. **c.** Left: Schematic of E22 (CS8) *Cy* monkey embryo. Right: Representative confocal micrographs of E22 *Cy* monkey embryos stained for different indicated lineage markers. Magnified views are included for regions highlighted by colored boxes. DAPI counterstains cell nuclei. White arrowheads mark TFAP2C^+^BLIMP1^+^SOX17^+^ cells. Scale bars, 200 µm. **d.** Left: Cartoon of D8 PTED embryoid. Right: Representative light-sheet micrographs of whole-mount D8 PTED embryoids stained for indicated lineage markers. Scale bar, 200 µm. **e.** Left: Cartoon of D8 PTED embryoid, with its trilaminar embryonic disc-like region marked by a red box. Right: Representative confocal micrographs showing lineage marker expression pattern in the trilaminar embryonic disc-like structure. DAPI counterstains cell nuclei. White arrowheads mark BRA^+^ cells. Scale bar, 50 µm. **f.** Left: Cartoon of D8 PTED embryoid, with two regions highlighted where magnified views are provided on the right. Right: Representative confocal micrographs showing expression of lineage markers as indicated. DAPI counterstains cell nuclei. White arrowheads mark TFAP2C^+^BLIMP1^+^SOX17^+^ cells. Scale bars, 20 µm.

In primates, PGCs emerge during peri-gastrulation development (**Fig. 2c**)^48–50^. PTED embryoids also give rise to primordial germ cell-like cells (PGCLCs). On Day 3, small clusters of TFAP2C^+^SOX17^+^ PGCLCs emerge near TFAP2A^+^ NNE cells (**Extended Data Fig. 7**). In D8 PTED embryoids, some TFAP2C^+^SOX17^+^ PGCLCs co-express BLIMP1 or NANOG, suggesting a fully committed stage to germline cell development (**Fig. 2f**, **Extended Data Fig. 7d&e**), consistent with CS8 *Cy* monkey embryo data (**Fig. 2c**).

Embryoids are amenable to genetic studies^28,51^. We next sought to investigate the role of NODAL signaling, given its involvement in embryonic mesoderm and endoderm development during mammalian gastrulation^31,33^. To this end, a *NODAL*-knockout (KO) H9 hESC line^52^ was utilized. Immunostaining analysis of D3 *NODAL*-KO PTED embryoids reveals the presence of TFAP2A^+^ NNE cells, NANOG^+^ or SOX2^+^ epiblast-like cells, BRA^+^ gastrulating cells, and TFAP2C^+^SOX17^+^ PGCLCs (**Extended Data Fig. 8a&b**). However, endoderm differentiation is completely blocked in D3 or D5 *NODAL*-KO PTED embryoids, with an absence of FOXA2^+^ cells (**Extended Data Fig. 8a&c**). No cavities are present in D5 *NODAL*-KO PTED embryoids (**Extended Data Fig. 8c**). *NODAL*-KO PTED embryoids were also generated using another hESC line (ESI017), revealing consistent results (**Extended Data Fig. 8d&e**). Together, our data support the role of NODAL signaling in human peri-gastrulation development.

PTED embryoids can be generated from different hPSC lines, including both hESCs (H1 and H9 hESCs) and human induced pluripotent stem cells (hiPSCs) (**Extended Data Fig. 9**). Efficiencies of PTED embryoids from H9 hESCs and the hiPSC line are comparable, but both are less than those from H1 hESCs (**Extended Data Fig. 4c**, **Extended Data Fig. 9**). This likely is due to a greater endoderm differentiation efficiency of H1 hESCs (**Extended Data Fig. 9a-f**).

### Effects of colony size and cell seeding density in PTED embryoid development

Initial culture conditions have critical impacts on peri-gastrulation embryoid development^5,6,8^. To examine the effect of colony size, H9 hESCs are seeded onto smaller Geltrex adhesive islands with a diameter of 400 µm, under the same cell seeding density of 5,000 cells mm^-2^. On Day 3, TFAP2A^+^ NNE cells envelop the entire cell colony, whereas the number of FOXA2^+^ endodermal cells is notably reduced (**Extended Data Fig. 10a-c**). On Day 5, no cavities are present, suggesting failed PTED embryoid formation (**Extended Data Fig. 10d**).

Cell seeding density also has a profound effect. As seeding densities decrease from 5,000 cells mm^-2^ to 313 cells mm^-2^ for 800 µm diameter adhesive islands, the efficiency of PTED embryoid formation decreases significantly (**Extended Data Fig. 10e&f**). With low cell seeding densities, hESC colonies tend to remain as 2D structures, with TFAP2A^+^ NNE cells developing only at colony borders (**Extended Data Fig. 10e&f**), a feature noted in other 2D human peri-gastrulation embryoids^5,53^.

### Single-cell transcriptomic analysis

We applied single-cell RNA-sequencing (scRNA-seq) for transcriptomic analysis of D2, D5, and D8 PTED embryoids (**Fig. 3**). Unbiased clustering of cells from all three PTED embryoids reveals distinct cell populations annotated as EpiLC (epiblast-like cell), Gast (gastrulating cell), NasM / EmgM (nascent / emergent mesoderm), AdvM (advanced mesoderm), ExM, Endo (endoderm), NNE (non-neural ectoderm, including amniotic ectoderm and embryonic non-neural ectoderm), NE (neural ectoderm), PGCLC and HEP (**Fig. 3a&b**). Annotations of cell clusters for PTED embryoids are consistent with those for a CS7 human gastrula^46^. Specifically, NNE cluster exhibits heightened expression of *ISL1*, *TFAP2A*, *GATA3*, and *GABRP*, Endo shows notable expression of *SOX17* and *FOXA2*, Gast upregulates *TBXT* (gene encoding BRA) but downregulates *NANOG* and *SOX2*, cells with NE identity upregulate *SOX2* but downregulate *POU5F1* (gene encoding OCT4) and *NANOG*, PGCLC cluster exhibits heightened expression of *POU5F1*, *NANOG*, *TFAP2C* and *SOX17*, and HEP upregulates *CD34* and *CDH5* (gene encoding VE-Cadherin) (**Fig. 3b**, **Extended Data Fig. 11**, **Extended Data Fig. 12a-d**). *PAX6* is not highly expressed in the NE cluster, suggesting these cells might not yet be fully committed to the neural ectoderm fate.

**Figure 3.**
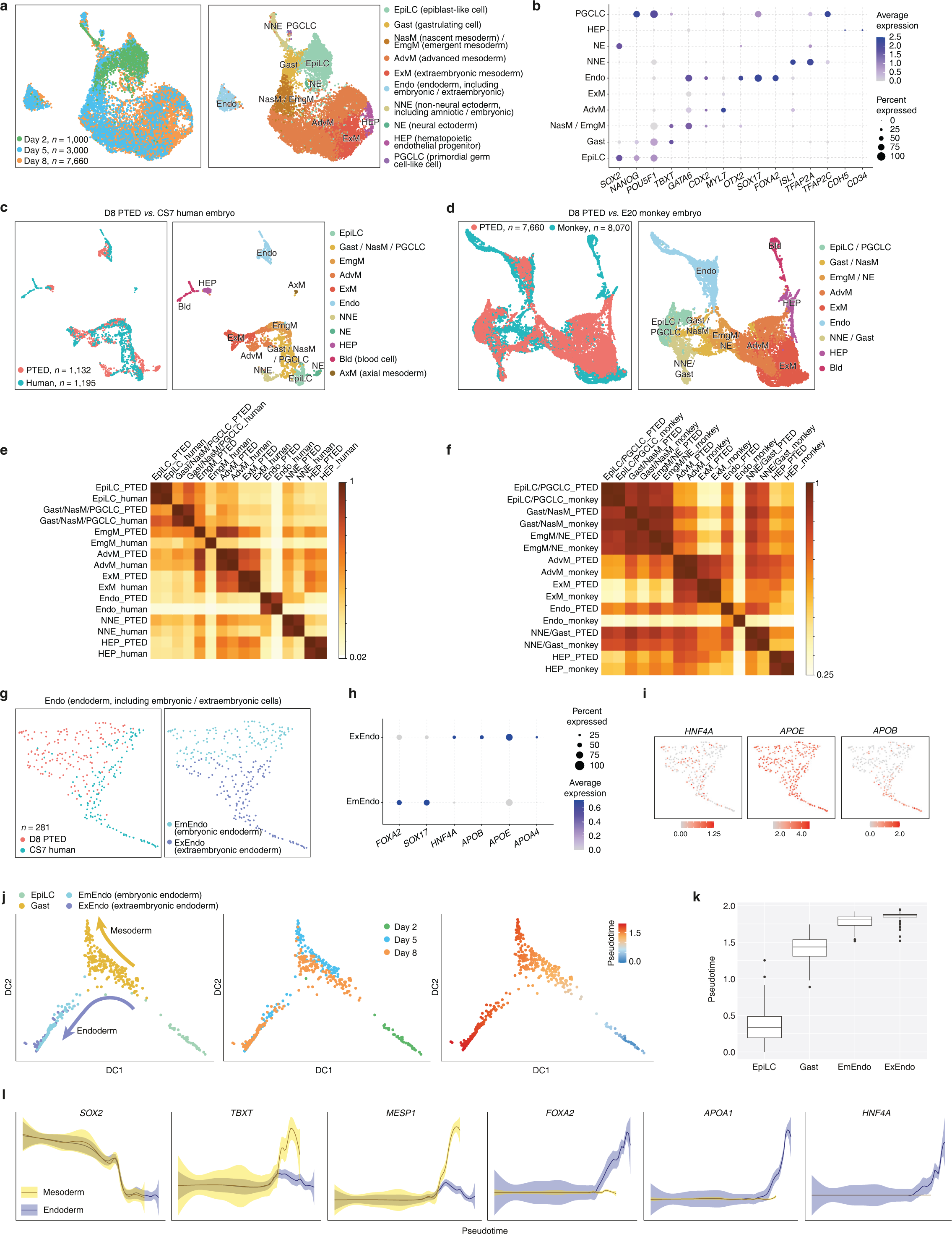
Single-cell transcriptomic analysis of PTED embryoids. **a.** Unified manifold approximation and projection (UMAP) visualization of integrated dataset combining scRNA-seq data of D2, D5, and D8 PTED embryoids, color-coded according to either culture times (*left*) or cell identity annotations (*right*). *n* indicates cell number. **b.** Dot plot showing expression of key marker genes across different cell clusters in PTED embryoids. Dot sizes represent proportions of cells expressing corresponding genes, while dot colors indicate averaged scaled values of log-transformed expression levels. **c.** UMAP visualization of integrated dataset combining scRNA-seq data of D8 PTED embryoids and CS7 human embryo^46^, color-coded according to either sample origins (*left*) or cell identity annotations (*right*). *n* indicates cell number. **d.** UMAP visualization of integrated dataset combining scRNA-seq data of D8 PTED embryoids and E20 (CS8) monkey embryo^48^, color-coded according to either sample origins (*left*) or cell identity annotations (*right*). *n* indicates cell number. **e.** Heatmap showing correlation coefficients between paired cell clusters from CS7 human embryo and D8 PTED embryoids as indicated. **f.** Heatmap showing correlation coefficients between paired cell clusters from E20 monkey embryo and D8 PTED embryoids as indicated. **g.** UMAP of scRNA-seq data from Endo cluster separated from integrated scRNA-seq dataset of D8 PTED embryoids and CS7 human embryo, revealing two subclusters annotated as embryonic endoderm (EmEndo) and extraembryonic endoderm (ExEndo). Data are color-coded according to cell origins (*left*) or cell identity annotations (*right*). *n* indicates cell number. **h.** Dot plot showing expression of key marker genes in EmEndo and ExEndo subclusters. Dot sizes represent proportions of cells expressing corresponding genes, while dot colors indicate averaged scaled values of log-transformed expression levels. **i.** Feature plots showing expression patterns of selected lineage markers in EmEndo and ExEndo subclusters. **j.** Diffusion map showing lineage relationships between EpiLC, Gast, EmEndo, and ExEndo cells. These cell clusters are separated from integrated dataset combining scRNA-seq data of D2, D5, and D8 PTED embryoids. Data are color-coded according to cell identity annotations (*left*), culture times (*middle*), or pseudotime values (*right*). Arrows indicate mesodermal and endodermal differentiation trajectories. **k.** Box plot showing pseudotime values of EpiLC, Gast, EmEndo, and ExEndo cells. Box: 25% - 75%, bar-in-box: median, and whiskers: 5% and 95%. **l.** Expression dynamics of selected genes along pseudotime during mesodermal and endodermal development. Level of confidence (0.95) is indicated by band width.

Analyzing cellular compositions of D2, D5, and D8 PTED embryoids, respectively, based on scRNA-seq data, reveals that only cells associated with EpiLC, Gast, and NNE identities are present on Day 2, whereas all other cell clusters, except NE and HEP, emerge on Day 5 (**Extended Data Fig. 11a-d**). NE and HEP clusters only appear in D8 PTED embryoids (**Extended Data Fig. 11e-g**), consistent with our immunostaining data (**Fig. 1d&e**, **Extended Data Fig. 4**, **Extended Data Fig. 5**). Between Day 2 to Day 8, proportions of mesodermal lineages, including Gast, NasM, EmgM, AdvM, and ExM cells, increases drastically (**Extended Data Fig. 12a-d**). Differential gene expression analysis (**Extended Data Fig. 12c**) reveals gene expression profiles of different cell clusters consistent with immunostaining results in **Extended Data Fig. 2-7**.

Subclustering analysis was conducted for NNE cluster, revealing two distinct subpopulations annotated as amniotic ectoderm or Amnion and embryonic non-neural ectoderm or EmNNE (**Extended Data Fig. 12e**). Amnion subcluster upregulate *ISL1*, *TFAP2A*, *GABRP*, *VTCN1*, *ACTC1*, and *TGFBI*. EmNNE subcluster highly expresses *CLDN10*, *MCM5*, *TUBG1*, *GRHL3*, and *GATA3*, consistent with the gene expression profile of embryonic surface ectoderm^46^ (**Extended Data Fig. 12f&g**).

Mesodermal lineages, including cells in Gast, NasM, EmgM, AdvM, ExM, and HEP clusters, were isolated for further analysis (**Extended Data Fig. 12h-k**). These cells show non-specific expression of lineage markers associated with paraxial or lateral plate mesoderm (**Extended Data Fig. 12i&j**). This suggests that the mesodermal clusters in D8 PTED embryoids might not yet become fully specified into specific mesodermal subtypes and thus might correspond to some transitional states, consistent with CS7 human embryo data^46^.

Nevertheless, marker expression among embryonic (Gast, NasM, EmgM, rostral advanced mesoderm (RAdvM), and caudal advanced mesoderm (CAdvM)) and extraembryonic mesodermal lineages (ExM and HEP) is generally consistent with immunostaining data in **Extended Data Fig. 5a-d**. Diffusion map and pseudotime analysis were conducted for the mesodermal lineages, suggesting a sequential lineage progression of Gast → NasM → EmgM → RAdvM / CAdvM → ExM → HEP (**Extended Data Fig. 12k**). Thus, our data hint at the possibility that ExM cells might originate from gastrulating cells in PTED embryoids.

CellChat analysis was conducted to infer cell-cell interactions in PTED embryoids (**Extended Data Fig. 13a**). Mesodermal lineages (Gast, NasM, EmgM, AdvM, ExM, and HEP clusters) are identified as main sources of BMP, WNT, non-canonical WNT, and TGF-β signals and Endo cluster as a signaling center of FGF signals (**Extended Data Fig. 13a**). Notably, VEGF and KIT ligands, which are critically involved in primary hematopoiesis^54,55^, are mainly produced from AdvM / ExM / HEP / Endo and HEP / Endo cells, respectively (**Extended Data Fig. 13a**). Heatmaps of incoming and outgoing signaling patterns further support that cells in AdvM / ExM / HEP / Endo clusters are main sources of various developmental signals (**Extended Data Fig. 13b**). Specifically, cells in ExM, HEP, and Endo clusters show greater outgoing signals associated with ECM-related pathways (**Extended Data Fig. 13b**), suggesting their active ECM production states, consistent with immunostaining results in **Extended Data Fig. 5d**. Both CellChat and heatmap analyses of incoming and outgoing signaling patterns support that cells in the EpiLC, Gast, and PGCLC clusters show heightened outgoing NODAL signals, whereas NasM/EmgM- and AdvM-associated cells are the only cells perceiving NODAL signals (**Extended Data Fig. 13a&b**), consistent with the important role of NODAL in driving mesoendoderm development *in vivo*^31,33^ and our data in **Extended Data Fig. 8**.

Developmental trajectory branches were inferred based on pseudotime analysis of integrated scRNA-seq data of D2, D5, and D8 PTED embryoids (**Extended Data Fig. 13c**). This analysis suggests that EPI cluster gives rise to three main branches, one towards NE development, another towards PGC specification, and the third one leading to NNE / Endo / HEP / ExM / AdvM development (**Extended Data Fig. 13c**). This analysis also supports that embryonic and extraembryonic mesoderm lineages in PTED embryoids share common progenitors (**Extended Data Fig. 13c**). Gene regulatory network (GRN) analysis for each cell cluster reveals their respective regulons (**Extended Data Fig. 13d**).

We next compared D8 PTED embryoid transcriptome with scRNA-seq data of the CS7 human gastrula^46^ or of an E20 or CS8 *Cy* monkey embryo^48^, respectively, revealing high concordances between them (**Fig. 3c-f**, **Extended Data Fig. 14**). Cell clusters of PTED embryoids overlap with their counterparts in the human gastrula and in the *Cy* monkey embryo, respectively (**Fig. 3c&d**, **Extended Data Fig. 14a&e**). Some notable differences are revealed as well. D8 PTED embryoid contains a NE population, not present in the human gastrula nor in the *Cy* monkey embryo (**Fig. 3c&d**, **Extended Data Fig. 14a&e**). In contrast, D8 PTED embryoid does not contain axial mesoderm (AxM), which is present in the human gastrula, nor blood cells (Bld), which exist in both the human gastrula and *Cy* monkey embryo (**Fig. 3c&d**, **Extended Data Fig. 14a&e**). Correlation analysis between paired cell clusters from the PTED embryoid and human gastrula or *Cy* monkey embryo, respectively, further supports transcriptome similarities between corresponding cell clusters, except for EmgM and Endo clusters (**Fig. 3e&f**).

The presence of both EmEndo- and ExEndo-like cells in PTED embryoids prompted us to conduct subclustering analysis for the Endo cluster identified in the integrated data of D8 PTED embryoids and CS7 human gastrula (**Fig. 3g**, **Extended Data Fig. 14b**). Two subclusters are revealed and annotated as EmEndo and ExEndo. EmEndo subcluster is characterized by enriched expression of *FOXA2* and *SOX17*, whereas ExEndo is marked by upregulated *HNF4A* and *APOA*/*B*/*E* and downregulated *SOX17* (**Fig. 3h&i**, **Extended Data Fig. 14c**). ExEndo subcluster contains cells from both PTED embryoids and human gastrula (**Fig. 3g**, **Extended Data Fig. 14b&f**). We posit that ExEndo-like cells in PTED embryoids are those cells lining yolk sac-like cavities. Pathway enrichment analysis based on highly expressed genes suggests activation of signaling pathways related to metabolic absorption and digestion in the ExEndo subcluster (**Extended Data Fig. 14d**), consistent with the nutritional supply function of the definitively yolk sac. Similar subclustering analyses conducted for the Endo cluster identified in the integrated data of D8 PTED embryoids and E20 *Cy* monkey embryo lead to similar conclusions (**Extended Data Fig. 14e-i**).

### Lineage relationship in PTED embryoid

During peri-gastrulation human development, ExEndo and ExM in the definitive yolk sac are thought to be originated from the hypoblast^20,43,44,56^. However, PTED embryoids, which are generated from primed hPSCs, do not contain any hypoblast-related lineages. Thus, we hypothesize that gastrulating-like cells in PTED embryoids might give rise to both embryonic- or extraembryonic-like lineages, with those allocated in the yolk sac-like structure being specified into ExEndo- and ExM-like cells (**Fig. 4a**). To examine this hypothesis, clusters of EpiLC, Gast, and Endo (including both EmEndo and ExEndo cells) were isolated for diffusion map analysis with pseudotime calculations (**Fig. 3j-l**). Diffusion maps support that a subpopulation of Gast cells undergo lineage diversification and differentiate into both EmEndo- and ExEndo-like cells (**Fig. 3j**). Even though these two lineages share similar trajectories, ExEndo-like cells show a greater pseudotime value than EmEndo-like cells (**Fig. 3j&k**). Pseudotime analysis reveals similar downregulation dynamics of *SOX2* for Gast and Endo cells (**Fig. 3l**). However, compared with Endo cells, Gast cells exhibit greater increases of *TBXT* and *MESP1* expression (**Fig. 3l**). Endo cells, in contrast, show higher increases of *FOXA2*, *APOA1*, and *HNF4A* (**Fig. 3l**).

**Figure 4.**
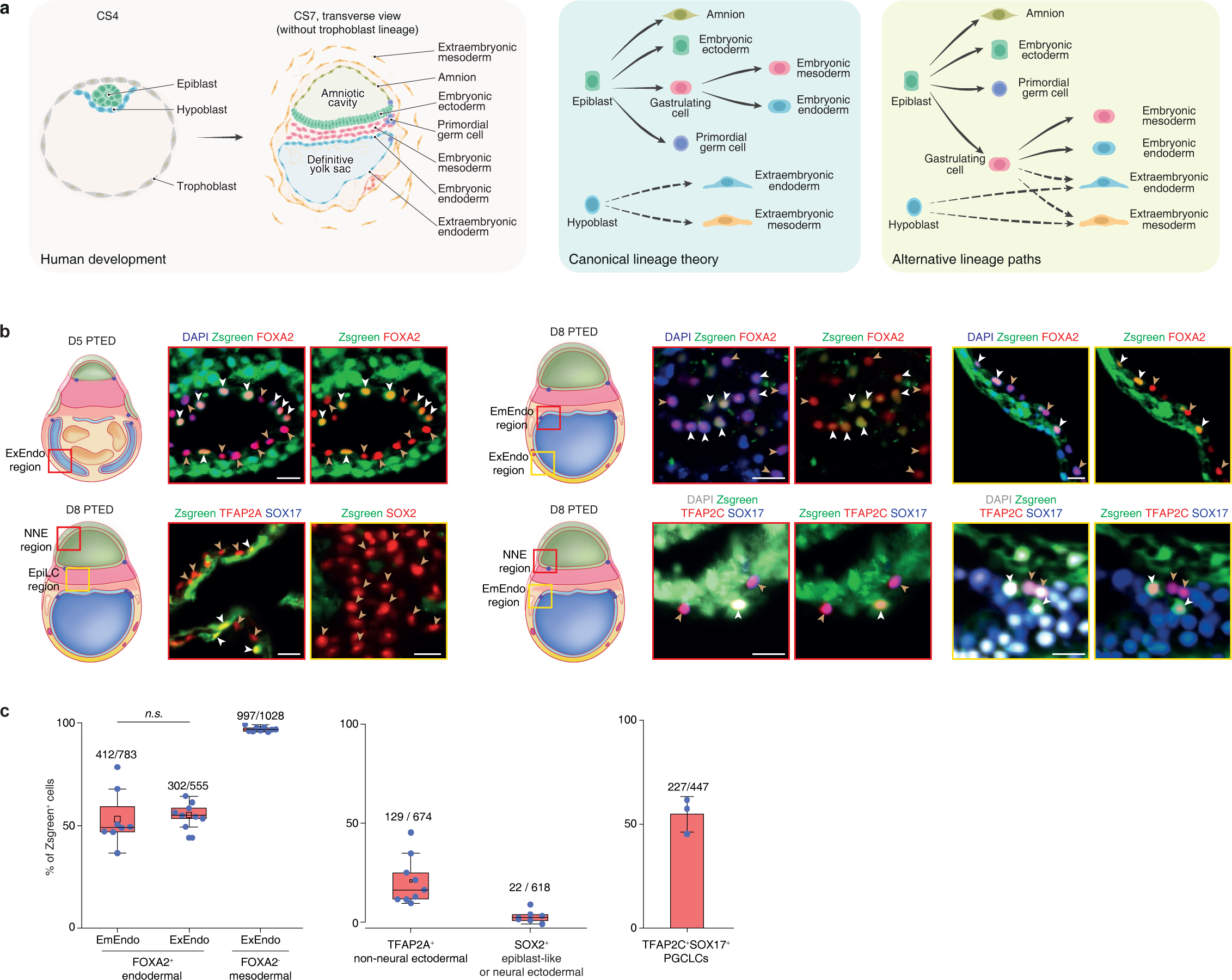
Lineage tracing in PTED embryoids. **a.** Left: Schematic showing human development from CS4 blastocyst stage to CS7 peri-gastrulation stage. Right: Schematics showing canonical lineage development theory *vs.* alternative lineage development paths, as indicated. **b.** Left: Cartoons of D5 or D8 PTED embryoids, with colored boxes marking different regions, where magnified views are provided on the right. Right: Representative confocal micrographs showing ZsGreen and lineage marker expression in different regions of PTED embryoids as indicated. PTED embryoids are generated from *TBXT*::T2A-Cre lineage tracer. DAPI counterstains cell nuclei. White and brown arrowheads mark ZsGreen^+^ and ZsGreen^-^ cells, respectively. Scale bars, 20 µm. **c.** Plots showing percentages of ZsGreen^+^ cells among different lineage populations and in different regions of D8 PTED embryoids as indicated. The numbers above the boxes and bar indicate the number of ZsGreen^+^ and the total number of cells counted in indicated populations. *n.s.*, statistically non-significant. Box: 25% - 75%, bar-in-box: median, rectangle-in-box: mean, and whiskers: 5% and 95%.

We further utilized a *TBXT*::T2A-Cre H9 hESC lineage tracer to generate PTED embryoids (**Fig. 4b&c**, **Extended Data Fig. 15**). This tracer contains a Cre-responsive ZsGreen cassette knocked into the AAVS1 locus^57^. *TBXT* plays a conserved role in mesoderm differentiation across vertebrates and is a canonical marker of gastrulation^38,58,59^. Thus, this tracer allows us to track progenies of gastrulating-like cells that upregulate *TBXT*. In both ‘embryonic’ and ‘extraembryonic’ yolk sac-like regions of D8 PTED embryoids, about 50% FOXA2^+^ endodermal cells are ZsGreen^+^ (**Fig. 4b&c**, **Extended Data Fig. 15b**), supporting the notion that gastrulating-like cells might provide common progenitors to both cell populations. Notably, about 97% FOXA2^-^ cells in yolk sac-like regions, presumptively ExM-like cells, are ZsGreen^+^, suggesting that ExM-like cells are progenies of gastrulating-like cells (**Fig. 4b&c**, **Extended Data Fig. 15b**). We further examined other cell lineages in D8 PTED embryoids. About 20% TFAP2A^+^ NNE cells are ZsGreen^+^ (**Fig. 4b&c**, **Extended Data Fig. 15c**), consistent with transient but heterogeneous expression of *TBXT* during amnion specification^38^. Only about 3% SOX2^+^ cells, presumptively associated with EpiLC or NE lineages, are ZsGreen^+^ (**Fig. 4b&c**, **Extended Data Fig. 15c**), suggesting that development of these two lineages does not involve *TBXT*. About 50% TFAP2C^+^SOX17^+^ PGCLCs are ZsGreen^+^, consistent with transient but heterogeneous expression of *TBXT* during PGCLC specification^38^ (**Fig. 4b&c**, **Extended Data Fig. 15d**). We further repeated these lineage tracing experiments with a different clone of the lineage tracer, obtaining consistent results (**Extended Data Fig. 15a-d**).

Our data suggest notable heterogeneity of *TBXT* expression during the development of different peri-gastrulation lineages. To confirm this, we applied the *TBXT*::T2A-Cre H9 hESC lineage tracer in 2D directed differentiation assays to obtain amniotic ectoderm, neural ectoderm, mesoderm, and definitive endoderm cells (**Extended Data Fig. 16**). 0.4% TFAP2A^+^ISL^+^ amniotic cells are ZsGreen^+^ (**Extended Data Fig. 16a**). No ZsGreen^+^ cells could be detected among SOX2^+^ or SOX2^+^PAX6^+^ neural ectodermal cells (**Extended Data Fig. 16b**). 94% SNAIL^+^ mesodermal cells and 21% FOXA2^+^SOX17^+^ definitive endodermal cells are ZsGreen^+^ (**Extended Data Fig. 16c&d**). The percentages of ZsGreen^+^ cells in different lineages obtained from these 2D differentiation assays are notably less than those from PTED embryoids, particularly for amnion and endodermal populations, suggesting dependence of *TBXT* expression on culture conditions and developmental environments.

### Hematopoiesis in PTED embryoid

During mammalian development, the first wave of blood cell production, or primary hematopoiesis, begins when vascular and hematopoietic cells differentiate from ExM progenitors in the yolk sac and progressively organize themselves into blood islands^20,54,55^. Our data suggest hematopoietic sites in the ExM-like compartment surrounding the yolk sac endoderm-like tissue in PTED embryoids (**Fig. 1e**, **Fig. 3a-d**, **Extended Data Fig. 5d&e**, **Extended Data Fig. 11e-g**, **Extended Data Fig. 12a-d**). CellChat analysis further shows signaling activities in the Endo population that promote HEP induction and hematopoiesis (**Extended Data Fig. 13a&b**). We thus sought to study primary hematopoiesis through PTED embryoids with enhanced endoderm differentiation. To this end, Activin A was supplemented into PTED embryoid culture between Day 0 and Day 3 (see **Methods**; **Extended Data Fig. 17a**). On Day 3, both FOXA2^+^SOX17^HIGH^HNF4A^-^ EmEndo- and FOXA2^+^SOX17^LOW^HNF4A^+^ ExEndo-like cells are evident, and they form concentric ring patterns at colony borders (**Extended Data Fig. 17b**). We conducted scRNA-seq for these D3 PTED embryoids (**Extended Data Fig. 17c-e**). Unbiased clustering reveals cell clusters annotated as EpiLC, Gast / NNE, NasM, EmgM, AdvM, EmEndo, ExEndo, and PGCLC (**Extended Data Fig. 17c-d**). Annotations of these cell clusters still follow those used for the CS7 human gastrula^46^. Thus, with enhanced endoderm differentiation, PTED embryoids give rise to consistent peri-gastrulation cell lineages, albeit with expedited differentiation and segregation of EmEndo- and ExEndo-like cells (**Extended Data Fig. 17c-d**). D3 PTED embryoids upregulate *KDR* (gene encoding VEGFR) and *GYPA* (gene encoding CD235a) (**Extended Data Fig. 17e**). Nonetheless, very few cells in D3 PTED embryoids express HEP markers *CDH5* and *CD34*^60^ (**Extended Data Fig. 17e**).

Geometric boundary confinement is necessary for ExEndo-like cell differentiation in PTED embryoids. Under the same culture condition as for Activin A-treated PTED embryoids, hPSCs seeded onto confinement-free, 2D tissue culture plates, either as single cells or as cell clusters, give rise to very few FOXA2^+^SOX17^+^HNF4A^+^ ExEndo-like cells (see **Methods**; **Extended Data Fig. 17f**).

Activin A-treated PTED embryoids start to detach from underlying glass coverslips on Day 3. Once they are free-floating, it becomes difficult to analysis hematopoiesis. To address this, we re-plated PTED embryoids on glass coverslips on Day 3 and kept them under basal E6 medium from Day 3 onwards. On Day 6, positive immunostaining for CD34 and VE-cadherin is evident, particularly in cells at colony borders, supporting HEP development (**Fig. 5a**, **Extended Data Fig. 18a&b**). To examine the roles of VEGF and KIT signaling, which are implicated in PTED embryoid development (see **Extended Data Fig. 13a&b**) and are critical for hematopoiesis^54,55,60^, Axitinib, motesanib, linifanib, or cediranib was supplemented into E6 medium between Day 3 and Day 6. Each of these inhibitors completely blocks HEP development in PTED embryoids (**Extended Data Fig. 18b**).

**Figure 5.**
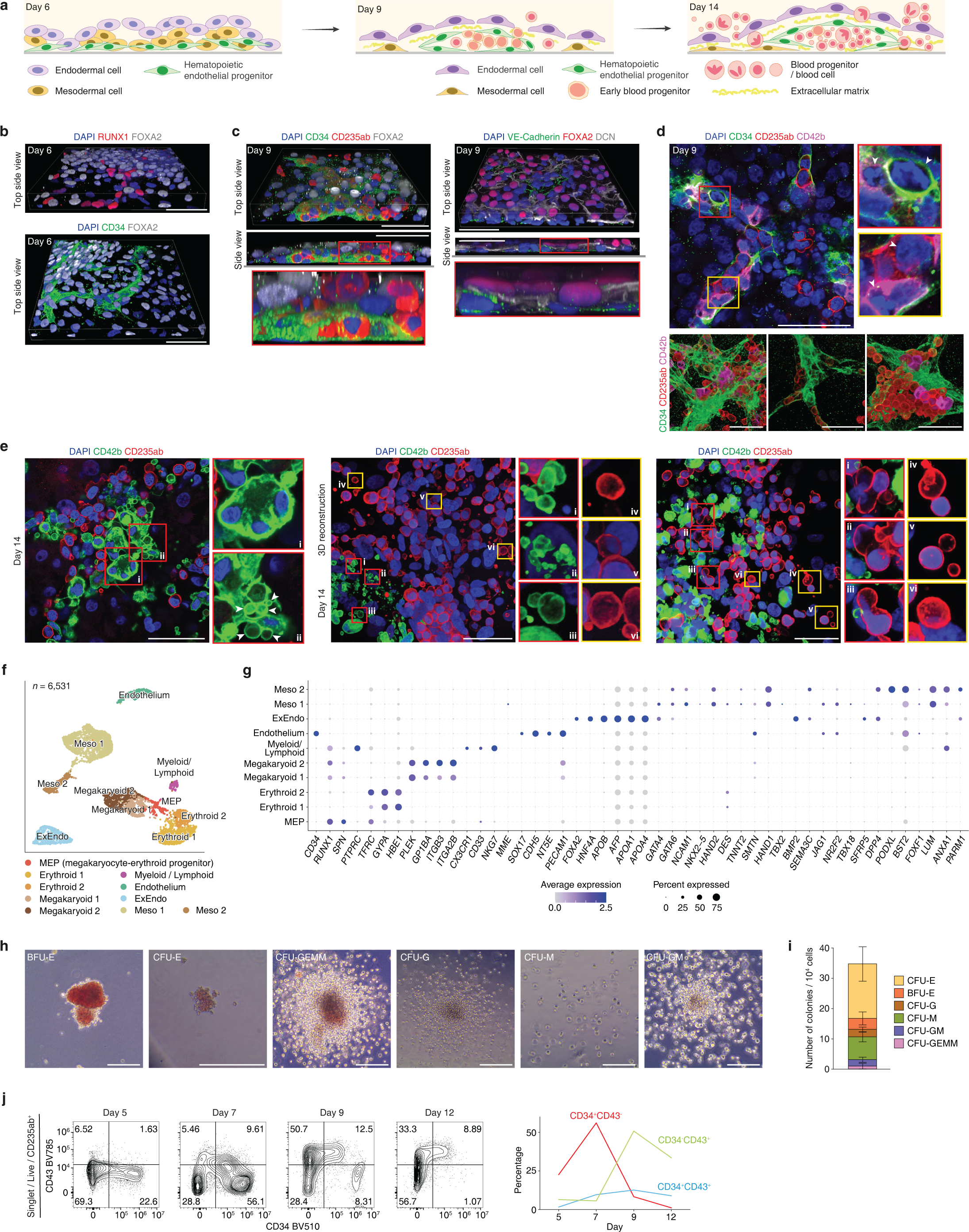
Hematopoiesis and blood cell development in PTED embryoids. **a.** Schematics showing progressive hematopoiesis and blood cell development in PTED embryoids with enhanced endoderm differentiation (see **Methods**). **b.** Representative 3D reconstruction micrographs showing D6 PTED embryoids stained for indicated lineage markers. DAPI counterstains cell nuclei. Scale bars, 50 µm. **c.** Representative 3D reconstruction micrographs showing D9 PTED embryoids stained for indicated lineage makers. DAPI counterstains cell nuclei. Zoom-in views are for boxed regions as indicated. Scale bars, 50 µm. **d.** Representative confocal micrograph (*top*) and 3D reconstruction images (*bottom*) showing D9 PTED embryoids stained for indicated lineage markers. DAPI counterstains cell nuclei. White arrowheads mark hematopoietic endothelial or blood progenitors. Scale bars, 50 µm. **e.** Representative confocal micrographs (*left*) and 3D reconstruction images (*middle* and *right*) of D14 PTED embryoids stained for indicated lineage markers. Zoom-in views are provided for boxed regions as indicated. White arrowheads mark membrane buddings of CD42b^+^ megakaryoid-like cells. DAPI counterstains cell nuclei. Scale bars, 50 µm. **f.** UMAP visualization of scRNA-seq data of D14 PTED embryoids, color-coded according to cell identity annotations. *n* indicates cell number. **g.** Dot plot showing expression of key marker genes across different cell clusters in D14 PTED embryoids. Dot sizes represent proportions of cells expressing corresponding genes, while dot colors indicate averaged scaled values of log-transformed expression levels. **h.** Representative bright-field images showing colonies generated from individual cells in D14 PTED embryoids from colony forming unit (CFU) experiments. Scale bars, 200 µm. **i.** Stacked bar plot showing cell colony numbers per 10,000 cells from D14 PTED embryoids in CFU experiments. Error bars indicate standard deviations. **j.** Left: Flow cytometry scatterplots showing CD34 and CD43 expression in CD235ab^+^ erythroid populations on different culture days of PTED embryoids. Right: Plot shows the percentages of CD34^+^CD43^+^, CD34^+^CD43^-^, and CD34^-^CD43^+^ erythroid progenitors in CD235ab^+^ erythroid populations as a function of culture days.

Spatial organization and differentiation of blood cell lineages were examined based on immunostaining of PTED embryoids. On Day 6, HEPs and blood progenitors, marked by CD34 or RUNX1, emerge beneath or among FOXA2^+^ endodermal cells (**Fig. 5b**, **Extended Data Fig. 18b**). By Day 9, tubular structures composed of CD34^+^ HEPs are evident, occasionally with PU.1^+^ blood progenitors and CD235ab^+^ erythroid cells appearing inside or right outside of these tubular structures (**Fig. 5c**, **Extended Data Fig. 18c**). Decorin (DCN), an ECM molecule, is detectable to separate FOXA2^+^ endodermal layers and hematopoietic endothelial tubes (**Fig. 5c**, **Extended Data Fig. 18c**). This spatial tissue organization is consistent with hematopoietic islands in the definitive yolk sac^15,20,61^. Such hematopoietic island-like spatial organization in PTED embryoids is maintained over time, with the hematopoietic sites continuously expanding and more blood cells forming (**Extended Data Fig. 18d**). On Day 14, some blood cell colonies without adjacent CD34^+^ HEPs are observed (**Extended Data Fig. 18d**), possibly due to HEP exhaustion or blood colony formation from re-inhabited blood progenitors.

During primary hematopoiesis, blood cells are thought to be derived from hemogenic endothelial cells^61–66^. We thus sought to examine lineage relationship between HEPs and blood progenitors. On Day 5, CD42b, a marker of megakaryocytes, begin to express alongside CD34 in HEP clusters (**Extended Data Fig. 19a**). On Day 6 and Day 7, CD235ab^+^CD34^+^ and CD42b^+^CD34^+^ cells are evident, suggesting emergence of erythroid and megakaryocyte progenitors from CD34^+^ HEPs (**Extended Data Fig. 19b&c**). Notably, CD235ab^+^CD42b^+^ cells, likely common progenitors of erythrocytes and megakaryocytes and also known as megakaryocyte-erythroid progenitors (MEPs), are observable (**Extended Data Fig. 19b**). On Day 9 and Day 14, many CD235ab^+^CD34^+^, CD42b^+^CD34^+^ and PU.1^+^CD34^+^ cells are present and are adjacent to CD34^+^ cells (**Fig. 5d**, **Extended Data Fig. 19d&e**), highlighting a sustained HEP-to-blood progenitor transition. Notably, by Day 14, many blood progenitors and blood cells could be seen free-floating and exhibit a round-shaped morphology (**Extended Data Fig. 19e**).

We further examined erythrocyte and megakaryocyte development and maturation. On Day 14, both CD235ab^+^ erythroid and CD42b^+^ megakaryoid cells show varying morphologies, suggesting these cells at different developmental stages (**Fig. 5e**, **Extended Data Fig. 20a-c**). Specifically, CD235ab^+^ erythroid cells show different sizes, and some of them are clearly undergoing enucleation, with cell nuclei polarized to one side of the cells or being squeezed out from the cells, forming bleb-like structures (**Fig. 5e**, **Extended Data Fig. 20a-c**). There are also nucleus-free CD235ab^+^ erythroid cells (**Fig. 5e**, **Extended Data Fig. 20a-c**). Similarly, CD42b^+^ megakaryoid cells exhibit a range of maturation stages, including cells with single nuclei, cells with multi-lobed nuclei, enlarged cells with multi-lobed nuclei, cells showing membrane budding, and nucleus-free platelet-like structures (**Fig. 5e**, **Extended Data Fig. 20a-c**). The observed processes of erythropoiesis and megakaryopoiesis are consistent with previous knowledge^65,67–69^.

Other mesodermal and endodermal lineages were also investigated (**Extended Data Fig. 20d-f**). Spontaneous beating of PTED embryoids was sometimes observed as early as Day 12, with beating sites increasing over time (**Extended Data Fig. 20d**, **Supplementary Video 5**). Beating cells are cTnT^+^NKX2.5^+^, indicating their cardiomyocyte identity^45–47^ (**Extended Data Fig. 20e**). Additionally, many FOXA2^+^ endodermal cells on Day 7 and nearly all of them on Day 14 show upregulated expression of HNF4A, AFP, and APOA4, suggesting continuous development of the ExEndo-like compartment in PTED embryoids^20,29,46,48^ (**Extended Data Fig. 20f**).

We conducted scRNA-seq, colony forming unit (CFU) assays, and flow cytometry to further elucidate cell identities in PTED embryoids (**Fig. 5f-j**, **Extended Data Fig. 21-23**). scRNA-seq data of D14 PTED embryoids reveals distinct hematopoiesis-related cell clusters annotated as MEP, Erythroid 1 & 2, Megakaryoid 1 & 2, Myeloid & Lymphoid, and Endothelium, each expressing lineage-specific markers, in addition to Mesoderm 1 & 2 and ExEndo clusters (**Fig. 5f&g**, **Extended Data Fig. 21a&b**, **Extended Data Fig. 22**). Some myeloid and lymphoid subtypes (granulocyte, monocyte / macrophage, B cell, NK cell, and T cell) seem to be present, albeit with limited numbers (**Fig. 5g**, **Extended Data Fig. 21b**). A portion of cells in Endothelium cluster express *NT5E*, suggesting their non-hematopoietic endothelium identity (**Extended Data Fig. 21b**). Hematopoiesis-related cell clusters were isolated for diffusion map analysis with pseudotime calculations. Developmental trajectories of erythroid, megakaryoid, and myeloid & lymphoid lineages all appear to be originated from endothelium and MEP (**Extended Data Fig. 21c**).

CFU analysis supports the existence, albeit at low frequencies, of early erythrocyte progenitors (BFU-E), late erythrocyte progenitors (CFU-E), granulocyte-erythrocyte-megakaryocyte-monocyte common progenitors (CFU-GEMM), granulocyte progenitors (CFU-G), monocyte progenitors (CFU-M), and granulocyte-monocyte common progenitors (CFU-GM) in D14 PTED embryoids (**Fig. 5h&i**), consistent with scRNA-seq data. Flow cytometric analysis shows the presence of neutrophil-like cells (CD15^+^CD31^+^), lymphoid-like progenitors (CD117^+^), monocyte- and macrophage-like cells (CD14^+^), and B cell progenitor-like cells (CD19^+^) in CD43^+^CD45^+^ populations of D9 PTED embryoids (**Extended Data Fig. 21d**). In D12 PTED embryoids, erythroid (CD235ab^+^) and megakaryoid (CD42b^+^) populations are detected, together with myeloid progenitor-like cells (CD33^+^), erythrocyte progenitors (CD235ab^+^) and megakaryocyte progenitors (CD42b^+^) within CD43^+^ populations (**Extended Data Fig. 21d**), confirming distinct blood subtype identities. Natural killer-like cells (CD56^+^CD49d^+^) and a few T cell progenitor-like cells (CD3^+^) are identified within CD45^+^CD14^-^CD19^-^ populations on Day 18 (**Extended Data Fig. 21d**). Analysis of CD34 and CD43 expression reveals dynamic changes in the percentages of CD34^+^CD43^+^, CD34^+^CD43^-^, and CD34^-^CD43^+^ cells in PTED embryoids on different days, indicating transitions from CD34^+^ to CD43^+^ progenitors and suggesting additional waves of hematopoiesis in PTED embryoids^48,61,62,70^ (**Fig. 5j**, **Extended Data Fig. 21e**).

Additional transcriptomic analysis was conducted for the ExEndo and Mesoderm 1 & 2 clusters in D14 PTED embryoids (**Extended Data Fig. 22**). Two distinct ExEndo subtypes (ExEndo 1 & 2) are revealed, both expressing markers characteristic of ExEndo, including *HNF4A*, *AFP*, *APOA1/4* and *APOB* (**Extended Data Fig. 22a&b**). Notably, ExEndo 2 shows enriched expression of ECM- and hematopoiesis-related genes (**Extended Data Fig. 22c**). Based on feature plots of expression patterns of genes highly enriched in ExEndo and hepatocytes, ExEndo-like cells in PTED embryoids show a closer resemblance to ExEndo lineage^61^ (**Extended Data Fig. 22c**). Four mesodermal subclusters were identified from subclustering analysis of Mesoderm 1 & 2 clusters: Mesoderm 1, enriched with ECM-related genes; Mesoderm 2, expressing cardiac mesoderm-related genes^45–47^; Mesoderm 3, showing mesothelium-^71^ and ExM-related gene expression; and Mesoderm 4, featured by upregulated expression of hematopoiesis-related genes (**Extended Data Fig. 22d-f**). Notably, Endoderm 2 and Mesoderm 4 upregulate some hematopoiesis-related genes but without clear expression of HEP markers (**Extended Data Fig. 17e**, **Extended Data Fig. 22c,e,f**), suggesting a nuanced hematopoietic process^60,71^.

CellChat analysis for D14 PTED embryoids reveals intricate interactions among different cell types (**Extended Data Fig. 23**). WNT ligands are primarily secreted by endothelial cells and MEPs (**Extended Data Fig. 23b&c**). Interestingly, ncWNT ligands are almost exclusively secreted by MEPs (**Extended Data Fig. 23b&c**). Both NOTCH and TGF- β pathways show complex and extensive connectivity among various cell types (**Extended Data Fig. 23b&c**). Endogenous expression of hematopoiesis-specific ligands is evident, such as erythropoietin (EPO) and stem cell factor (SCF), supporting PTED embryoids possessing an intrinsic hematopoietic property (**Extended Data Fig. 23b&c**).

## DISCUSSION

In this study we developed a transgene-free human peri-gastrulation development model, termed PTED embryoid, solely using primed hPSCs. Development of PTED embryoids, mainly through spontaneous cell differentiation and organization, bypasses blastocyst- or implantation-like developmental stages and recapitulates certain important aspects of the peri-gastrulation human development, including formation of the trilaminar embryonic layers flanked by dorsal amnion and ventral definitive yolk sac and primary hematopoiesis and blood cell generation in the definitive yolk sac. Development of PTED embryoids does not follow canonical developmental sequences of lineage diversification and tissue organization. It also does not involve the formation of a structure mimicking the primitive streak, an iconic and transient feature associated with mammalian gastrulation^32,72^. The absence of a primitive streak-like structure in PTED embryoids and their other non-canonical developmental features support the idea that PTED embryoids bypass some canonical gastrulation-related cellular events while still achieving the ultimate outcome of gastrulation: the formation of a population of embryonic and extraembryonic cells, followed by the development of a recognizable structure with organized cell lineages and spatially defined identities in an emerging coordinate system^11,73,74^. Compared to other recently reported pre- and peri-gastrulation human embryoids, whose developments are designed to reconstitute natural developmental sequences to establish *in vivo*-like cellular organization and tissue architecture^9–15^, PTED embryoids offer an ethically less challenging tool to study the self-organizing properties of human gastrulation.

Development of PTED embryoids starts from BMP4-treated primed hPSCs confined in micropatterned adhesive surfaces, without any additional extraembryonic-like cells, critically different from other recently reported pre- and peri-gastrulation human embryoids^9–14^. BMP4 treatment is effective in initiating differentiation of primed hPSCs in PTED embryoids, leading to a diverse population of peri-gastrulation-stage embryonic- and extraembryonic-like lineages. Geometric boundary confinement endowed by micropatterned adhesive surfaces provides an effective morphogenetic field to promote cellular interaction and organization of these diverse lineages during PTED embryoid development. Even without external signal inputs, the intrinsic self-organizing properties and embryonic plasticity of these peri-gastrulation-stage embryonic- and extraembryonic-like lineages engender the emergence of *in vivo*-like tissue organization and function at a global scale in the PTED embryoid.

Formation of the definitive yolk sac-like structure in PTED embryoids is unexpected. In human embryos, formation of the definitive yolk sac occurs during the gastrulation and likely involves hypoblast-derived extraembryonic lineages^20,43,44,56^, which are not present in PTED embryoids. Nonetheless, our molecular characterization and lineage tracing assays support the notion that embryonic and extraembryonic mesoderm cells, as well as embryonic and extraembryonic endoderm cells, might share common progenitors that emerge during peri-gastrulation development. This notion supports gastrulating epiblast as an alternative origin of the extraembryonic lineages giving rise to the definitive yolk sac. Importantly, there are data from studies of mouse development, albeit limited, supporting this notion^21,75^. Our data further support extrinsic controls of embryonic and extraembryonic lineage segregation and allocation during peri-gastrulation human development.

Our data show PTED embryoids as a useful experimental tool to study yolk sac hematopoiesis and blood cell formation. Unlike other blood cell generation methods^22,54,65,66,76–78^, the PTED embryoid protocol does not involve supplementation of exogenous hematopoiesis-related soluble factors. This observation supports an autonomous and self-organizing hematopoietic process in the yolk sac through intricate interactions between the extraembryonic lineages comprising the definitive yolk sac structure. Future efforts could be directed to applying the PTED embryoid to study the molecular mechanisms guiding the development of hemogenic endothelium and diversification of blood progenitors, through live imaging of lineage tracers and genetic approaches. In addition, our data showing mature erythrocytes and megakaryocytes, the presence of myeloid and lymphoid cells, and the transition from CD34^+^ to CD43^+^ progenitors, all hint at a pro-definitive hematopoietic wave in the yolk sac-like structure^61,62,70^, another area that warrants future investigations.

Together, by recapitulating aspects of both embryonic and extraembryonic development during human gastrulation, including the formation and organization of embryonic germ layers and dorsal amnion and ventral definitive yolk sac (and associated primary hematopoiesis), the transgene-free PTED embryoid provides a promising tool for studying self-organizing properties of peri-gastrulation human development, shedding light on complex cellular events involved in early human embryogenesis.

## Supporting information

Supplementary Table 1. List of differentially expressed genes, signaling pathways, regulons, and correlation coefficients from scRNA-seq analysis.

Flow cytometry gates.

## ACKNOWLEDGEMENTS

This work is supported by the Michigan-Cambridge Collaboration Initiative, the University of Michigan Mcubed Fund, and the Mid-career Biosciences Faculty Achievement Recognition Award from the University of Michigan. We acknowledge the Michigan Medicine Microscopy Core for training and support in microscopy imaging, the Michigan Orthopaedic Research Laboratories Histology Core for support in paraffin sectioning, the Michigan Advanced Genomics Core for scRNA-seq service, the Michigan Flow Cytometry Core for flow cytometry analysis, and the Michigan Lurie Nanofabrication Facility for support in microfabrication. We thank Dr. Aryeh Warmflash and his colleagues for ESI017 hESC and *NODAL*-KO ESI017 hESC lines.

## AUTHOR CONTRIBUTIONS

S.S., Y.Z., and J.F. conceived and initiated the project; S.S. designed, performed and quantified most experiments, including scRNA-seq data analysis and interpretation; Y.Z. generated *NODAL*-KO H9 hESC line; Y.S.K. generated *TBXT*::T2A-Cre lineage tracer; Y.X., J.Z., and H.W. performed monkey immunohistochemistry analysis; Z. Zhong helped with flow cytometry experiment and related data analysis; Y.S.K., N.K., X.X., Y.L., and Z. Zhou helped with data interpretation and experimental designs; S.S. and J.F. wrote manuscript; J.F. supervised study. All authors edited and approved the manuscript.

## COMPETING FINANCIAL INTERESTS

University of Michigan, Ann Arbor filed a patent application describing the PTED embryoid development protocol (63/553,448), with J.F., S.S., and Y.Z. as co-inventors. The rest of the authors declare no competing interest.

## METHODS

### Ethics statement

PTED embryoids developed in this work lack the trophectoderm lineage. As such, they are not integrated, complete models of human embryos, and they do not possess the potential to form a viable entity. All studies involving human pluripotent stem cells (hPSCs) employed in this work have obtained prior approval from the Human Pluripotent Stem Cell Research Oversight Committee at the University of Michigan, Ann Arbor. Experiments involving *Cy* monkey embryos are conducted in accordance with the Principles for the Ethical Treatment of Non-Human Primates and have received prior approval from the Institutional Animal Care and Use Committee of the Institute of Zoology, Chinese Academy of Sciences (IOZ, CAS, Appl. No: IOZ-EU-20191113). All experiments conducted in this work strictly adhere to relevant guidelines and regulations.

### Cell culture

hPSC lines used in this study include H1 human embryonic stem cell (hESC; WA01, WiCell; NIH registration number: 0043), H9 hESC (WA09, WiCell; NIH registration number: 0062), ESI017 hESC (hESC; NIH registration number: 0093) gifted from Dr. Aryeh Warmflash^34^, and the 1196a human induced pluripotent stem cell (hiPSC) line from the University of Michigan Pluripotent Stem Cell Core. All cells are maintained in a feeder-free culture system using mTeSR1 medium (STEMCELL Technologies, cat #85850). Before cell seeding, culture plates are coated with 1% lactate dehydrogenase-elevating virus (LDEV)-free, hESC cell-qualified reduced growth factor basement membrane matrix Geltrex (Thermo Fisher Scientific, cat #A1413302), derived from Engelbreth-Holm-Swarm mouse tumors. To ensure quality of hPSCs, visual examination is performed during each passage to confirm absence of spontaneously differentiated, mesenchymal-like cells in cell culture. Cells are used before reaching passage number 70. Authentication of hPSCs is conducted by both original sources and in-house. Immunostaining for pluripotency markers and successful differentiation of hPSCs to the three definitive germ layers are performed as part of the authentication process. Additionally, karyotype analysis is carried out by Cell Line Genetics to confirm karyotypic normality of hPSCs. All hPSCs are routinely tested for negative mycoplasma contamination using LookOut Mycoplasma PCR Detection Kit (Sigma-Aldrich, cat #MP0035).

### Cynomolgus monkey embryo study

*Cynomolgus* (*Cy*) monkeys (*Macaca fascicularis*) are of Southeast Asian origin. They are maintained at around 25 °C and a relative humidity of 40% - 70% on a 12 h: 12 h light-dark schedule and raised at the Xieerxin Biology Resource with the accreditation of the laboratory animal care facility in Beijing. All *Cy* monkeys are given a commercial diet twice a day with tap water ad libitum and are fed with vegetables and fruits once every day under careful veterinary supervision. Before experiments, none of the *Cy* monkeys has a clinical or experimental (drug or test) history that would affect physiological aging or increase susceptibility to disease.

Oocyte collection, intracytoplasmic sperm injection, pre-implantation embryo culture, and transfer of pre-implantation embryos to foster mothers are performed as described by Yamasaki *et al*.^79^ Briefly, female *Cy* monkeys around 6-8 years of age are chosen for oocyte collection by superovulation with follicle-stimulating hormone using an implantable and programmable microfusion device subcutaneously implanted under ultrasound detection. The day on which collected ova are artificially fertilized by sperm injection is designated as embryonic day 0 (E0). When embryos show blastocoel cavities around E6-E7, high-quality embryos are selected and transferred to appropriate recipient female *Cy* monkeys. Implanted embryos are monitored by ultrasound scanning from E14 onwards to identify successful pregnancies. Ketamine hydrochloride (10 mg kg^-1^) is administered by intramuscular injection for anesthesia of pregnant *Cy* monkey, before implanted uterus is surgically removed at E22 to obtain *Cy* monkey embryonic tissues.

### Generation of NODAL-KO hPSCs

To generate *NODAL*-knockout (KO) hPSCs, as previously reported,^52^ a 58-bp segment of genomic DNA within *NODAL* exon 1 is deleted using CRISPR/Cas9 gene editing technology. Two crRNAs are purchased from Thermo Fisher Scientific: *NODAL*_crRNA_1 (5’-AGGCUCAGCAUGUACGCCAG-3’) and *NODAL*_crRNA_2 (5’- AGACAUCAUCCGCAGCCUAC-3’). Deplexes of crRNA:tracrRNA are prepared according to a standard protocol and introduced into H9 hESCs along with Cas9 enzyme and pCXLE-EGFP expression plasmid (a gift from Dr. Shinya Yamanaka; Addgene plasmid # 27082; RRID: Addgene_27082) to enable constitutive expression of EGFP. NEON electroporation system (Thermo Fisher Scientific) is used for introducing these components into hESCs. EGFP-expressing single hESCs are isolated by fluorescence-activated cell sorting (FACS, Aria Fusion, BD Biosciences) and subsequently seeded onto 96-well plates precoated with Matrigel (Corning, cat # 354277) under mTeSR Plus medium (STEMCELL Technologies, cat #100-0276) supplemented with CloneR single-cell culture supplement (STEMCELL Technologies, cat #05888). To confirm desired deletion, genomic DNA is isolated from single cell-derived clones, and PCR is performed using primers designed for amplification of *NODAL* exon 1 (forward primer: 5’-CTTCCTTCTGCACGCCTGGTGG-3’; reverse primer: 5’-CCAACCCACAGCACTTCCCGAG-3’). Resulting amplicons are subjected to Sanger sequencing using primer 5’-CTTCCTTCTGCACGCCTGGTGG-3’.

### Microcontact printing

Polydimethylsiloxane (PDMS; Sylgard 184; Dow Corning) stamps containing circular micropatterns are fabricated using a microfabricated silicon mold. PDMS prepolymer, with a curing agent to base polymer ratio of 1:20, is poured onto the mold and baked at 110°C for 1 h. After thermal curing, PDMS stamps are peeled off the mold, before being immersed in 1% Geltrex solution (*v* / *v*, diluted in DMEM/F12) and incubated at 37°C for 1 h. Concurrently, glass coverslips (Thermo Fisher Scientific) are treated with ultraviolet ozone using an Ozone cleaner (Jelight) for 7 min. After blowing dry with nitrogen gas, Geltrex-coated PDMS stamps are placed in conformal contact with ultraviolet ozone-treated coverslips, to transfer Geltrex adhesive patterns onto coverslips.

### Generation of PTED Embryoids

On Day -2, hPSCs in tissue culture plates are dissociated using Accutase (Millipore-Sigma, cat #A6964) at 37°C for 8 min. Cells are then centrifuged and re-suspended in mTeSR1 containing 10 µM Y27632 (Tocris, cat #1254), at a concentration of 4 × 10^6^ cells mL^-1^. 140 µL of cell solution is dropped onto 12-mm diameter coverslips that are pre-printed with Geltrex adhesive islands, resulting in the cell density of about 5,000 cells mm^-2^. Coverslips holding cell solutions are placed in a 24-well plate and incubated PTED embryoids with enhanced endoderm differentiation are generated for 30 min. Coverslips are then washed gently with DMEM/F12 to remove unattached cells, before 500 µL of mTeSR1 containing Y27632 (10 µM) is added to the well plate. On Day -1, culture medium is replenished with 500 µL of mTeSR1 without Y27632. On Day 0 and Day 1, culture medium is switched to 500 µL of mTeSR1 supplemented with BMP4 (50 ng mL^-1^; R&D Systems, cat #314-BP-050). On Day 2, culture medium is switch to 500 µL of mTeSR1. From Day 3 to Day 8, half of culture medium is replaced with fresh mTeSR1 every day. Embryoids detached from coverslips between Day 5 and Day 8 are collected and transferred to a low-attachment 96-well plate, with one embryoid per well. Half of culture medium in 96-well plates is replenished with fresh mTeSR1 daily between Day 5 and Day 8.

For the prolonged culture of yolk sac-like structures, well-formed Day 8 PTED embryoids are selected and cut under a stereomicroscope. The cut yolk sac-like regions are transferred into a tissue culture-treated dish containing Essential 6™ Medium (E6, Gibco, cat #A1516401) with 1% Geltrex. The medium is switched to E6 on Day 9, and replenished every 2 days. The tissue is fixed and examined on Day 14.

### Whole-mount immunocytochemistry

For embryoids collected from Day 0 to Day 4, embryoids on coverslips are washed gently with PBS before being fixed with 4% paraformaldehyde (PFA) buffered in 1× PBS at room temperature (RT) for 1 h. After fixation, embryoids are permeabilized and blocked with a solution containing 0.3% Triton X-100 and 4% donkey serum at 4°C for 1 day. Embryoids are then incubated with primary antibody solutions at 4°C for an additional 2 days. Following incubations with primary antibodies, embryoids are then labeled with DAPI and donkey-raised secondary antibodies at 4°C for 1 day. Both primary and secondary antibodies are prepared in 4% donkey serum and 0.3% Triton X-100. After the immunostaining process, embryoids are washed with PBS and mounted with Fluoromount-G®. Immunofluorescence images are then captured using the Nikon A1si confocal laser scanning microscope (inverted).

For Day 8 PTED embryoids, embryoids are washed gently with PBS before being fixed with 4% PFA buffered in 1× PBS at RT for 1 h. After fixation, embryoids are treated with permeabilization solution as PBS containing 0.5% Triton X-100, 2% Glycine, and 20% DMSO at RT for 1 day. Then the embryoids are then blocked with PBS containing 0.5% Triton X-100, 10% DMSO, and 6% donkey serum at RT for 1 day. Embryoids are incubated with primary antibody at RT for additional 2 days. Following incubations with primary antibodies, embryoids are then labeled with DAPI and donkey-raised secondary antibodies at RT for 2 days. Both primary antibodies and secondary antibodies are prepared in PBS supplemented 0.2% Tween-20, 10 µg/mL Heparin, 5% DMSO, and 3% donkey serum. After the immunostaining process, embryoids are washed and mounted with EasyIndex (LifeCanvas Technologies, RI = 1.52). Immunofluorescence images are then captured using the ZEISS LIGHT SHEET 7 and processed using Arivis.

### Quantification of embryoid formation efficiency, length, projected area and height

Efficiency of embryoid formation is assessed using their bright-field images. On Day 5, embryoids displaying asymmetric structural features, with one pole of a greater size than the opposite pole and containing a fluid-filled cavity and the middle part showing an epithelial-like structure, are characterized as successful embryoids. On Day 8, embryoids with one pole of a greater size and displaying a cavity with a thick wall, the opposite pole with a smaller size and displaying a cavity enclosed by a monolayer of cells, and a middle part displaying a columnar epithelial-like structure, are considered as successful embryoids. The percentage of successful embryoids is calculated to determine the efficiency of embryoid generation. Bright-field images of successful embryoids are recorded to quantify their length and projected area. Note that for measurements of characteristic size features of Day 5 embryoids, gentle application of a slow flow using a pipette is employed to detach embryoids from culture surfaces.

To measure the heights, cells are seeded and embryoids are cultured on a glass-bottom dish, while keeping the other procedures the same as described before. After the process of immunostaining, a tissue clearing solution was added. The tissue clearing solution comprised 6.3 mL ddH2O, 9.2 mL OptiPrep^TM^ Density Gradient Medium (MilliporeSigma, cat #D1556), 4 g N-methyl-D-glucamine (MilliporeSigma, cat #M2004), and 5 g Diatrizoic acid (MilliporeSigma, cat #D9268). The z-stack confocal images were captured, and the heights of embryoids were measured based on the orthogonal views of DAPI stained images.

### Paraffin-embedded sectioning and immunohistochemistry

Embryoids are collected and fixed in 4% PFA at 4℃ overnight. Embryoids are then washed with PBS and mounted in a mixture of 2% low-melting agarose and 2.5% gelatin^80^. Mounted embryoids are then soaked in 70% ethanol at RT for at least 1 day. Subsequently, a tissue processing dehydration run of 4 - 6 h is performed. Mounted embryoids are then embedded in paraffin. Tissue sections with a thickness of 5 µm are then obtained using a microtome and placed on Superfrost Plus Microscope Slides (Thermo Fisher Scientific). Tissue sections are allowed to dry at RT for 2 days.

To remove paraffin, tissue sections are washed twice with xylene, followed by twice with 100% ethanol, twice with 95% ethanol, and twice with water, each for 5 min. The slides are then soaked in 1X SignalStain® Citrate Unmasking Solution and heated to 90°C for at least 30 min before being cooled to RT. After permeabilization and blocking with 0.3% Triton X-100 and 4% donkey serum at RT for 1 h, tissue sections are incubated with primary antibody solutions at RT for 2 h. Following primary antibody incubations, tissue sections are labeled with DAPI and donkey-raised secondary antibodies at RT for 1 h. Both primary and secondary antibodies are prepared in 4% donkey serum and 0.3% Triton X-100. Finally, after being washed with PBS, the slides are mounted with Fluoromount-G®. Immunofluorescence images are captured using the Nikon A1si confocal laser scanning microscope (inverted).

### Paraffin-embedded sectioning and immunohistochemistry of monkey embryos

Immediately after their retrieves from maternal uterus on E22, *Cy* monkey embryos are fixed in 4% PFA at 4°C overnight. *Cy* monkey embryos are then washed 3 times with 1× PBS for 5 min, before being dehydrated through graded alcohol (30%, 50%, 75%, 85%, 95%, and 100%) and xylene. *Cy* monkey embryos are then embedded in paraffin, with embryos oriented with their sagittal planes directly facing the dissecting microscope. Embedded *Cy* monkey embryos are serially sectioned at the thickness of 5 µm by microtome, and sectioned *Cy* monkey embryonic tissues are then mounted on glass slides.

Sectioned *Cy* monkey embryonic tissues are dewaxed and rehydrated through xylene and graded alcohol (100%, 95%, 85%, and 75%). Glass slides are then immersed in 0.01 M citric acid buffer solution (C6H8O7.H2O:C6H5Na3O7.2H2O, 1:9, pH 6.0) and heated in a microwave oven at 92 - 98°C for 15 min for antigen retrieval. After cooling to RT for 2 h, glass slides are washed once with 1× PBS for 5 min, before being incubated with 1% Triton X-100 for 30 min and blocked with 2% bovine serum albumin (BSA) for 30 min at RT. Sectioned *Cy* monkey embryonic tissues are then incubated with primary antibodies diluted in 2% BSA overnight at 4°C and washed three times with PBST for 5 min. Sectioned *Cy* monkey embryonic tissues are then incubated with secondary antibodies diluted in 2% BSA and DAPI (1 mg mL^-1^) for 1 h. Finally, after being washed three times with PBST for 5 min, the slides are mounted with anti-fade mounting medium (Gibco), before immunofluorescence images are recorded using a laser scanning confocal microscopes LSM 780 (Carl Zeiss) and LSM 880 (Carl Zeiss) and processed by Zen 7.0 (Carl Zeiss).

### Directed differentiation of amnion, neural ectoderm, mesoderm, and definitive endoderm

On Day -1, hPSCs in tissue culture plates are dissociated using Accutase at 37°C for 8 min. Cells are then centrifuged and re-suspended in mTeSR1 containing 10 µM Y27632. For amniotic cell differentiation, hPSCs are seeded into a 24-well plate, at a cell density of about 15,000 cells per well. On Day 0, culture medium is switched to E6 medium containing 50 ng mL^-1^ BMP4 and 20 ng mL^-1^ FGF2 (Thermo Fisher Scientific, cat #PHG0261). Medium is replenished daily. Samples are fixed on Day 4 for immunocytochemistry to assess amniotic cell differentiation.

For neural ectoderm differentiation, hPSCs are seeded into a 24-well plate, at a cell density of about 100,000 cells per well. On Day 0, culture medium is switched to E6 medium containing 10 µM SB431542 (STEMCELL Technologies, cat #72234) and 0.2 µM LDN193189 (STEMCELL Technologies, cat #72147). Medium is replenished daily. Samples are fixed on Day 4 for immunocytochemistry to assess neural ectoderm differentiation.

For mesoderm differentiation, hPSCs are seeded into a 24-well plate, at a cell density of about 100,000 cells per well. On Day 0, culture medium is switched to E6 medium containing 3 µM CHIR99021 (STEMCELL Technologies, cat # 100-1042), 25 ng mL^-1^ BMP4 and 50 ng mL^-1^ FGF2. From Day 1 to Day 4, culture medium is replenished daily with E6 medium containing 25 ng mL^-1^ BMP4 and 50 ng mL^-1^ FGF2. Samples are fixed on Day 4 for immunocytochemistry to assess mesodermal differentiation.

For definitive endoderm differentiation, hPSCs are seeded into a 24-well plate, at a cell density of about 300,000 cells per well. On Day 0, culture medium is switched to RPMI 1640 medium (Thermo Fisher Scientific, cat #11875093) containing 50 ng mL^-1^ BMP4 and 100 ng mL^-1^ Activin A (R&D Systems, cat #338-AC-050). On Day 1, culture medium is replenished with 500 µL of RPMI 1640 medium containing 100 ng mL^-1^ Activin A and 0.2% defined FBS (dFBS, Fisher Scientific, cat #SH3007001HI). On Day 2 and Day 3, culture medium is replenished with 500 µL of RPMI 1640 medium containing 100 ng mL^-1^ Activin A and 2% dFBS. Samples are fixed on Day 4 for immunocytochemistry to assess definitive endoderm differentiation.

### Quantification of fluorescent intensity along the radius of PTED embryoids

Quantitative analysis of fluorescent intensity is performed using Fiji ImageJ ^81^. *Z*-stack confocal images are utilized to generate a maximum intensity projection. Using the oval selection tool in ImageJ, a sample region of interest (ROI) is chosen, before “Radial Profile” function is used to plot quantitative fluorescent intensities along the radial direction. The fluorescent intensities for a specific marker are normalized by dividing by the maximum intensities, before being plotted.

### Quantification of fluorescent intensity along the dorsal-ventral axis of PTED embryoids

Quantitative analysis of fluorescent intensity is performed using MATLAB (R2022b). Images of interest, captured in different channels, are imported into MATLAB as matrices. First, the image from the DAPI channel is binarized and processed to connect all the cells using a series of morphological transformations with the “bwmorph” function. The largest connected component is then selected and fitted with an ellipse. The longer axis of this fitted ellipse is designated as the dorsal-ventral (*D*-*V*) axis.

Subsequently, the images from different channels are rotated based on the angle of the *D*-*V* axis and masked using the DAPI channel. The embryoid is divided into 80 equal sections along the *D*-*V* axis, and the average intensity in each section is calculated separately. The average intensity of each channel in each section is normalized by dividing it by the corresponding average intensity of the DAPI channel. For each channel, these normalized average intensities are further normalized by dividing them by the maximum intensity value within the channel. Finally, the normalized average intensities are smoothed and plotted. If necessary, extreme values are removed.

### Generation of PTED embryoids with enhanced endoderm differentiation

On Day -1, hPSCs in tissue culture plates are dissociated with Accutase at 37°C for 8 min to obtain single-cell suspension. Cells are then centrifuged and re-suspended in mTeSR1 containing 10 µM Y27632 at a density of 4 × 10^6^ cells mL^-1^. 140 µL of cell solution is dropped onto a 12-mm diameter coverslip pre-coated with Geltrex adhesive islands, resulting in a cell density of about 5,000 cells mm^-2^. Coverslips holding the cell solution are then placed in a 24-well plate and incubated at 37°C for 30 min. Coverslips are washed multiple times with DMEM/F12 to remove unattached cells. After washing, 500 µL of mTeSR1 containing 10 µM Y27632 is added to replenish medium. On Day 0, culture medium is switched to 500 µL of E6 medium containing 50 ng mL^-1^ BMP4 and 100 ng mL^-1^ Activin A. On both Day 1 and Day 2, culture medium is replenished with 500 µL of E6 containing 100 ng mL^-1^ Activin A. Samples are fixed on Day 3 for immunocytochemistry to assess endoderm differentiation.

For endoderm 2D cultures, hPSCs are dissociated on Day -1 using either Accutase or Dispase (STEMCELL Technologies, cat #07923) to obtain single cells or cell clusters, respectively. Singly dissociated hPSCs are re-suspended in mTeSR1 containing 10 µM Y27632, whereas hPSC clusters are suspended in mTeSR1 without Y27632. Singly dissociated hPSCs or hPSC clusters are then seeded into a 24-well plate, with a 12-mm coverslip placed in each well, at a cell density of about 300,000 cells per well. On Day 0, culture medium is switched to E6 medium containing 50 ng mL^-1^ BMP4 and 100 ng mL^-1^ Activin A. On both Day 1 and Day 2, culture medium is replenished with 500 µL of E6 medium containing 100 ng mL^-1^ Activin A. Samples are fixed on Day 3 for immunocytochemistry to assess endoderm differentiation.

For prolonged culture of PTED embryoids with enhanced endoderm differentiation, embryoids on Day 3 are detached from coverslips by mechanically removing with pipette tips. Detached cell colonies are then transferred to a new 24-well plate, with each well holding one coverslip. E6 supplemented with 1% Geltrex is added. On Day 4, culture medium is switched to E6, and it is replenished with fresh E6 daily from Day 4 to Day 14.

For the VEGF and KIT inhibitors assay, PTED embryoids with enhanced endoderm differentiation are generated as previously described. Specifically, E6 containing 50 ng mL-1 BMP4 and 100 ng mL-1 Activin A is replenished from Day 0 to Day 1, and then E6 containing 100 ng mL-1 Activin A is replenished from Day 1 to Day 3. Then VEGF and KIT inhibitors are supplemented from Day 3 to Day 6 in E6. The inhibitors used in the assay are 1.0 µM Axitinib (Selleckchem, cat #S1005), 1.0 µM Motesanib (Selleckchem, cat #S1032), 0.5 µM Linifanib (Selleckchem, cat #S1003), or 0.5 µM Cediranib (Selleckchem, cat #S1017), individually. The samples are fixed on Day 6 for hematopoiesis inhibition assessment.

### Colony forming unit (CFU) assay

Day 14 PTED embryoids with enhanced endoderm differentiation are trypsinized into single cells and suspended in IMDM medium (Thermo Fisher Scientific, Gibco™, cat # 12440053). The cells are then mixed with MethoCult medium (STEMCELL Technologies, MethoCult™ SF H4636, cat #04636) and seeded into 35 mm non-treated dishes at a density of 10,000 cells per dish, following the vendor’s instructions. The colonies were incubated at 37°C with 5% CO_2_ for 14 days before evaluation and counting.

### Flow cytometry analysis of PTED embryoids with enhanced endoderm differentiation

Flow cytometry analysis is performed on PTED embryoids with enhanced endoderm differentiation at Day 5, Day 7, Day 9, Day 12, and Day 18. The embryoids are dissociated into single cells as follows. Day 5 and Day 7 PTED embryoids are incubated with Accutase for 0.5-1 hour at 37°C until completely dissociated into single cells. Day 9, Day 12, and Day 18 PTED embryoids are incubated with a 1:1 mixture of 2.5 mg/mL Liberase TL (Millipore-Sigma, cat #5401020001) in HBSS and Dispase, supplemented with 0.1 mg/mL DNase I (Millipore-Sigma, cat #11284932001) for 0.5 hour at 37°C. The samples are then collected and centrifuged. The supernatant is aspirated, and the pellet is resuspended in Accutase and incubated for 0.5-1 hour at 37°C until the tissue is dissociated into single cells. During their incubation, the samples are pipetted up and down every 10 min to promote cell dissociation.

The resulting cell solution is strained through a 70 µm filter, pelleted by centrifugation, and resuspended in Live/Dead staining buffer (eBioscience™ Fixable Viability Dye eFluor™ 780, Invitrogen, cat #65-0865-14, 1:5000 dilution in HBSS). The cells are incubated at RT for 30 minutes, pelleted again, and washed once with Cell Staining Buffer (BioLegend, cat #420201). The cells are then resuspended in an antibody cocktail prepared in Cell Staining Buffer containing Fc Receptor Blocking Solution (BioLegend, cat #422301) and Brilliant Stain Buffer (Invitrogen, cat #00-4409-42), with a cell concentration of approximately 1 × 10^6^ cells per 100 µL. The mixture is incubated at RT for 30 minutes, pelleted by centrifugation, and resuspended in Fixation Buffer (BioLegend, cat #420801). The cells are incubated at RT for another 30 minutes, pelleted, and washed once with Cell Staining Buffer. Finally, the cells are resuspended in Cell Staining Buffer and stored at 4°C, protected from light.

Leukocytes (BioLegend, cat #426004) and CD34 PBMCs (BioLegend, cat #426901) are prepared according to the vendor’s instructions as positive controls. Unstained cells and cells stained only with live/dead dye are used as negative controls. Flow cytometry is performed using the Cytek Aurora Spectral Analyzer following standard procedures. UltraComp eBeads (Invitrogen, cat #01-2222-42) and the Arc Amine Reactive Kit (Invitrogen, cat #A10628) are used for compensation before analyzing cells. Data analysis is conducted using FlowJo 10.10.0.

### Single-cell dissociation and RNA-sequencing

PTED embryoids are washed by PBS for 2-3 times before dissociated into single cells. Day 2 and Day 5 PTED embryoids are incubated with Accutase for 0.5-1 h at 37°C until the embryoids are completely dissociated into single cells. For Day 8 PTED embryoids, they are incubated with Trypsin for 1 h at 37°C. For Day 3 PTED embryoids with enhanced endoderm differentiation, they are incubated with Accutase for 0.5 h at 37°C. For Day 14 PTED embryoids with enhanced endoderm differentiation, they are incubated with Trypsin for 50 min at 37°C. During their incubation, the samples are pipetted up and down every 10 min to promote cell dissociation. Dissociated single cells are collected into PBS containing 2% BSA before being centrifuged at 200 g for 5 min. Resultant cell pellets are resuspended in PBS containing 2% BSA. Cell filtration is performed using a 40-µm cell strainer for Day 2 and Day 5 PTED embryoids, a 70-µm cell strainer for Day 8 PTED embryoids, a 40-µm cell strainer for Day 3 PTED embryoids with enhanced endoderm differentiation, and a 70-µm cell strainer for Day 14 PTED embryoids with enhanced endoderm differentiation to obtain single-cell suspensions.

Within 1 h after cell dissociation, cells are loaded into the 10× Genomics Chromium system (10× Genomics). 10× Genomics v.3 libraries are prepared according to the manufacturer’s instructions. Libraries are then sequenced using paired-end sequencing with a minimum coverage of 20,000 raw reads per cell using Illumina NovaSeq-6000. scRNA-seq data are aligned and quantified using Cell Ranger Single-Cell Software 719 Suite (v.3.1.0, 10× Genomics) against the Homo sapiens (human) genome assembly GRCh38.p13 from ENSEMBL.

### Data integration, dimensionality reduction, and clustering

Analysis of scRNA-seq data and integration of scRNA-seq datasets are performed using Seurat (v.4.3.0) R package ^82–85^. Default settings in Seurat R package are used unless noted otherwise. Briefly, scRNA-seq data for a single cell batch is first filtered based on the total number of detected genes and percentage of mitochondrial genes. Gene expression is then normalized by the raw count divided by the total count before multiplying by 10,000 and log transformed. Top 2,000 highly variable genes are identified for each dataset using FindVariableFeatures. Cell cycle is regressed out based on cell cycle scores using CellCycleScoring during data scaling process using ScaleData. PCA analysis (RunPCA) is then performed on filtered data followed by embedding into low dimensional space with Uniform Manifold Approximation and Projection (UMAP; RunUMAP). Identification of cell clusters by a shared nearest neighbor (SNN) modularity optimization-based clustering algorithm is achieved using FindClusters.

For integration of different scRNA-seq datasets, count matrices of different datasets are filtered separately for each dataset. It should be noted that Day 2 and Day 5 datasets are downsampled, by randomly selecting 1,000 cells in Day 2 dataset and 3,000 cells in Day 5 dataset, in order to balance cell numbers in different clusters in integrated datasets. Each integrated dataset is then normalized separately, and highly variable features are selected before being integrated using IntegrateData based on 2,000 anchor features. After dataset integration, integrated scRNA-seq dataset is analyzed following standard Seurat pipeline. Annotation of cell clusters is based on expression of canonical lineage marker genes.

Dot plots and feature plots are generated using DotPlot and FeaturePlot in Seurat, respectively. Differentially expressed genes (DEGs) for each cluster are identified using FindAllMarkers, with a minimal fold difference of 0.25 in the logarithmic scale between the cluster of interest and all other clusters. Heatmaps are plotted based on top 15 highly differentially expressed genes using plot_heatmap in Scillus package.

### Analysis of cell-cell interactions

R package CellChat v.1.6.1 is used to perform cell-cell communication analysis ^86^. Briefly, based on manually curated databases that consider known structural compositions of ligand-receptor interactions, CellChat infers and analyzes intercellular communication networks from scRNA-seq data using network analysis and pattern recognition. Seurat object including count matrix and clustering results is imported to CellChat. A default human database is used for analysis, and only secreted signaling pathways from Kyoto Encyclopedia of Genes and Genomes (KEGG) are used. Default values are used for all parameters.

### Diffusion map and pseudotime prediction

Diffusion maps are obtained using the R-package Density^87^, which computes kernel density estimates with parametric starts and asymmetric kernels. To generate diffusion maps, PCA embeddings of the Seurat object are used as input to the “diffmap” function. Diffusion pseudotime values are calculated using the “DPT” function. The roots of the diffusion map are manually initiated based on conventional developmental knowledge and then automatically updated and chosen by the algorithm. Expression levels of selected genes are fitted as a function of the pseudotime with “loess” method with the “geom_smooth” function. Cells with extreme pseudotime values are counted as outliner and excluded from the diffusion map plot and gene expression dynamic analysis.

### Trajectory branches interference using pseudotime

Developmental trajectories are inferred for embryoid development in a form of a branching tree using R package URD v1.1.1 ^88,89^. The diffusion map is computed using R package Destiny ^87^ by invoking calcDM function from URD on normalized counts from embryoid datasets (sigma.use = “local”). Root cells are assigned manually. Specifically, for analysis of integrated Day 2 / Day 5 / Day 8 PTED embryoids dataset, EpiLC cluster from Day 2 embryoids is assigned as root cells. Simulating diffusion from the root to each cell, cells are ordered in pseudotime by floodPseudotime (*n* = 30, minimum.cells.flooded = 5) and floodPseudotimeProcess functions. Terminal states of the tree (“tips”) are manually selected based on *in vivo* knowledge. To reveal developmental trajectories from scRNA-seq data, 10,000 biased random walks per tip are simulated using simulateRandomWalksFromTips. Walks are then processed into visitation frequencies using RandomWalksFromTips. Finally, branching trees are established using buildTree. Following parameters are used for building the branching tree in this work: visit.threshold = 0.7, minimum.visits = 10, bins.per.pseudotime.window = 5, cells.per.pseudotime.bin = 50, divergence.method = “preference”, and p.thresh = 0.05.

### Gene regulatory network analysis

Regulatory activity of transcription factors associated with specific cell types is assessed using the R-package SCENIC (Single Cell Regulatory Network Inference and Clustering, v.1.1.2-01) and Python package Arboreto ^90^. Briefly, filtered counts of the integrated Seurat object are used as inputs of SCENIC. GRNBoost2 in Arboreto is used to infer co-expression modules between transcription factors and candidate target genes. Each co-expression module is then analyzed using cis-regulatory motif analyses (RcisTarget). Only modules with significant motif enrichment of the correct up-stream regulator are retained. The human motif collection v9 and the cisTarget databases for hg38 are used in the pipeline (resources.aertslab.org/cistarget/databases/old/homo_sapiens/hg19/refseq_r45/mc9nr/gene_based/). Filtered counts of the integrated Seurat object are used as input of SCENIC. All default parameters are used in SCENIC unless noted otherwise. Normalized intensity of regulons with top AUC are plotted in the heatmap.

### Integration and re-analysis of published human embryo data

The scRNA-seq dataset of the CS7 human gastrula ^46^ is integrated with Day 8 PTED embryoid scRNA-seq dataset. Since human gastrula scRNA-seq dataset is generated using Smart-seq2, to compare with Day 8 embryoid scRNA-seq dataset, which is generated using the 10× Genomics Chromium system, raw counts of each cell in the human gastrula are normalized to exon sizes before being utilized to create the Seurat object (input count = raw count × 1,000 / exon size). Exon size information is obtained from GRCh38.p13, ENSEMBL. To re-analysis of the scRNA-seq dataset of the human gastrula, relevant cell types are processed with the default Seurat pipeline, including data normalization, highly variable feature selection, scaling (including cell cycle regression), and processed by PCA and UMAP. To integrate scRNA-seq data from the human gastrula with those of embryoids, scRNA-seq datasets from Day 8 PTED embryoids are filtered, normalized, scaled (including cell cycle regression), processed by PCA and UMAP, before cluster analysis is performed. Downsampling of Day 8 embryoid dataset is performed using Subset function, which randomly selects 150 cells from every cluster in the original datasets to form a new Seurat object. This step helps prevent larger datasets from dominating downstream analysis. Thereafter, Day 8 embryoid dataset and human embryo dataset are integrated using IntegrateData function. After integration, the integrated scRNA-seq dataset is analyzed following the standard Seurat pipeline. Annotation of cell clusters is based on expression of canonical lineage marker genes. To calculate correlation matrix, average expression of the top 2,000 highly variable genes is calculated for each cluster (grouped by cell lineage identification and cell origin) using AverageExpression function. The Pearson’s correlation coefficients are then obtained using cor function.

### Integration and re-analysis of published cynomolgus monkey embryo data

The scRNA-seq dataset of CS8 E20 *cynomolgus* (*Cy*) monkey (*Macaca fascicularis*) embryo ^48^ is integrated with Day 8 embryoid scRNA-seq dataset. To integrate datasets from different species, gene names from each dataset are matched according to orthologs between human and macaque ensembl genes (http://useast.ensembl.org/biomart/martview). Datasets of embryoid and *Cy* monkey are then processed with the default Seurat pipeline, including data normalization, highly variable feature selection, scaling (including cell cycle regression), processed by PCA and UMAP, before their integration using IntegrateData function. Integrated scRNA-seq dataset is then analyzed following the standard Seurat pipeline. Annotations of cell clusters are based on expression of canonical lineage marker genes. To calculate correlation matrix, average expression of the top 2,000 highly variable genes is calculated for each cluster (grouped by cell lineage identification and cell origin) using AverageExpression function. The Pearson’s correlation coefficients are then obtained using cor function.

### Sub-clustering analysis

The clusters of interested are selected using subset function. Then the selected dataset is processed with scaling (including cell cycle regression), PCA, UMAP, and clustering identification, as described in the default Seurat pipeline. Annotation of cell clusters was based on expression of canonical lineage marker genes.

### Pathway enrichment analysis

Pathway enrichment analysis is conducted using clusterProfiler v.4.6.2 R package ^91,92^. Specifically, positive DEGs are selected for extraembryonic endoderm (ExEndo) clusters using FindAllMarkers in Seurat package. Enriched pathways are then calculated based on DEGs using gseKEGG function.

### Statistical analysis

Statistical analysis is conducted using independent, two-tailed Student’s *t*-test in Excel (Microsoft). *p* < 0.05 is considered statistically significant.

## Code availability

R and Python scripts used in this work are available from the corresponding author upon request.

## Data availability

Data supporting findings of this study are available within the article and its Supplementary Information files and from the corresponding author upon request. scRNA-seq data supporting this study’s results are deposited at Gene Expression Omnibus with accession number GSE239707. All Source Data for graphs included in the paper are available in the online version of the manuscript.

## EXTENDED DATA FIGURE LEGENDS

**Extended Data Figure 1. Histocytochemistry analysis of human and monkey embryos.**

**a.** Schematics of human embryo development (*top*) and corresponding H&E staining of human embryonic tissues (*bottom*) at Carnegie stages (CS) 5c, 6, and 7, with different cell lineages indicted (https://www.ehd.org/virtual-human-embryo/). *D*, dorsal. *V*, ventral. Scale bars, 100 µm.
**b.** Schematic of E22 (CS8) *Cy* monkey embryo, with different cell lineages indicted. *A*, anterior. *P*, posterior.
**c-f.** Representative confocal micrographs of E22 (CS8) *Cy* monkey embryos stained for different lineage markers as indicated, showing marker expression for amnion (**c**), embryonic and extraembryonic endoderm (**d**), hematopoietic sites in definitive yolk sac (**e**), and embryonic disk (**f**). Zoom-in views are provided for boxed regions as indicated. White arrowheads mark GATA3^+^, ISL1^+^, or TFAP2A^+^ cells in **c**, SOX17^+^ or HNF4A^+^ cells in **d**, and CD34^+^ cells in **e**. Brown arrowheads in **d** mark HNF4A^-^ cells. Scale bars, 200 µm.

**Extended Data Figure 2. Development of PTED embryoids between Day 0 and Day 4.**

**a.** Top: Protocol for generating PTED embryoids. Middle and bottom: Cartoons (*middle*) and bright-field images (*bottom*) showing PTED embryoids on different days. Different colored regions in cartoons mark distinct cellular compartments as indicated. Scale bar, 200 µm.
**b.** Phase-contrast images showing regular arrays of D0 and D4 PTED embryoids. Insets show zoom-in views of single PTED embryoids. Scale bars, 1,000 µm. See **Supplementary Video 2** for live imaging of PTED embryoid development from D0 to D5.
**c.** Box plot showing PTED embryoid heights from D0 to D4. Box: 25% - 75%, bar-in-box: median, rectangle-in-box: mean, and whiskers: 5% and 95%.
**d-i.** Representative confocal micrographs showing orthogonal views from *x*-*y*, *x*-*z* and / or *y*-*z* planes of PTED embryoids on D0 (**d**), D1 (**e**), D2 (**f&g**), D3 (**h**) and D4 (**i**). PTED embryoids are stained for indicated lineage markers. Different *x*-*y* views of PTED embryoids are provided at different *z*-focal planes as indicated. The *x*-*z* and *y*-*z* views are expanded threefold along *z*-axis for visualization. Zoom-in views in **f** highlight “gastrulation-like” sites marked by emergences of BRA^+^ cell clusters. Zoom-in views in **g** mark gastrulating cells, labeled by white arrowheads, stained positive for different markers. Zoom-in view in **h** shows FOXA2^+^SOX17^+^ endodermal cells marked by white arrowheads. Plots show normalized maximum *z*-projection intensities of indicated lineage markers along colony radius of PTED embryoids on different culture days.Scale bars, 200 µm.

**Extended Data Figure 3. Characterization of D5 PTED embryoids.**

**a.** Phase-contrast image showing a regular array of D5 PTED embryoids (*left*) and bright-field image showing a single D5 PTED embryoid (*right*). White arrowhead marks a PTED embryoid detached from underlying glass coverslip. Scale bars, 1,000 µm (*left*) and 200 µm (*right*).
**b.** Left: Cartoon showing D5 PTED embryoid structure, with different colored regions marking distinct cellular compartments as indicated. Right: Representative light-sheet micrographs of whole-mount D5 PTED embryoids stained for indicated lineage markers. Scale bars, 200 µm.
**c-e.** Representative confocal micrographs showing sectioned D5 PTED embryoids stained for different lineage markers as indicated. Zoom-in views are provided for boxed regions as indicated. White arrowheads in **d** mark ISL1^+^TFAP2A^+^FOXA2^-^ or ISL1^+^SNAIL^+^HAND2^+^ cells. White arrowheads in **e** mark FOXA2^+^SOX17^+^HNF4A^-^ or FOXA2^+^HNF4A^-^TFAP2A^-^ cells. Scale bars, 200 µm.

**Extended Data Figure 4. Characterization of D8 PTED embryoids.**

**a.** Cartoon showing D8 PTED embryoid, with different colored regions marking distinct cellular compartments as indicated.
**b.** Photo showing D8 PTED embryoids collected in tissue culture plate. Scale bar, 1 mm.
**c.** Left: Box plot of PTED embryoid length along the *D*-*V* axis on D5 and D8. Box: 25% - 75%, bar-in-box: median, rectangle-in-box: mean, and whiskers: 5% and 95%. Middle: Stacked bar plot showing projected areas of different compartments of PTED embryoids on D5 and D8 as indicated. Error bars represent standard deviations. Right: Box plot showing efficiency of PTED embryoid formation on D5 and D8. Box: 25% - 75%, bar-in-box: median, rectangle-in-box: mean, and whiskers: 5% and 95%. See **Methods** for criteria used to define successful PTED embryoid formation.
**d.** Left: Representative light-sheet micrograph of whole-mount D8 PTED embryoid. Right: Confocal micrographs of sectioned D8 PTED embryoids stained for indicated lineage markers. Plots show average relative intensity of indicated marker expression along the *D*-*V* axis. Scale bars, 200 µm.
**e.** Representative confocal micrographs of sectioned D8 PTED embryoids stained for indicated lineage markers. Zoom-in views are provided for boxed regions as indicated. White arrowheads mark TFAP2A^+^GATA3^+^FOXA2^-^ or TFAP2A^+^ISL1^+^FOXA2^-^ cells. Scale bars, 200 µm.
**f.** Representative confocal micrographs of sectioned D8 PTED embryoids stained for indicated lineage markers. Zoom-in views of boxed regions are included as indicated. White arrowheads mark FOXA2^+^HNF4A^-^TFAP2A^-^ or FOXA2^+^SOX17^+^HNF4A^-^ cells. Brown arrowheads mark FOXA2^+^ HNF4A^+^TFAP2A^-^, FOXA2^+^SOX17^+^HNF4A^+^ or FOXA2^+^GATA4^+^GATA6^+^ cells. Scale bars, 200 µm.
**g.** Left: Cartoon of D8 PTED embryoid, with embryonic and extraembryonic regions marked by red boxes. Right: Box plot of aspect ratio of endodermal cell nuclei in embryonic and extraembryonic regions. Box: 25% - 75%, bar-in-box: median, rectangle-in-box: mean, and whiskers: 5% and 95%. *n.s.*, statistically non-significant. ****, *p*-value < 0.0001.

**Extended Data Figure 5. Characterization of D8 PTED embryoids and prolonged culture of yolk sac-like compartments.**

**a.** Cartoon of D8 PTED embryoid, with different colored regions marking distinct cellular compartments as indicated.
**b.** Representative confocal micrographs of sectioned D8 PTED embryoids stained for indicated lineage markers. Zoom-in views are provided for boxed regions as indicated. White arrowheads mark BRA^+^N-CAD^+^FOXF1^-^ cells, brown arrowheads label BRA^-^N-CAD^+^FOXF1^-^ cells, and light blue arrowheads mark BRA^-^N-CAD^+^FOXF1^+^ cells. Scale bars, 200 µm.
**c.** Representative confocal micrographs of sectioned D8 PTED embryoids stained for indicated lineage markers. Zoom-in views of boxed regions are provided as indicated. White arrowheads mark ISL1^+^NKX2.5^+^FOXF1^-^ or HAND2^+^ISL1^+^SNAIL^+^ cells, while brown arrowheads mark ISL1^-^NKX2.5^-^FOXF1^+^ or HAND2^-^ISL1^-^SNAIL^+^ cells, as indicated. Scale bars, 200 µm.
**d.** Representative confocal micrographs of sectioned D8 PTED embryoids stained for indicated lineage markers. Zoom-in views of boxed regions are provided as indicated. White arrowheads mark VE-CAD^+^CD34^+^FOXA2^-^ cells. Scale bars, 200 µm.
**e.** Top: Schematic showing prolonged culture of yolk sac-like tissues dissected from D8 PTED embryoids. Representative confocal micrographs showing yolk sac-like tissues on D14 stained for lineage markers associated with blood progenitors and extraembryonic and embryonic endoderm as indicated. White arrowheads mark AFP^+^HNF4A^+^FOXA2^+^, AFP^+^APOA4^+^FOXA2^+^, AFP^+^SOX17^-^FOXA2^+^ cells or PU.1^+^RUNX1^+^FOXA2^-^ cells. Brown arrowheads mark AFP^-^HNF4A^-^FOXA2^+^ or APOA4^-^FOXA2^+^ cells. Scale bars, 20 µm.

**Extended Data Figure 6. Characterization of trilaminar embryonic disc-like structures in D8 PTED embryoids.**

Representative confocal micrographs of D8 PTED embryoids stained for different lineage markers as indicated. Zoom-in views of boxed regions are provided as indicated. In **c**, white arrowheads mark OCT4^+^NANOG^-^SOX2^+^ cells, whereas brown arrowheads label OCT4^+^NANOG^+^SOX2^-^ cells. Scale bars, 200 µm.

**Extended Data Figure 7. Development of human primordial germ cell-like cells in PTED embryoids.**

**a-b.** Representative confocal micrographs showing orthogonal views from *x*-*y*, *x*-*z* and *y*-*z* planes of PTED embryoids on D3 (**a**) and D4 (**b**). PTED embryoids were stained for TFAP2C, TFAP2A, and SOX17, as indicated. Different *x*-*y* views of PTED embryoids are provided at different *z*-focal planes as indicated. The *x*-*z* and *y*-*z* views are expanded threefold along *z*-axis for visualization. Zoom-in views are provided for boxed regions as indicated. White arrowheads mark TFAP2C^+^SOX17^+^ human primordial germ cell-like cells (hPGCLCs). Scale bars, 200 µm.
**c.** Representative confocal micrographs showing D5 PTED embryoids stained for TFAP2C, TFAP2A, and SOX17. Zoom-in views are provided for color-coded boxed regions as indicated. White arrowheads mark TFAP2C^+^SOX17^+^ hPGCLCs. Scale bar, 200 µm.
**d.** Representative confocal micrographs showing D8 PTED embryoids stained for PGC markers as indicated. Zoom-in views are provided for color-coded boxed regions as indicated. White arrowheads mark TFAP2C^+^BLIMP1^+^SOX17^+^, TFAP2C^+^NANOG^+^SOX17^+^, or TFAP2C^+^SOX17^+^TFAP2A^+^ hPGCLCs. Scale bars, 200 µm.
**e.** Box plot showing percentages of cells positive for specific markers among TFAP2C^+^SOX17^+^ cells in D8 PTED embryoids. Box: 25% - 75%, bar-in-box: median, rectangle-in-box: mean, and whiskers: 5% and 95%.

**Extended Data Figure 8. PTED embryoids generated from *NODAL*-KO hPSCs.**

**a.** Left: Cartoon of D3 *NODAL*-KO PTED embryoids, with different colored regions marking distinct cellular compartments as indicated. Right: Representative confocal micrographs showing orthogonal views from *x*-*y*, *x*-*z* and *y*-*z* planes of D3 *NODAL*-KO PTED embryoids stained for indicated lineage markers. Different *x*-*y* views of PTED embryoids are provided at different *z*-focal planes as indicated. The *x*-*z* and *y*-*z* views are magnified threefold along the *z*-axis. Scale bars, 200 µm. Plots on the right show normalized maximum *z*-projection intensities of indicated lineage markers along colony radius of D3 *NODAL*-KO PTED embryoids.
**b.** Representative confocal micrographs showing D3 *NODAL*-KO PTED embryoids stained for indicated lineage markers. Different *x*-*y* views of PTED embryoids are provided at different *z*-focal planes as indicated. The *x*-*z* and *y*-*z* views are expanded threefold along the *z*-axis for visualization. Zoom-in views are provided for boxed regions as indicated. White arrowheads mark TFAP2C^+^SOX17^+^ hPGCLCs. Scale bar, 200 µm.
**c.** Left: Cartoon of D5 *NODAL*-KO PTED embryoids, with different colored regions marking distinct cellular compartments as indicated. Right: Representative confocal micrographs showing orthogonal views from *x*-*y*, *x*-*z* and *y*-*z* planes of D5 *NODAL*-KO PTED embryoids stained for indicated lineage markers. Different *x*-*y* views of PTED embryoids are provided at different *z*-focal planes as indicated. The *x*-*z* and *y*-*z* views are expanded threefold along the *z*-axis for visualization. Scale bar, 200 µm. Plot on the right shows normalized maximum *z*-projection intensities of indicated lineage markers along colony radius of D5 *NODAL*-KO PTED embryoids.
**d-e.** Representative confocal micrographs showing orthogonal views from *x*-*y*, *x*-*z* and *y*-*z* planes of D3 PTED embryoids generated from wild-type (WT; **d**) and *NODAL*-KO (**e**) ESI017 hESCs stained for indicated lineage markers. Different *x*-*y* views of PTED embryoids are provided at different *z*-focal planes as indicated. The *x*-*z* and *y*-*z* views are magnified threefold along the *z*-axis. White arrowheads mark TFAP2C^+^SOX17^+^ hPGCLCs. Scale bars, 200 µm.

**Extended Data Figure 9. PTED embryoids generated from H1 hESCs and another hiPSC line.**

**a.** Box plot of lengths of D5 and D8 H1-PTED embryoids along the *D*-*V* axis. Box: 25% - 75%, bar-in-box: median, rectangle-in-box: mean, and whiskers: 5% and 95%.
**b.** Stacked bar plot showing projected areas of different compartments of D5 and D8 H1-PTED embryoids. Error bars indicate standard deviations.
**c.** Box plot of H1-PTED embryoid formation efficiency on D2, D5, and D8. Box: 25% - 75%, bar-in-box: median, rectangle-in-box: mean, and whiskers: 5% and 95%. See **Methods** for criteria used to define successful PTED embryoid formation.
**d.** Left: Representative light-sheet micrograph of wholemount D8 H1-PTED embryoids stained for indicated lineage markers. Right: Plot showing average relative intensity of marker expression along the *D*-*V* axis. Scale bar, 200 µm.
**e.** Confocal micrographs of sectioned D5 H1-PTED embryoids stained for indicated lineage markers. Scale bars, 200 µm.
**f.** Confocal micrographs of sectioned D8 H1-PTED embryoids stained for indicated lineage markers. Zoom-in views are provided for boxed regions as indicated. White arrowheads mark VE-CAD^+^CD34^+^ cells. Plot shows average relative intensity of marker expression along the *D*-*V* axis. Scale bars, 200 µm.
**g.** Box plot of lengths of D5 and D8 hiPSC-PTED embryoids along the *D*-*V* axis. Box: 25% - 75%, bar-in-box: median, rectangle-in-box: mean, and whiskers: 5% and 95%.
**h.** Stacked bar plot showing projected areas of different compartments of D5 and D8 hiPSC-PTED embryoids as indicated. Error bars indicate standard deviations.
**i.** Box plot of hiPSC-PTED embryoid formation efficiency on D2, D5, and D8. Box: 25% - 75%, bar-in-box: median, rectangle-in-box: mean, and whiskers: 5% and 95%.
**j.** Left: Representative light-sheet micrograph of wholemount D8 hiPSC-PTED embryoid stained for indicated lineage markers. Right: Plot showing average relative intensity of marker expression along the *D*-*V* axis. Scale bar, 200 µm.
**k.** Confocal micrographs of sectioned D8 hiPSC-PTED embryoids stained for indicated lineage markers. Zoom-in views are provided for boxed regions as indicated. Plot shows average relative intensity of marker expression along the *D*-*V* axis. Scale bars, 200 µm.

**Extended Data Figure 10. Effects of colony size and cell seeding density on PTED embryoid development.**

**a.** Phase-contrast images showing regular arrays of D0 PTED embryoids with initial cell colony diameters of 400 µm (*left*) and 800 µm (*right*), respectively. Scale bars, 500 µm.
**b.** Left: Cartoon of D3 PTED embryoids generated from initial hPSC colonies with a diameter of 400 µm, with different colored regions marking different cellular compartments as indicated. Right: Representative confocal micrographs showing orthogonal views from *x*-*y*, *x*-*z* and *y*-*z* planes of D3 PTED embryoids stained for indicated lineage markers. Different *x*-*y* views of PTED embryoids are provided at different *z*-focal planes as indicated. The *x*-*z* and *y*-*z* views are expanded threefold along the *z*-axis for visualization. Scale bars, 100 µm.
**c.** Plots showing normalized maximum *z*-projection intensities of indicated lineage markers along colony radius of D3 PTED embryoids generated from hPSC colonies of different sizes as indicated.
**d.** Left: Cartoon of D5 PTED embryoids generated from initial hPSC colonies with a diameter of 400 µm, with different colored regions marking different cellular compartments as indicated. Right: Representative confocal micrographs showing orthogonal views from *x*-*y*, *x*-*z* and *y*-*z* planes of D5 PTED embryoids stained for indicated lineage markers. Different *x*-*y* views of PTED embryoids are provided at different *z*-focal planes as indicated. The *x*-*z* and *y*-*z* views are expanded threefold along the *z*-axis for visualization. Scale bars, 100 µm.
**e.** Left: Cartoon of D2 PTED embryoids generated from initial hPSC colonies with a diameter of 800 µm under different cell seeding density conditions as indicated. Different colored regions in cartoons mark different cellular compartments as indicated. Right: Representative confocal micrographs showing orthogonal views from *x*-*y*, *x*-*z* and *y*-*z* planes of D2 PTED embryoids stained for indicated lineage markers. Different *x*-*y* views of PTED embryoids are provided at different *z*-focal planes as indicated. The *x*-*z* and *y*-*z* views are expanded threefold along the *z*-axis for visualization. Scale bars, 200 µm. Plots show normalized maximum *z*-projection intensities of indicated lineage markers along colony radius of D2 PTED embryoids.
**f.** Box plot showing percentages of D2 PTED embryoids with a top center amnion (NNE) region as a function of initial cell seeding density. Box: 25% - 75%, bar-in-box: median, rectangle-in-box: mean, and whiskers: 5% and 95%. ***, *p* < 0.001.

**Extended Data Figure 11. Single-cell transcriptomic analysis of D2, D5, and D8 PTED embryoids, respectively.**

**a.** UMAP visualization of scRNA-seq data of D2 PTED embryoids, color-coded according to cell identity annotations. *n* indicates cell number.
**b.** Feature plots showing expression patterns of selected lineage markers used for cell identity annotations in UMAP plot of D2 PTED embryoid.
**c.** UMAP visualization of scRNA-seq data of D5 PTED embryoids, color-coded according to cell identity annotations. *n* indicates cell number.
**d.** Feature plots showing expression patterns of selected lineage markers used for cell identity annotations in UMAP plot of D5 PTED embryoid.
**e.** UMAP visualization of scRNA-seq data of D8 PTED embryoids, color-coded according to cell identity annotations. *n* indicates cell number.
**f.** Dot plot showing expression of key marker genes across different cell clusters in D8 PTED embryoids. Dot sizes represent proportions of cells expressing corresponding genes, while dot colors indicate averaged scaled values of log-transformed expression levels.
**g.** Feature plots showing expression patterns of selected lineage markers used for cell identity annotations in UMAP plot of D8 PTED embryoid.

EpiLC, epiblast-like cell; Gast, gastrulating cell; NasM, nascent mesoderm; EmgM, emergent mesoderm; AdvM, advanced mesoderm; ExM, extraembryonic mesoderm; Endo, endoderm; NNE, non-neural ectoderm; NE, neural ectoderm; HEP, hematopoietic endothelial progenitor; PGCLC, primordial germ cell-like cell.

**Extended Data Figure 12. Single-cell transcriptome analysis of integrated dataset combining scRNA-seq data of D2, D5, and D8 PTED embryoids.**

**a.** UMAP visualization of integrated scRNA-seq dataset of D2, D5, and D8 PTED embryoids, separated by culture day from integrated UMAP plot in **Fig. 3a**. Data are color-coded according to cell identity annotations. *n* indicates cell numbers.
**b.** Stacked bar plot showing cellular compositions in D2, D5, and D8 PTED embryoids as indicated.
**c.** Heatmap showing expression levels of differentially expressed genes identified from integrated scRNA-seq dataset of D2, D5, and D8 PTED embryoids. Color bars above the heatmap indicate cell identities and culture days. Selected genes are highlighted as indicated.
**d.** Feature plots showing expression patterns of selected lineage markers used for cell identity annotations in integrated UMAP plot in **Fig. 3a**.
**e.** Subclustering analysis of scRNA-seq data of NNE cluster separated from integrated dataset of D2, D5, and D8 PTED embryoids, revealing two subclusters annotated as Amnion and EmNNE (embryonic non-neural ectoderm). UMAP plots are color-coded according to culture times (*left*) or cell subcluster identity annotations (*right*). *n* indicates cell number.
**f.** Dot plot showing expression of key marker genes in EmNNE and Amnion subclusters. Dot sizes represent proportions of cells expressing corresponding genes, while dot colors indicate averaged scaled values of log-transformed expression levels.
**g.** Feature plots showing expression patterns of selected lineage markers in UMAP plots of EmNNE and Amnion subclusters.
**h.** Subclustering analysis of scRNA-seq data of mesodermal clusters separated from integrated dataset of D2, D5, and D8 PTED embryoids. UMAP plots are color-coded according to culture times (*left*) or cell subcluster identity annotations (*right*). *n* indicates cell number.
**i.** Dot plot showing expression of key marker genes in mesodermal subclusters. Dot sizes represent proportions of cells expressing corresponding genes, while dot colors indicate averaged scaled values of log-transformed expression levels.
**j.** Feature plots showing expression patterns of selected lineage markers in UMAP plots of mesodermal clusters.
**k.** Diffusion map of cells in mesodermal clusters. Data are color-coded according to cell identity annotations (*top*) or pseudotime values (*bottom*) as indicated.

EpiLC, epiblast-like cell; Gast, gastrulating cell; NasM, nascent mesoderm; EmgM, emergent mesoderm; AdvM, advanced mesoderm; ExM, extraembryonic mesoderm; Endo, endoderm; NNE, non-neural ectoderm; NE, neural ectoderm; HEP, hematopoietic endothelial progenitor; PGCLC, primordial germ cell-like cell; EmNNE, embryonic non-neural ectoderm; RAdvM, rostral advanced mesoderm; CAdvM, caudal advanced mesoderm.

**Extended Data Figure 13. Analyses of cell lineage interactions, signaling pathways, lineage branching, and gene regulatory networks using integrated dataset combining scRNA-seq data of D2, D5, and D8 PTED embryoids.**

**a.** Circle plots depicting incoming and outgoing signaling pathways across different cell clusters in integrated scRNA-seq dataset of D2, D5, and D8 PTED embryoids.
**b.** Heatmap showing incoming and outgoing signaling pathways across different cell clusters in integrated scRNA-seq dataset of D2, D5, and D8 PTED embryoids. Selected signaling pathways are highlighted as indicated.
**c.** Lineage branching based on pseudotime analysis of integrated scRNA-seq dataset of D2, D5, and D8 PTED embryoids.
**d.** Gene regulatory network (GRN) analysis of different cell clusters in integrated scRNA-seq dataset of D2, D5, and D8 PTED embryoids. Selected regulons are highlighted as indicated.

EpiLC, epiblast-like cell; Gast, gastrulating cell; NasM, nascent mesoderm; EmgM, emergent mesoderm; AdvM, advanced mesoderm; ExM, extraembryonic mesoderm; Endo, endoderm; NNE, non-neural ectoderm; NE, neural ectoderm; HEP, hematopoietic endothelial progenitor; PGCLC, primordial germ cell-like cell.

**Extended Data Figure 14. Transcriptomic comparisons between D8 PTED embryoids and CS7 human gastrula and between D8 PTED embryoids and E20 *Cy* monkey embryo, respectively.**

**a.** UMAP showing scRNA-seq data of D8 PTED embryoids (*left*) and CS7 human gastrula (*right*), separated from integrated dataset of D8 PTED embryoids and CS7 human gastrula. Data are color-coded according to cell identity annotations. *n* indicates cell numbers.
**b.** UMAP of integrated scRNA-seq data of endodermal cells (Endo) separated from integrated scRNA-seq dataset of D8 PTED embryoids and CS7 human embryo, revealing two subclusters annotated as embryonic endoderm (EmEndo) and extraembryonic endoderm (ExEndo). Data are color-coded according to cell origins (*left*) or cell identity annotations (*right*) as indicated. *n* indicates cell number.
**c.** Feature plots showing expression patterns of selected lineage markers in EmEndo and ExEndo subclusters, which are separated from integrated scRNA-seq dataset of D8 PTED embryoids and CS7 human embryo.
**d.** Pathway enrichment analysis of ExEndo subcluster identified from integrated scRNA-seq dataset of D8 PTED embryoids and CS7 human embryo.
**e.** UMAP of scRNA-seq data of D8 PTED embryoids (*left*) and E20 *Cy* monkey embryo (*right*), separated of integrated scRNA-seq dataset of D8 PTED embryoids and E20 *Cy* monkey embryo. Data are color-coded according to cell identity annotations. *n* indicates cell numbers.
**f.** UMAP of integrated scRNA-seq data of endodermal cells (Endo) isolated from integrated scRNA-seq dataset of D8 PTED embryoids and E20 *Cy* monkey embryo, revealing three subclusters annotated as EmEndo, ExEndo1, and ExEndo2. Data are color-coded according to cell origins (*left*) or cell identity annotations (*right*) as indicated. *n* indicates cell numbers.
**g.** Dot plot showing expression levels of key marker genes in EmEndo, ExEndo1, and ExEndo2 subclusters, which are separated from integrated scRNA-seq dataset of D8 PTED embryoids and E20 *Cy* monkey embryo. Dot sizes represent proportions of cells expressing corresponding genes, while dot colors indicate averaged scaled values of log-transformed expression levels.
**h.** Feature plots showing expression patterns of selected lineage markers in EmEndo, ExEndo1, and ExEndo2 subclusters, which are separated from integrated scRNA-seq dataset of D8 PTED embryoids and E20 *Cy* monkey embryo.
**i.** Pathway enrichment analysis of ExEndo subclusters (ExEndo1 and ExEndo2) identified from integrated scRNA-seq dataset of D8 PTED embryoids and E20 *Cy* monkey embryo.

EpiLC, epiblast-like cell; Gast, gastrulating cell; NasM, nascent mesoderm; EmgM, emergent mesoderm; AdvM, advanced mesoderm; ExM, extraembryonic mesoderm; Endo, endoderm; NNE, non-neural ectoderm; NE, neural ectoderm; HEP, hematopoietic endothelial progenitor; PGCLC, primordial germ cell-like cell; Bld, blood cell; AxM, axial mesoderm; EmEndo, embryonic endoderm; ExEndo, extraembryonic endoderm.

**Extended Data Figure 15. Lineage tracing in PTED embryoids.**

**a-d.** Top: Cartoons of D5 or D8 PTED embryoids as indicated, with colored boxes marking different regions, where magnified views are provided at the bottom. Bottom: Representative confocal micrographs showing ZsGreen and lineage marker expression in different regions of PTED embryoids as indicated. PTED embryoids are generated from two different clones of the *TBXT*::T2A-Cre lineage tracer as indicated. DAPI counterstains cell nuclei. White and brown arrowheads mark ZsGreen^+^ and ZsGreen^-^ cells, respectively. Scale bars, 20 µm.

**Extended Data Figure 16. Directed differentiation of hPSCs towards different lineages using 2D cultures.**

Left: Schematics of 2D directed differentiation protocols of hPSCs to generate amniotic cells (**a**), neural ectoderm cells (**b**), mesodermal cells (**c**), and definitive endoderm cells (**d**). Middle: Representative confocal micrographs showing ZsGreen and lineage marker expression in cell cultures. Right: Box plots showing percentages of cells positive for indicated lineage markers (*left*) and percentages of ZsGreen^+^ cells among indicated lineage populations (*right*). In **c**, white arrowheads mark ZsGreen^+^SNAIL^+^ cells, while brown arrowheads label ZsGreen^-^SNAIL^+^ cells. In **d**, white arrowheads mark ZsGreen^+^FOXA2^+^SOX17^+^ cells, while brown arrowheads label ZsGreen^-^FOXA2^+^SOX17^+^ cells. DAPI counterstains cell nuclei. Box: 25% - 75%, bar-in-box: median, rectangle-in-box: mean, and whiskers: 5% and 95%. Scale bars, 20 µm.

**Extended Data Figure 17. Development of PTED embryoids with enhanced endoderm differentiation.**

**a.** Schematic showing development of PTED embryoids with enhanced endoderm differentiation (see **Methods**). Y27, Y2732. ExEndo, extraembryonic endoderm. EmEndo, embryonic endoderm.
**b.** Representative confocal micrographs showing D3 PTED embryoids stained for indicated endodermal lineage markers. Zoom-in views are provided for boxed regions as indicated. White dashed line marks the boundary between FOXA2^+^SOX17^HIGH^HNF4A^-^ EmEndo-like cells and FOXA2^+^SOX17^LOW^HNF4A^+^ ExEndo-like cells. DAPI counterstains cell nuclei. Scale bar, 200 µm.
**c.** UMAP of scRNA-seq data of D3 PTED embryoids with enhanced endoderm differentiation. Data are color-coded according to cell identity annotations. *n* indicates cell number.
**d.** Dot plot showing expression of key marker genes across different cell clusters identified in D3 PTED embryoids. Dot sizes represent proportions of cells expressing corresponding genes, while dot colors indicate averaged scaled values of log-transformed expression levels.
**e.** Feature plots showing expression patterns of selected hematopoiesis-related markers in UMAP plots.
**f.** Schematics (*left*) and representative confocal micrographs (*right*) showing endoderm differentiation under 2D culture conditions, with hPSCs seeded as either single cells (*top*) or cell clusters (*bottom*). Cells were stained for indicated endodermal lineage markers on D3. DAPI counterstains cell nuclei. Scale bars, 100 µm.

**Extended Data Figure 18. Hematopoiesis and blood cell development in PTED embryoids.**

**a.** Schematic showing progressive hematopoiesis and blood cell development in PTED embryoids with enhanced endoderm differentiation (see **Methods**).
**b.** Left: Representative confocal micrographs of D6 PTED embryoids stained for indicated lineage markers. Zoom-in views are provided for boxed regions as indicated. Scale bar, 200 µm. Middle: Representative confocal micrographs showing PTED embryoids treated with indicated VEGF/KIT inhibitors on D6 stained for different lineage markers. Scale bars, 200 µm. Right: Representative confocal micrographs showing orthogonal views from *x*-*y*, *x*-*z* and *y*-*z* planes of D6 PTED embryoids, stained for indicated lineage markers. Scale bars, 50 µm. DAPI counterstains cell nuclei.
**c.** Representative confocal micrographs showing orthogonal views from *x*-*y*, *x*-*z* and *y*-*z* planes and 3D reconstruction micrographs of D9 PTED embryoids, stained for indicated lineage markers. DAPI counterstains cell nuclei. Zoom-in views are provided for boxed regions as indicated. Scale bars, 50 µm.
**d.** Representative confocal micrographs showing orthogonal views from *x*-*y*, *x*-*z* or *y*-*z* planes and 3D reconstruction images of D14 PTED embryoids, stained for indicated lineage markers. DAPI counterstains cell nuclei.

**Extended Data Figure 19. Blood cell development in PTED embryoids.**

**a.** Representative confocal micrographs showing D5 PTED embryoids stained for indicated lineage markers. White arrowhead marks CD34^+^CD42b^+^ cells. DAPI counterstains cell nuclei. Scale bar, 50 µm.
**b.** Representative confocal micrographs showing D6 PTED embryoids stained for indicated lineage markers. White arrowheads mark CD235ab^+^CD42b^+^ cells, brown arrowhead marks CD235ab^+^CD34^+^ cells, and blue arrowhead marks CD34^+^CD235ab^-^ CD42b^-^ cells. DAPI counterstains cell nuclei. Scale bars, 50 µm.
**c.** Representative confocal micrographs showing D7 PTED embryoids stained for indicated lineage markers. DAPI counterstains cell nuclei. Scale bar, 50 µm.
**d.** Representative confocal micrographs showing D9 PTED embryoids stained for indicated lineage markers. Zoom-in views are provided for boxed regions as indicated. White arrowheads mark CD34^+^CD42b^+^ cells, brown arrowheads label CD235ab^+^CD34^+^ cells, and blue arrowheads mark CD34^+^CD235ab^-^CD42b^-^ cells. DAPI counterstains cell nuclei. Scale bars, 50 µm.
**e.** Representative phase-contrast and confocal micrographs showing D14 PTED embryoids stained for indicated lineage markers. Zoom-in views are provided for boxed regions as indicated. White arrowheads mark CD34^+^PU.1^-^CD42b^-^ cells, brown arrowheads label CD34^+^ blood progenitors that are also CD235ab^+^, CD42b^+^, or PU.1^+^, and blue arrowheads mark blood cells that have detached from CD34^+^ hematopoietic endothelial progenitors. DAPI counterstains cell nuclei. Scale bars, 200 µm (for the first row of images) or 50 µm.

**Extended Data Figure 20. Blood cell, mesodermal and endodermal lineage developments in PTED embryoids.**

**a.** Schematic showing blood cell development from hematopoietic endothelial progenitors.
**b.** Representative phase-contrast and confocal micrographs showing erythroid- and megakaryoid-like cells in D14 PTED embryoids stained for indicated lineage markers. DAPI counterstains cell nuclei. Scale bars, 200 µm.
**c.** Representative phase-contrast, confocal, and 3D reconstruction images showing CD235ab^+^ erythroid- and CD42b^+^ megakaryoid-like cells in D14 PTED embryoids. Zoom-in views are provided for boxed regions as indicated. White arrowheads mark RUNX1^+^CD235ab^+^ or RUNX1^+^CD42b^+^ cells, while brown arrowheads mark RUNX1^-^ CD235ab^+^ or RUNX1^-^CD42b^+^ cells. Scale bars, 50 µm.
**d.** Representative bright-field micrographs showing spontaneous beating cardiac tissues in PTED embryoids on different culture days. White arrowheads mark spontaneous beating sites. See **Supplementary Video 5** for spontaneous beating tissues in PTED embryoid. Scale bars, 100 µm.
**e.** Representative confocal micrographs and 3D reconstruction image showing D14 PTED embryoids stained for different lineage markers associated with myocardium tissues. Scale bars, 20 µm.
**f.** Representative confocal micrographs showing D7 and D14 PTED embryoids stained for indicated extraembryonic endoderm markers. Scale bars, 20 µm.

**Extended Data Figure 21. Single-cell transcriptomic and flow cytometry analysis of hematopoiesis in PTED embryoids.**

**a.** Subclustering analysis of scRNA-seq data of hematopoiesis-related cell clusters (Endothelium, MEP, Erythroid 1, Erythroid 2, Megakaryoid 1, Megakaryoid 2, and Myeloid/Lymphoid) separated from D14 PTED embryoids with enhanced endoderm differentiation. UMAP plot is color-coded according to cell subcluster identity annotations. *n* indicates cell number.
**b.** Feature plots showing expression patterns of selected lineage markers in UMAP plots of hematopoiesis-related cell clusters.
**c.** Diffusion map of cells in hematopoiesis-related cell clusters. Data are color-coded according to cell identity annotations (*top*) or pseudotime (*bottom*) as indicated.
**d.** Representative flow cytometric analysis of CD15^+^CD31^+^ neutrophil-like cells, CD14^+^ monocyte- or macrophage-like cells, CD117^+^ lymphoid-like progenitors, and CD19^+^ B cell progenitor-like cells in the CD43^+^CD45^+^ population of D9 PTED embryoid cultures; CD235ab^+^ erythroid-like and CD42b^+^ megakaryoid-like populations in D12 PTED cultures; CD33^+^ myeloid-like cells, CD235ab^+^ erythrocyte progenitors-like cells, and CD42b^+^ megakaryocyte progenitors-like cells in the CD43^+^ population of D12 PTED cultures; CD56^+^CD49d^+^ natural killer-like cells and CD3^+^ T cell progenitor-like cells in the CD45^+^CD14^-^CD19^-^ populations of D18 PTED cultures.
**e.** Left: Representative flow cytometric analysis of CD34 and CD43 expression on cells in the entire PTED culture (*top*) or in the CD235ab^+^ erythroid-like population (*bottom*) on different culture days as indicated. Right: Plots showing percentages of CD34^+^CD43^+^, CD34^+^CD43^-^, and CD34^-^CD43^+^ cells among the entire PTED cell population (*top*) or in the CD235ab^+^ erythroid-like population (*bottom*) on different culture days.

**Extended Data Figure 22. Single-cell transcriptomic analysis of endodermal and mesodermal clusters in PTED embryoids with enhanced endoderm differentiation.**

**a.** Subclustering analysis of scRNA-seq data of ExEndo cluster separated from D14 PTED embryoids with enhanced endoderm differentiation. UMAP plot is color-coded according to cell subcluster identity annotations. ExEndo, extraembryonic endoderm. *n* indicates cell number.
**b.** Feature plots showing expression patterns of selected endodermal lineage markers in UMAP plots of endodermal clusters.
**c.** Feature plots showing expression patterns of selected lineage markers, which are highly expressed in ExEndo or hepatocytes, or associated with extracellular matrix (ECM) or different hematopoietic lineages, as indicated.
**d.** Subclustering analysis of scRNA-seq data of mesodermal clusters (Meso 1 and Meso 2) separated from D14 PTED embryoids with enhanced endoderm differentiation. UMAP plot is color-coded according to cell subcluster identity annotations. *n* indicates cell number.
**e.** Dot plot showing expression of key marker genes in different mesodermal subclusters. Dot sizes represent proportions of cells expressing corresponding genes, while dot colors indicate averaged scaled values of log-transformed expression levels.
**f.** Feature plots showing expression patterns of selected mesodermal lineage markers in UMAP plots of mesodermal clusters.

**Extended Data Figure 23. Analysis of cell lineage interactions and signaling pathways using scRNA-seq data of D14 PTED embryoids with enhanced endoderm differentiation.**

**a.** UMAP visualization of scRNA-seq data of D14 PTED embryoids, with endodermal and mesodermal subcluster information incorporated. UMAP plot is color-coded according to cell cluster identity annotations. *n* indicates cell number.
**b.** Circle plots depicting incoming and outgoing signaling pathways across different cell clusters in scRNA-seq data of D14 PTED embryoids.
**c.** Heatmap showing incoming and outgoing signaling pathways across different cell clusters in scRNA-seq data of D14 PTED embryoids.

## SUPPLEMENTARY VIDEOS

**Supplementary Video 1. *Z*-stack immunostaining images showing PTED embryoid development from Day 0 to Day 4.** This video shows confocal images with different *z* focuses for PTED embryoids from Day 0 to Day 4 stained for indicated markers. Scale bar, 200 µm.

**Supplementary Video 2. Live imaging of PTED embryoid development from Day 0 to Day 5.** Time-lapse imaging captures dynamic development of PTED embryoids during their early developmental stages. Scale bar, 500 µm.

**Supplementary Video 3. Live imaging of PTED embryoids generated from the *TBXT*::T2A-Cre lineage tracer from Day 5.5 to Day 7.5.** Time-lapse imaging captures dynamic development of PTED embryoids generated from the *TBXT*::T2A-Cre lineage tracer. Scale bar, 500 µm.

**Supplementary Video 4. Light-sheet imaging of whole-mount Day 8 PTED embryoids immunostained for different lineage markers.** This video shows light-sheet imaging of whole-mount D8 PTED embryoids, revealing structural organization and marker expression patterns in them. Scale bar, 200 µm.

**Supplementary Video 5. Video showing spontaneous beating tissues in PTED embryoids with enhanced endoderm differentiation.** This video captures spontaneous beating cells in PTED embryoid cultures on different days. White arrowheads mark spontaneous beating sites. Scale bar, 100 µm.

## SUPPLEMENTARY TABLES

**Supplementary Table 1.** List of differentially expressed genes, signaling pathways, regulons, and correlation coefficients from scRNA-seq analysis.

**Supplementary Table 2.**
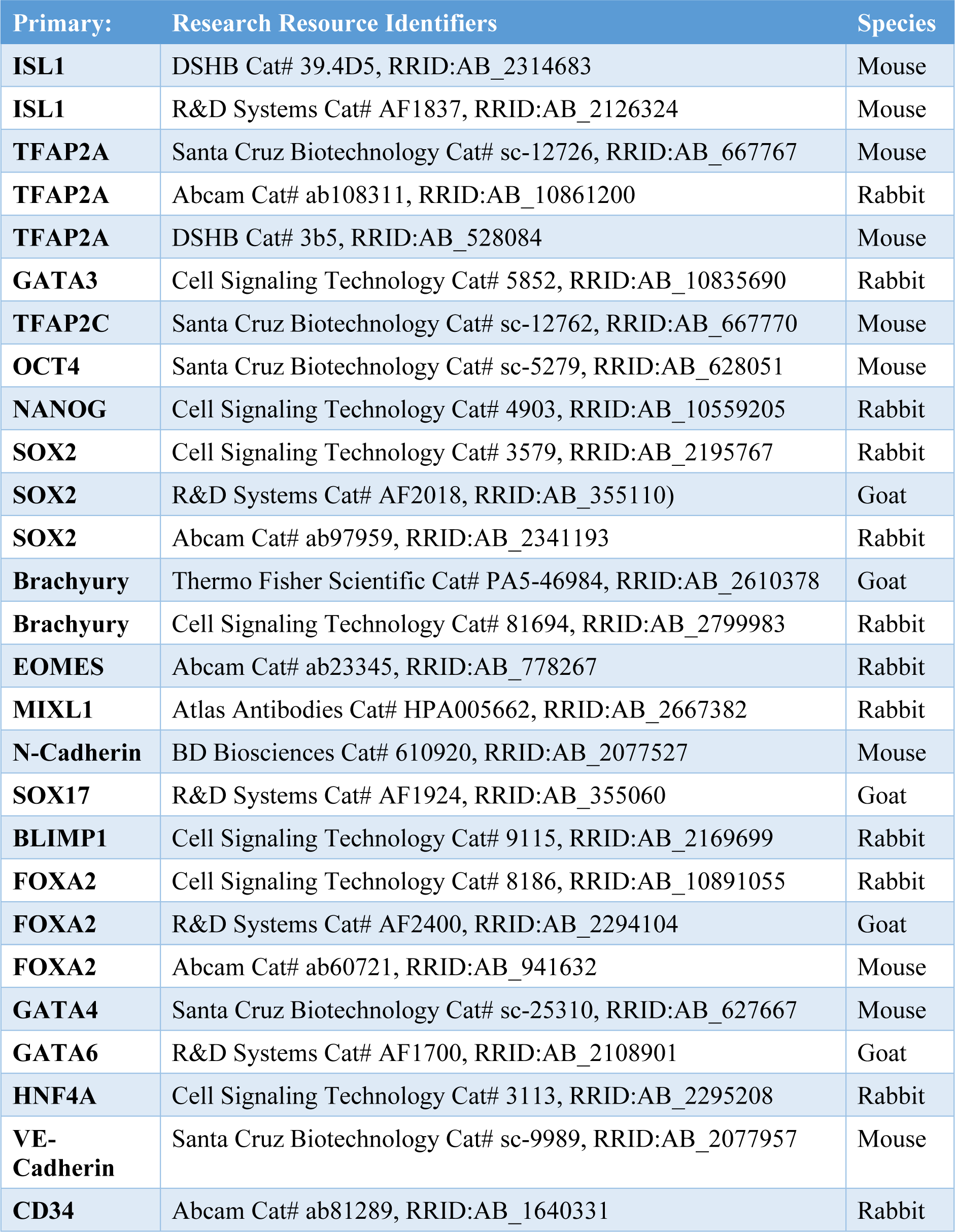
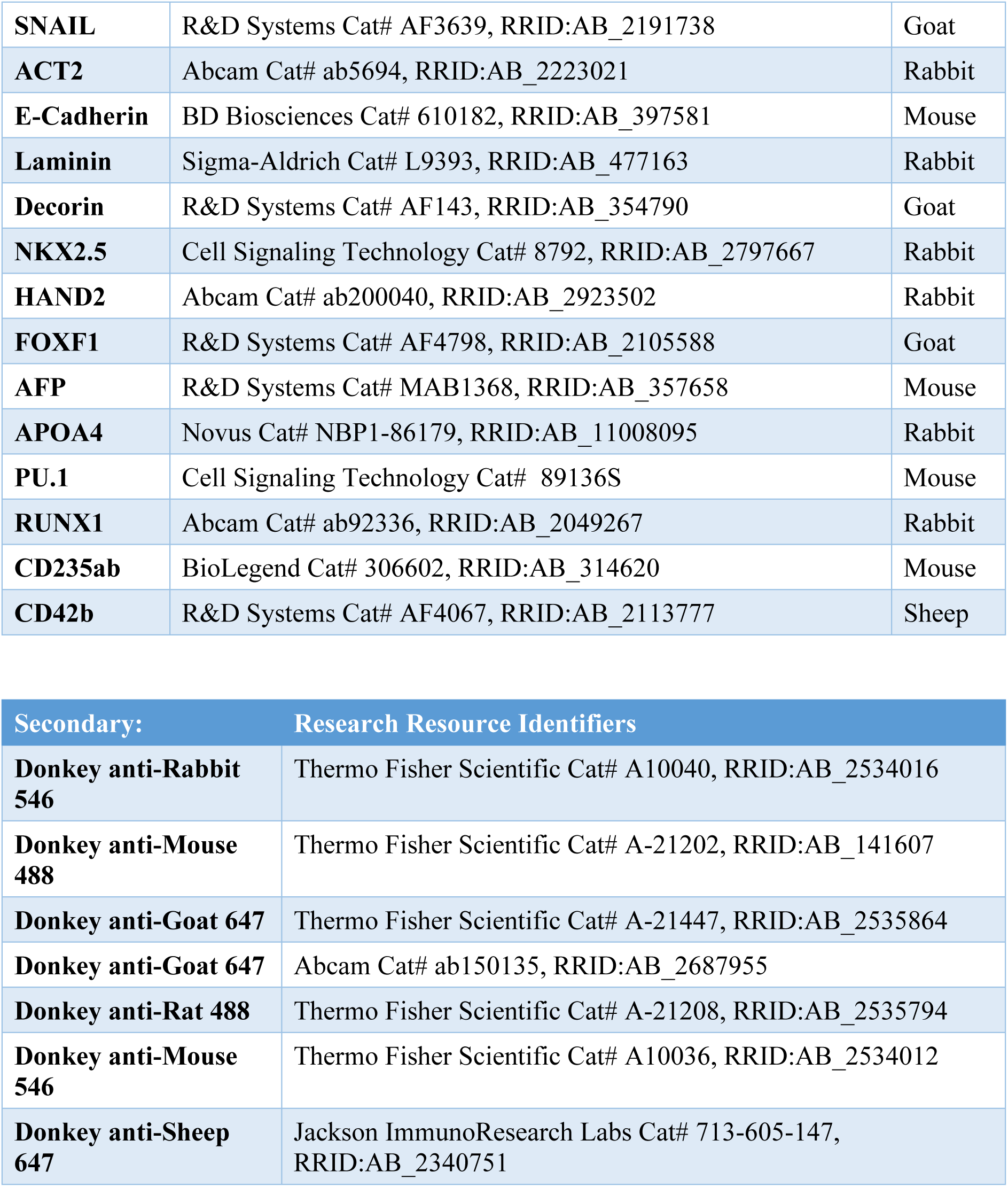
List of antibodies used in immunocytochemistry and immunohistochemistry.

**Supplementary Table 3.** List of antibodies used in flow cytometry.

| **Protein (clone):** | **Fluoro** | **Research Resource Identifiers** |
| --- | --- | --- |
| **CD15 (HI98)** | BUV395 | BD Biosciences Cat# 563872, RRID:AB_2738461 |
| **CD3 (SK7)** | BUV615 | BD Biosciences Cat# 751252, RRID:AB_2875269 |
| **CD14 (M5E2)** | BUV737 | BD Biosciences Cat# 612764, RRID:AB_2870095 |
| **CD117 (104D2)** | BV421 | BD Biosciences Cat# 563856, RRID:AB_2738452 |
| **CD34 (581)** | BV510 | BioLegend Cat# 343527, RRID:AB_2561985 |
| **CD33 (P67.6)** | BV605 | BioLegend Cat# 366611, RRID:AB_2566404 |
| **CD49d (9F10)** | BV711 | BioLegend Cat# 304331, RRID:AB_2687197 |
| **CD45 (HI30)** | BV785 | BioLegend Cat# 304047, RRID:AB_2563128 |
| **CD19 (HIB19)** | FITC | BioLegend Cat# 302205, RRID:AB_314235 |
| **CD43 (CD43-10G7)** | PE | BioLegend Cat# 343203, RRID:AB_1659201 |
| **CD56 (MEM-188)** | PE-Cy5 | BioLegend Cat# 304607, RRID:AB_314449 |
| **CD235ab (HIR2)** | PE-Cy7 | BioLegend Cat# 306619, RRID:AB_2716163 |
| **CD31 (WM59)** | APC | BioLegend Cat# 303115, RRID:AB_1877152 |
| **CD42b (HIP1)** | AF700 | BioLegend Cat# 303927, RRID:AB_2715928 |

## REFERENCES

1 Tam, P. P. L. & Loebel, D. A. F. Gene function in mouse embryogenesis: Get set for gastrulation. Nat Rev Genet 8, 368–381 (2007).

2 Arnold, S. J. & Robertson, E. J. Making a commitment: Cell lineage allocation and axis patterning in the early mouse embryo. Nat. Rev. Mol. Cell Biol. 10, 91–103 (2009).

3 Solnica-Krezel, L. Gastrulation: From embryonic pattern to form (2020).

4 Sheng, G. J., Arias, A. M. & Sutherland, A. The primitive streak and cellular principles of building an amniote body through gastrulation. Science 374, eabg1727 (2021).

5 Warmflash, A. et al. A method to recapitulate early embryonic spatial patterning in human embryonic stem cells. Nat Methods 11, 847–854 (2014).

6 Shao, Y. et al. A pluripotent stem cell-based model for post-implantation human amniotic sac development. Nat Commun 8, 208 (2017).

7 Simunovic, M. et al. A 3d model of a human epiblast reveals BMP4-driven symmetry breaking. Nat. Cell Biol. 21, 900–910 (2019).

8 Moris, N. et al. An in vitro model of early anteroposterior organization during human development. Nature 582, 410–415 (2020).

9 Ai, Z. et al. Dissecting peri-implantation development using cultured human embryos and embryo-like assembloids. Cell Res 33, 661–678 (2023).

10 Karvas, R. M. et al. 3d-cultured blastoids model human embryogenesis from pre-implantation to early gastrulation stages. Cell Stem Cell 30, 1148–1165 e1147 (2023).

11 Liu, L. et al. Modeling post-implantation stages of human development into early organogenesis with stem-cell-derived peri-gastruloids. Cell 186, 3776–3792 (2023).

12 Oldak, B. et al. Complete human day 14 post-implantation embryo models from naïve ES cells. Nature 622, 562–573 (2023).

13 Pedroza, M. et al. Self-patterning of human stem cells into post-implantation lineages. Nature 622, 574–583 (2023).

14 Weatherbee, B. A. T. et al. Pluripotent stem cell-derived model of the post-implantation human embryo. Nature 622, 584–593 (2023).

15 Hislop, J. et al. Modeling post-implantation human development to yolk sac blood emergence. Nature 626, 367–376 (2023).

16 Shahbazi, M. N., Siggia, E. D. & Zernicka-Goetz, M. Self-organization of stem cells into embryos: A window on early mammalian development. Science 364, 948–951 (2019).

17 Fu, J., Warmflash, A. & Lutolf, M. P. Stem-cell-based embryo models for fundamental research and translation. Nat Mater 20, 132–144 (2021).

18 Rossant, J. & Tam, P. P. L. Opportunities and challenges with stem cell-based embryo models. Stem Cell Rep 16, 1031–1038 (2021).

19 O’Rahilly, R. & Muller, F. Developmental stages in human embryos: Revised and new measurements. Cells Tissues Organs 192, 73–84 (2010).

20 Ross, C. & Boroviak, T. E. Origin and function of the yolk sac in primate embryogenesis. Nat Commun 11, 3760 (2020).

21 Nowotschin, S. et al. The emergent landscape of the mouse gut endoderm at single-cell resolution. Nature 569, 361–367 (2019).

22 Mikkola, H. K. A. Yolk sac steps up to the plate. J Exp Med 219, e20212315 (2022).

23 Simunovic, M., Siggia, E. D. & Brivanlou, A. H. In vitro attachment and symmetry breaking of a human embryo model assembled from primed embryonic stem cells. Cell Stem Cell 29, 962–972 (2022).

24 Sanaki-Matsumiya, M. et al. Periodic formation of epithelial somites from human pluripotent stem cells. Nature Communications 13, 2325 (2022).

25 Miao, Y. et al. Reconstruction and deconstruction of human somitogenesis in vitro. Nature 614, 500–508 (2023).

26 Yaman, Y. I. & Ramanathan, S. Controlling human organoid symmetry breaking reveals signaling gradients drive segmentation clock waves. Cell 186, 513–527.e519 (2023).

27 Yamanaka, Y. et al. Reconstituting human somitogenesis in vitro. Nature 614, 509–520 (2023).

28 Yang, R. et al. Amnion signals are essential for mesoderm formation in primates. Nat Commun 12, 5126 (2021).

29 Mackinlay, K. M. L. et al. An in vitro stem cell model of human epiblast and yolk sac interaction. Elife 10, e63930 (2021).

30 Chen, L. et al. The nuclear receptor hnf4 drives a brush border gene program conserved across murine intestine, kidney, and embryonic yolk sac. Nat Commun 12, 2886 (2021).

31 Brennan, J. et al. Nodal signalling in the epiblast patterns the early mouse embryo. Nature 411, 965–969 (2001).

32 Rivera-Pérez, J. A. & Magnuson, T. Primitive streak formation in mice is preceded by localized activation of brachyury and wnt3. Dev Biol 288, 363–371 (2005).

33 Ben-Haim, N. et al. The nodal precursor acting via activin receptors induces mesoderm by maintaining a source of its convertases and bmp4. Dev Cell 11, 313–323 (2006).

34 Chhabra, S. et al. Dissecting the dynamics of signaling events in the bmp, wnt, and nodal cascade during self-organized fate patterning in human gastruloids. Plos Biol 17 (2019).

35 Muncie, J. M. et al. Mechanical tension promotes formation of gastrulation-like nodes and patterns mesoderm specification in human embryonic stem cells. Dev Cell 55, 679–694 (2020).

36 Tesar, P. J. et al. New cell lines from mouse epiblast share defining features with human embryonic stem cells. Nature 448, 196–199 (2007).

37 O’Leary, T. et al. Tracking the progression of the human inner cell mass during embryonic stem cell derivation. Nat Biotechnol 30, 278–282 (2012).

38 Zheng, Y. et al. Controlled modelling of human epiblast and amnion development using stem cells. Nature 573, 421–425 (2019).

39 Karzbrun, E. et al. Human neural tube morphogenesis in vitro by geometric constraints. Nature 599, 268–272 (2021).

40 Xue, X. et al. Mechanics-guided embryonic patterning of neuroectoderm tissue from human pluripotent stem cells. Nat Mater 17, 633–641 (2018).

41 Minn, K. T. et al. High-resolution transcriptional and morphogenetic profiling of cells from micropatterned human esc gastruloid cultures. Elife 9, e59445 (2020).

42 Luckett, W. P. Origin and differentiation of the yolk sac and extraembryonic mesoderm in presomite human and rhesus monkey embryos. Am. J. Anat. 152, 59–97 (1978).

43 Bianchi, D. W., Wilkinshaug, L. E., Enders, A. C. & Hay, E. D. Origin of extraembryonic mesoderm in experimental-animals: Relevance to chorionic mosaicism in humans. Am J Med Genet 46, 542–550 (1993).

44 Nakamura, T. et al. A developmental coordinate of pluripotency among mice, monkeys and humans. Nature 537, 57–62 (2016).

45 Hofbauer, P. et al. Cardioids reveal self-organizing principles of human cardiogenesis. Cell 184, 3299–3317 (2021).

46 Tyser, R. C. V. et al. Single-cell transcriptomic characterization of a gastrulating human embryo. Nature 600, 285–289 (2021).

47 Schmidt, C. et al. Multi-chamber cardioids unravel human heart development and cardiac defects. Cell 186, 5587–5605 (2023).

48 Zhai, J. et al. Primate gastrulation and early organogenesis at single-cell resolution. Nature 612, 732–738 (2022).

49 Chen, D. et al. Human primordial germ cells are specified from lineage-primed progenitors. Cell Rep 29, 4568–4582 (2019).

50 Sasaki, K. et al. The germ cell fate of cynomolgus monkeys is specified in the nascent amnion. Dev Cell 39, 169–185 (2016).

51 Morgani, S. M. et al. Micropattern differentiation of mouse pluripotent stem cells recapitulates embryo regionalized cell fate patterning. Elife 7, e32839 (2018).

52 Zheng, Y. et al. Single-cell analysis of embryoids reveals lineage diversification roadmaps of early human development. Cell Stem Cell 29, 1402–1419 (2022).

53 Manfrin, A. et al. Engineered signaling centers for the spatially controlled patterning of human pluripotent stem cells. Nat Methods 16, 640–648 (2019).

54 Tamaoki, N. et al. Self-organized yolk sac-like organoids allow for scalable generation of multipotent hematopoietic progenitor cells from induced pluripotent stem cells. Cell Rep Methods 3, 100460 (2023).

55 Ivanovs, A. et al. Human haematopoietic stem cell development: From the embryo to the dish. Development 144, 2323–2337 (2017).

56 Enders, A. C. & King, B. F. Formation and differentiation of extraembryonic mesoderm in the rhesus-monkey. Am. J. Anat. 181, 327–340 (1988).

57 Xue, X. et al. A patterned human neural tube model using microfluidic gradients. Nature 628, 391–399 (2024).

58 Technau, U. Brachyury, the blastopore and the evolution of the mesoderm. Bioessays 23, 788–794 (2001).

59 Bulger, E. A. et al. Tbxt dose sensitivity and the decoupling of nascent mesoderm specification from emt progression in 2d human gastruloids. Development 151, dev202516 (2024).

60 Sturgeon, C. M. et al. Wnt signaling controls the specification of definitive and primitive hematopoiesis from human pluripotent stem cells. Nat Biotechnol 32, 554–561 (2014).

61 Goh, I. et al. Yolk sac cell atlas reveals multiorgan functions during human early development. Science 381, eadd7564 (2023).

62 Canu, G. & Ruhrberg, C. First blood: The endothelial origins of hematopoietic progenitors. Angiogenesis 24, 199–211 (2021).

63 Lange, L., Morgan, M. & Schambach, A. The hemogenic endothelium: A critical source for the generation of psc-derived hematopoietic stem and progenitor cells. Cell Mol Life Sci 78, 4143–4160 (2021).

64 Calvanese, V. et al. Mapping human haematopoietic stem cells from haemogenic endothelium to birth. Nature 604, 534–540 (2022).

65 Sugimura, R. et al. Haematopoietic stem and progenitor cells from human pluripotent stem cells. Nature 545, 432–438 (2017).

66 Motazedian, A. et al. Multipotent rag1+ progenitors emerge directly from haemogenic endothelium in human pluripotent stem cell-derived haematopoietic organoids. Nat. Cell Biol. 22, 60–73 (2020).

67 Dzierzak, E. & Philipsen, S. Erythropoiesis: Development and differentiation. Csh Perspect Med 3, a011601 (2013).

68 Italiano, J. E. & Hartwig, J. H. Megakaryocyte development and platelet formation. Platelets, 2nd Edition, 23–44 (2007).

69 Machlus, K. R. & Italiano, J. E. The incredible journey: From megakaryocyte development to platelet formation. J Cell Biol 201, 785–796 (2013).

70 Calvanese, V. & Mikkola, H. K. A. The genesis of human hematopoietic stem cells. Blood 142, 519–532 (2023).

71 Botting, R. A. et al. Multi-organ functions of yolk sac during human early development. bioRxiv, 2022.2008.2003.502475 (2022).

72 Bergmann, S. et al. Spatial profiling of early primate gastrulation in utero. Nature 609, 136–143 (2022).

73 Cui, G. et al. Spatial molecular anatomy of germ layers in the gastrulating cynomolgus monkey embryo. Cell Rep 40, 111285 (2022).

74 Peng, G. D. et al. Spatial transcriptome for the molecular annotation of lineage fates and cell identity in mid-gastrula mouse embryo. Dev Cell 36, 681–697 (2016).

75 Mascetti, V. L. & Pedersen, R. A. Human-mouse chimerism validates human stem cell pluripotency. Cell Stem Cell 18, 67–72 (2016).

76 Zheng, H. Q. et al. Generating hematopoietic cells from human pluripotent stem cells: Approaches, progress and challenges. Cell Regen 12, 31 (2023).

77 Atkins, M. H. et al. Modeling human yolk sac hematopoiesis with pluripotent stem cells. J Exp Med 219, e20211924 (2021).

78 Lim, W. F. et al. Hematopoietic cell differentiation from embryonic and induced pluripotent stem cells. Stem Cell Res Ther 4, 71 (2013).

79 Yamasaki, J. et al. Vitrification and transfer of cynomolgus monkey (macaca fascicularis) embryos fertilized by intracytoplasmic sperm injection. Theriogenology 76, 33–38 (2011).

80 McClelland, K. S., Ng, E. T. & Bowles, J. Agarose/gelatin immobilisation of tissues or embryo segments for orientated paraffin embedding and sectioning. Differentiation 91, 68–71 (2016).

81 Schindelin, J., et al. Fiji: An open-source platform for biological-image analysis. Nat Methods 9, 676-682 (2012).

82 Hao, Y. H. et al. Integrated analysis of multimodal single-cell data. Cell 184, 3573–3587 (2021).

83 Stuart, T. et al. Comprehensive integration of single-cell data. Cell 177, 1888–1902 (2019).

84 Butler, A. et al. Integrating single-cell transcriptomic data across different conditions, technologies, and species. Nat Biotechnol 36, 411–420 (2018).

85 Satija, R. et al. Spatial reconstruction of single-cell gene expression data. Nat Biotechnol 33, 495–502 (2015).

86 Jin, S. Q. et al. Inference and analysis of cell-cell communication using cellchat. Nat Commun 12, 1088 (2021).

87 Angerer, P. et al. Destiny: Diffusion maps for large-scale single-cell data in r. Bioinformatics 32, 1241–1243 (2016).

88 Farrell, J. A. et al. Single-cell reconstruction of developmental trajectories during zebrafish embryogenesis. Science 360, eaar3131 (2018).

89 Uzquiano, A. et al. Proper acquisition of cell class identity in organoids allows definition of fate specification programs of the human cerebral cortex. Cell 185, 3770–3788 (2022).

90 Aibar, S. et al. Scenic: Single-cell regulatory network inference and clustering. Nat Methods 14, 1083–1086 (2017).

91 Wu, T. et al. Clusterprofiler 4.0: A universal enrichment tool for interpreting omics data. Innovation 2, 100141 (2021).

92 Yu, G. C., Wang, L. G., Han, Y. Y. & He, Q. Y. Clusterprofiler: An r package for comparing biological themes among gene clusters. Omics 16, 284–287 (2012).

