## Extended Data Figure for "A transgene-free, human peri-gastrulation embryo model with trilaminar embryonic disc-, amnion- and yolk sac-like structures"

### Extended Data Figure 1

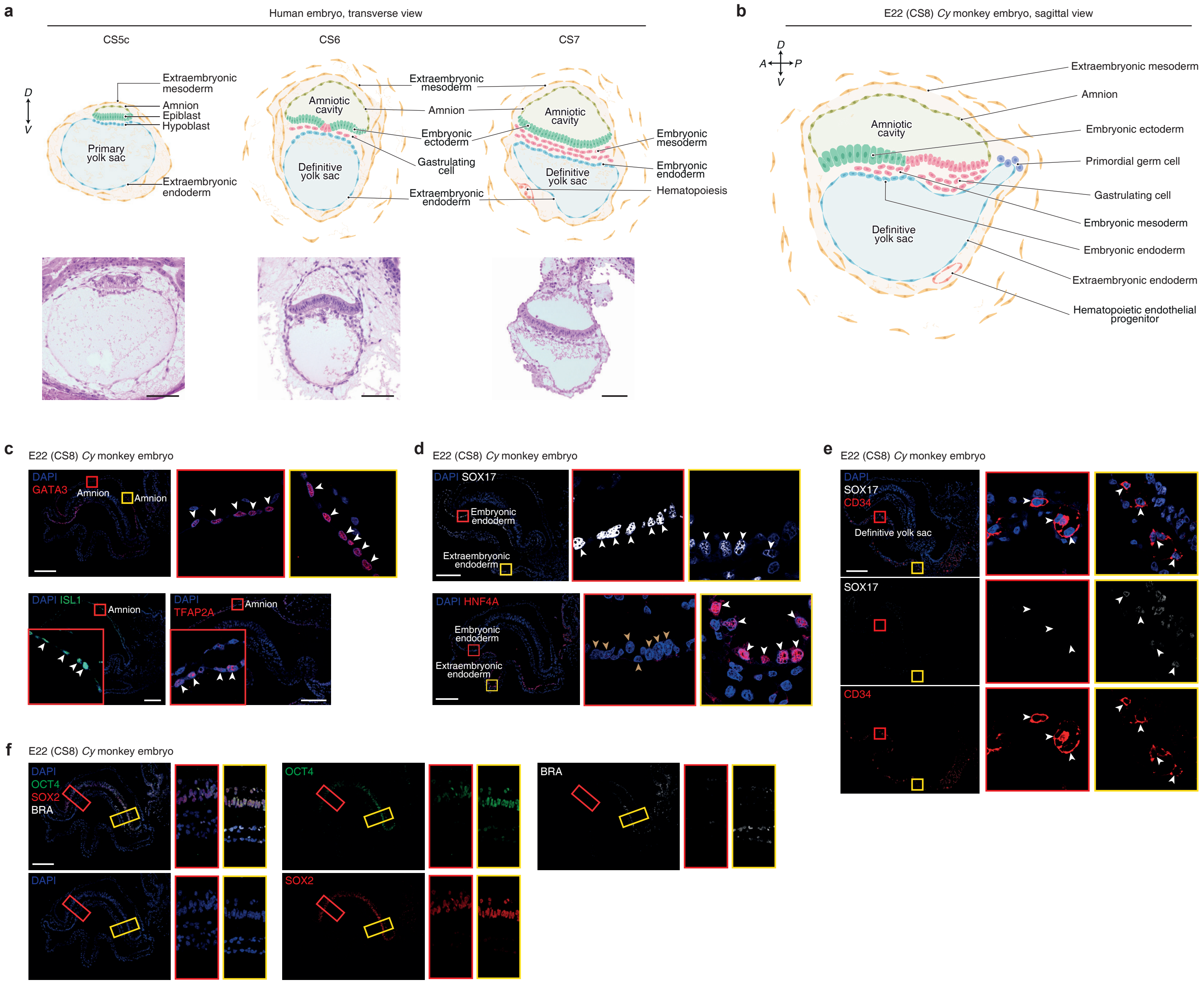

### Extended Data Figure 2

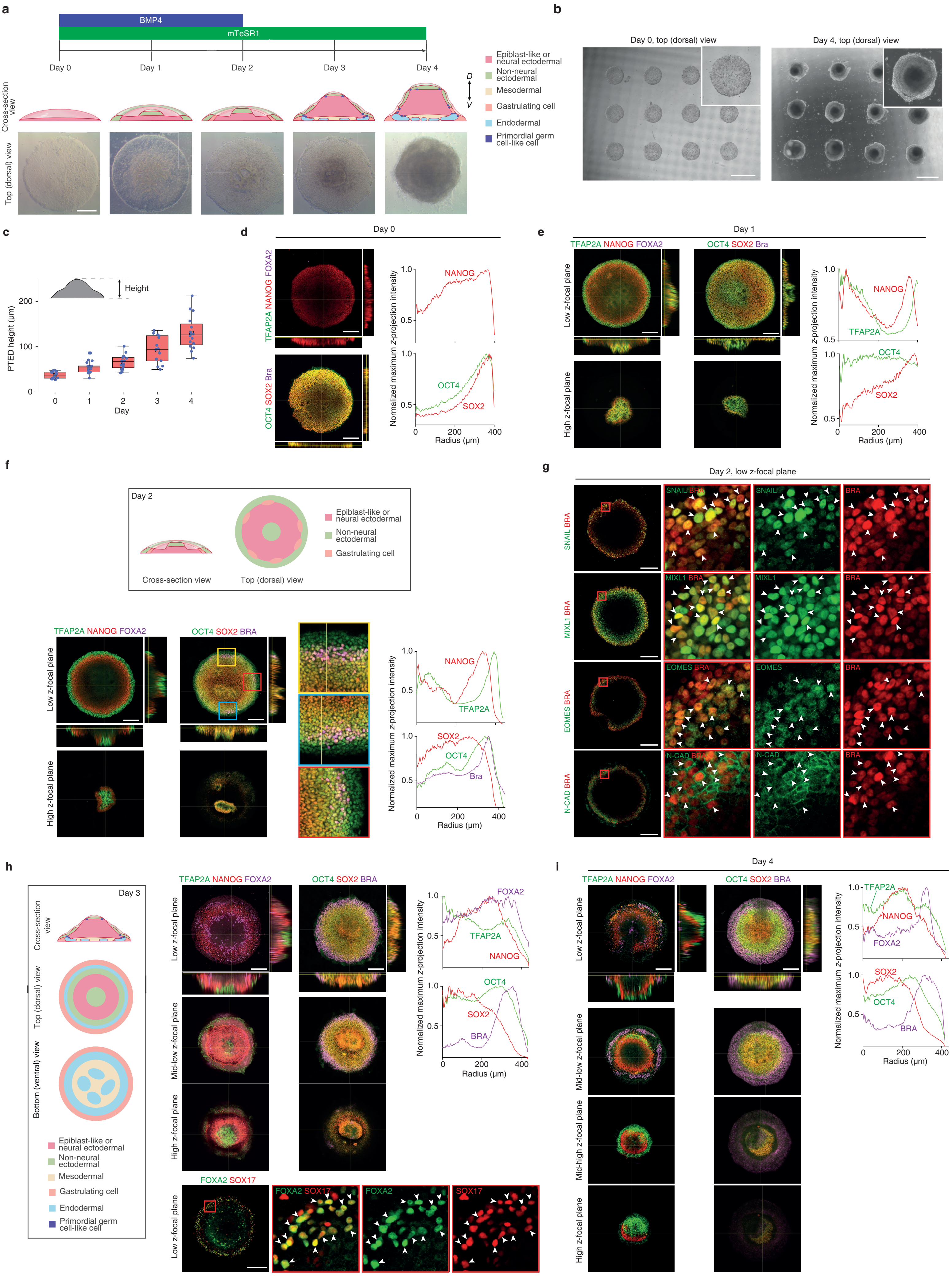

### Extended Data Figure 3

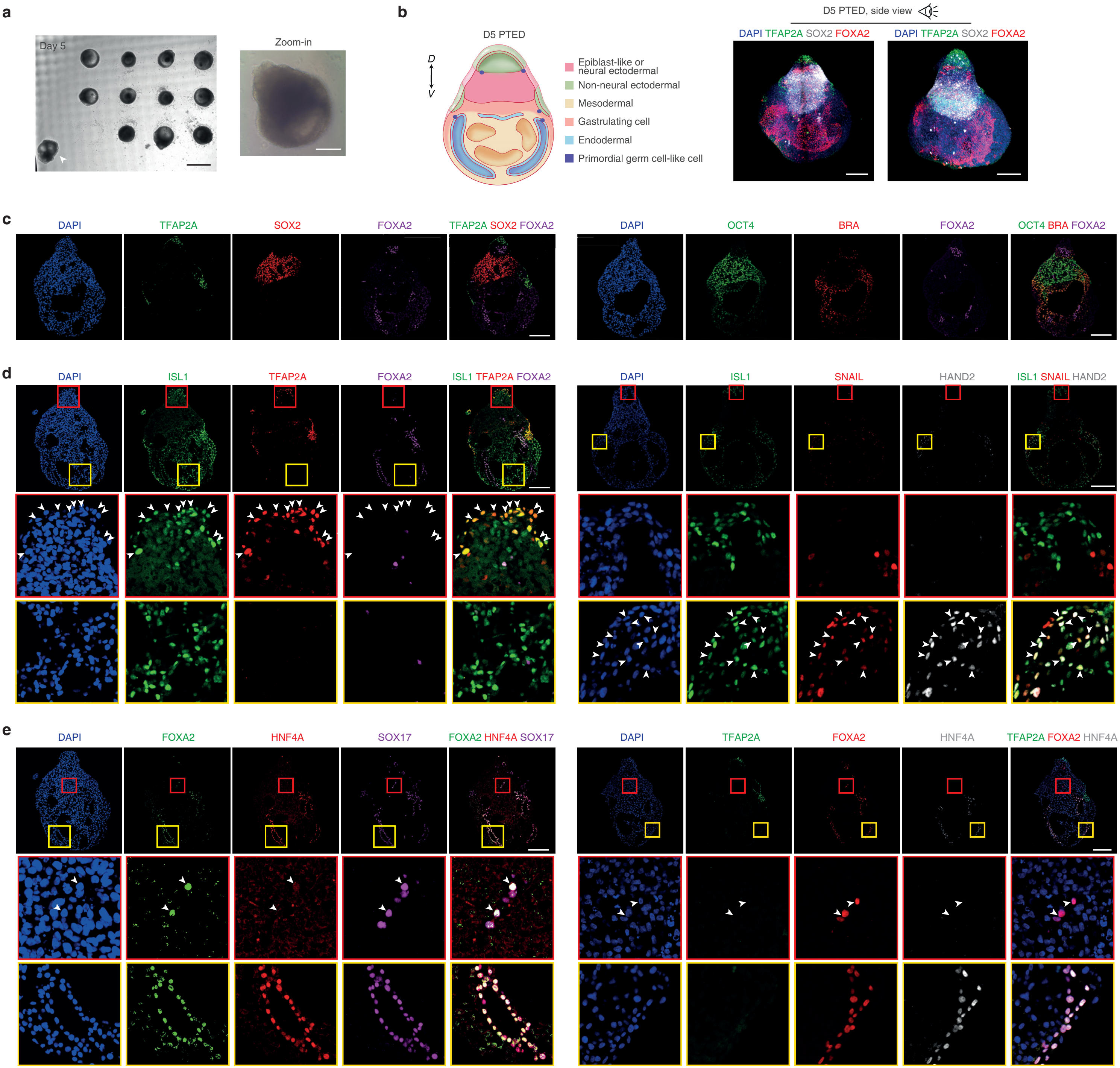

### Extended Data Figure 4

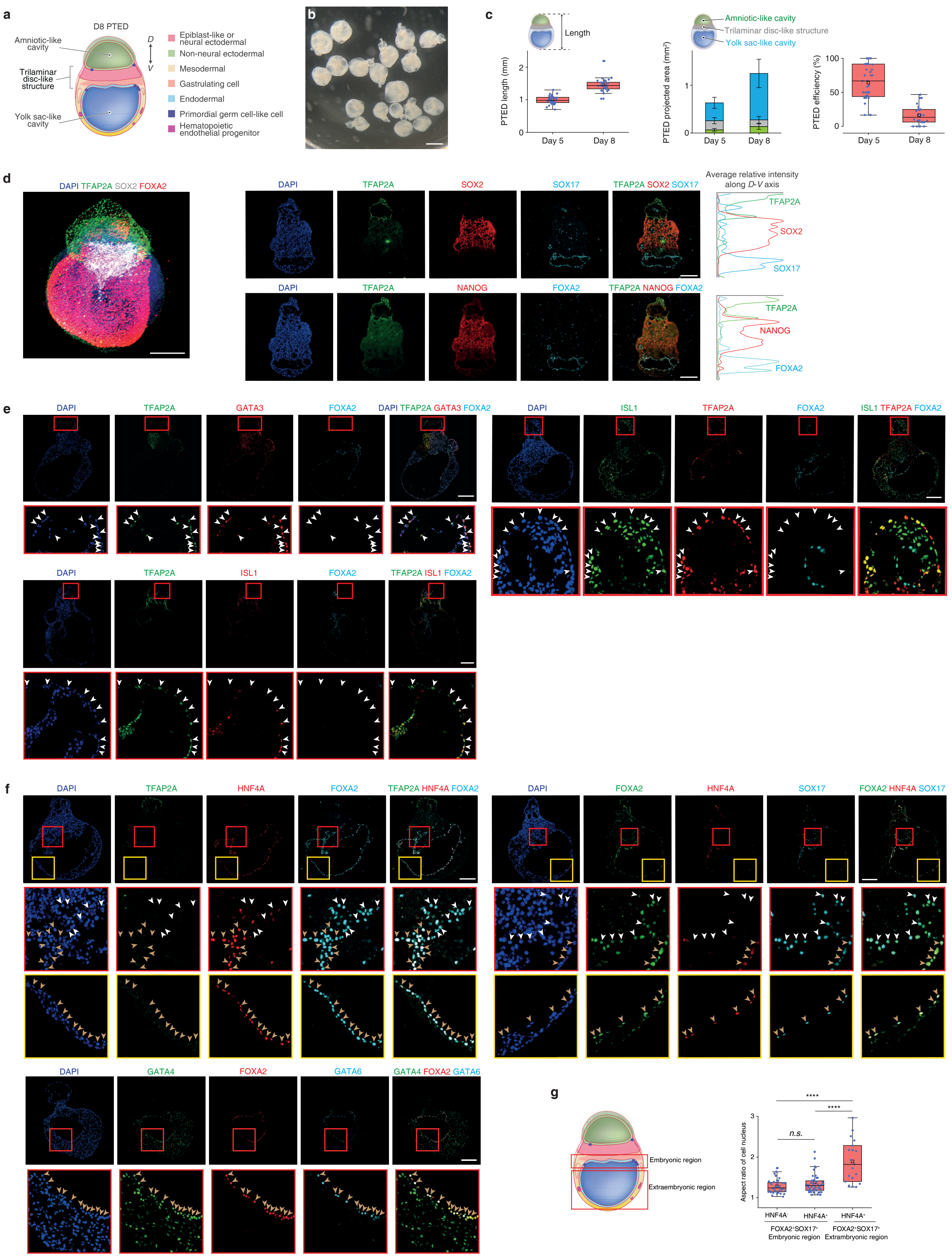

### Extended Data Figure 5

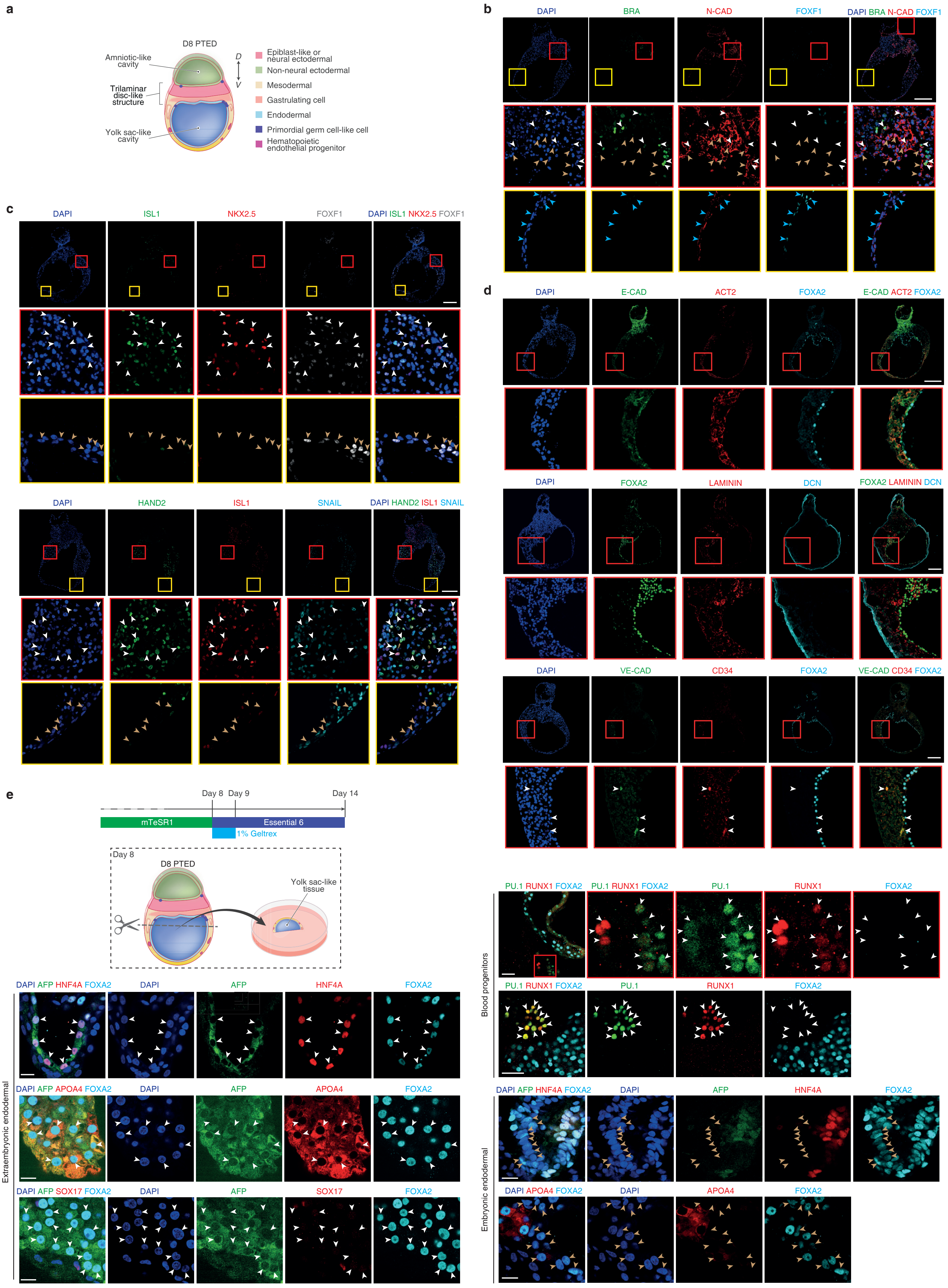

### Extended Data Figure 6

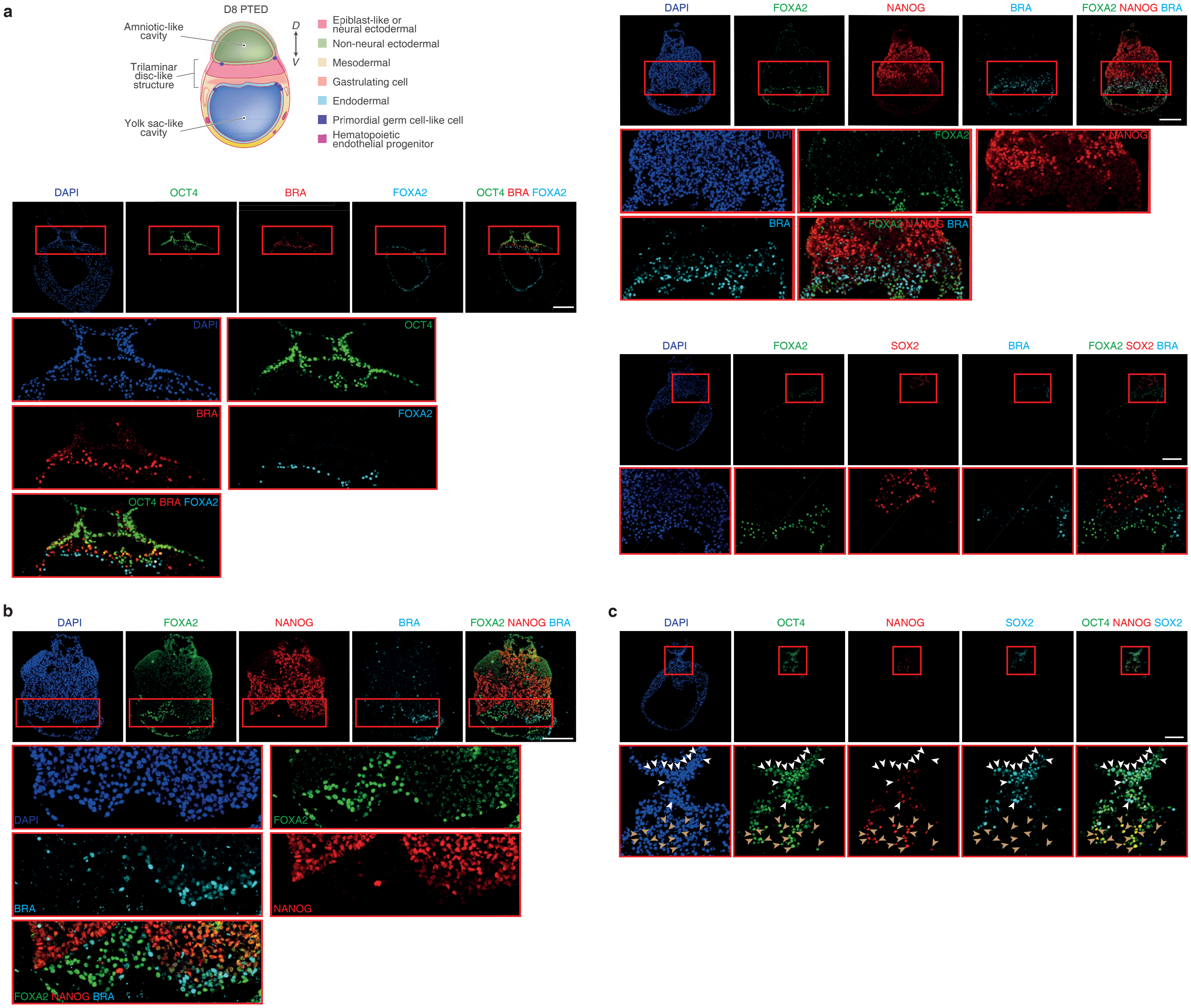

### Extended Data Figure 7

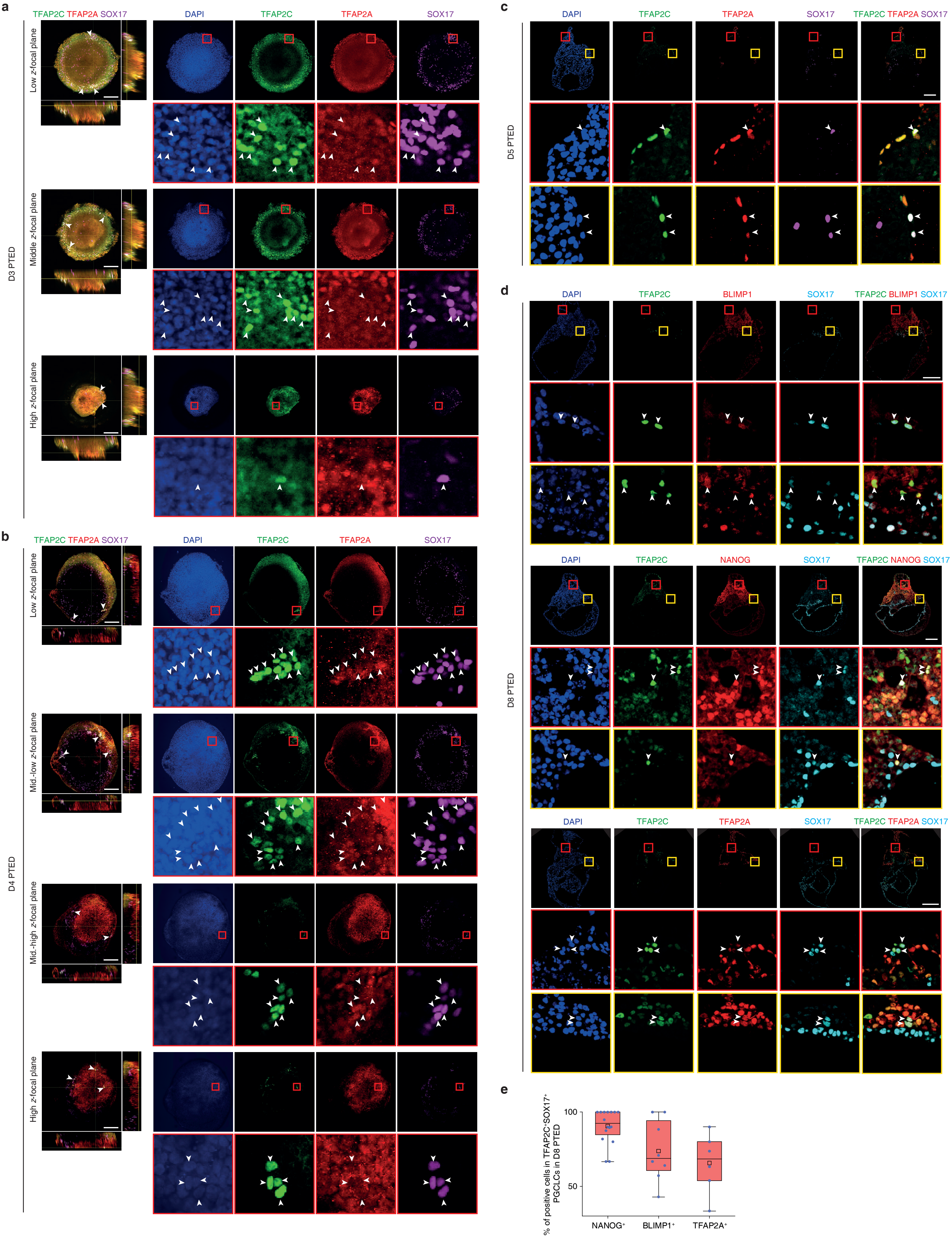

### Extended Data Figure 8

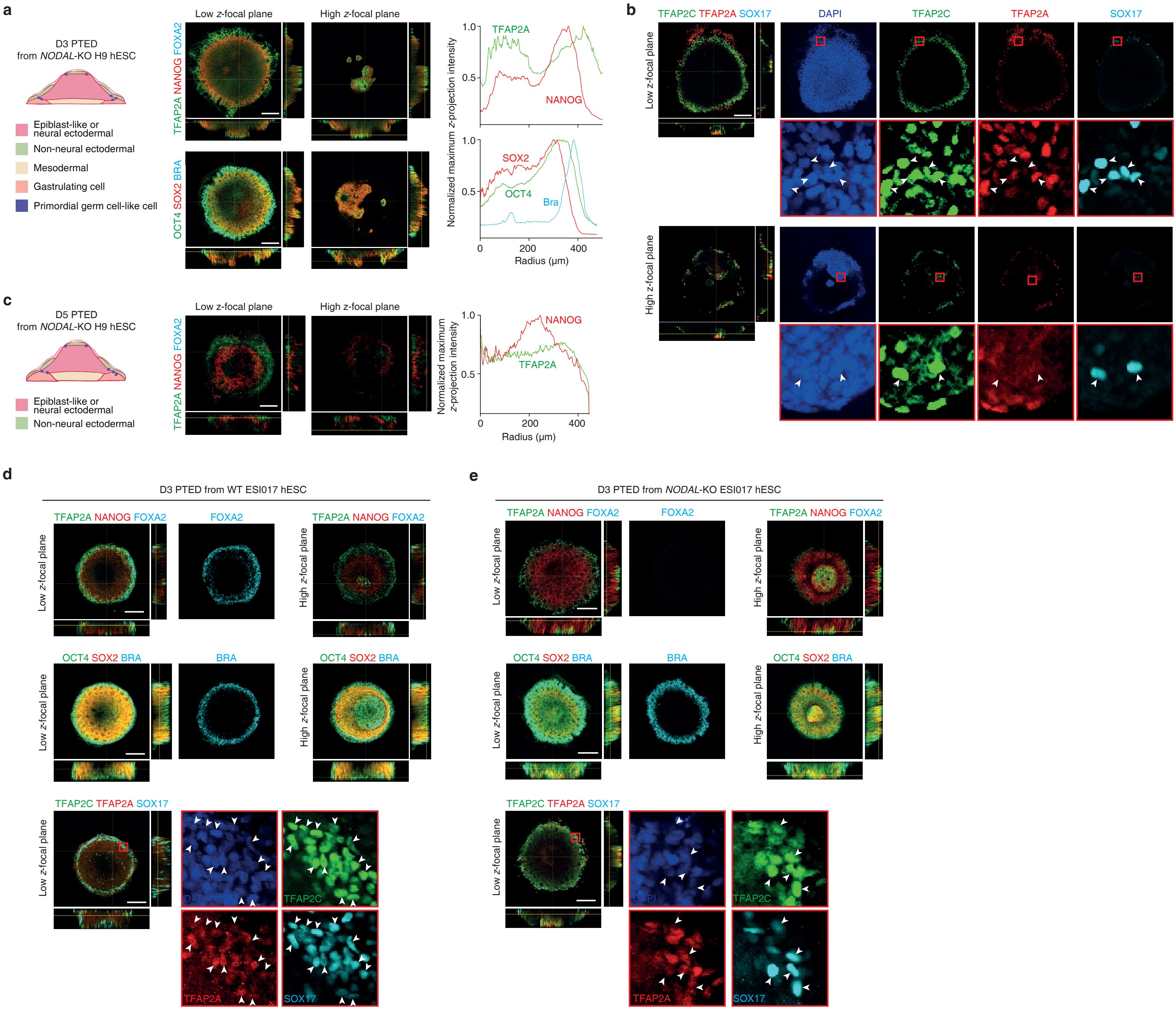

### Extended Data Figure 9

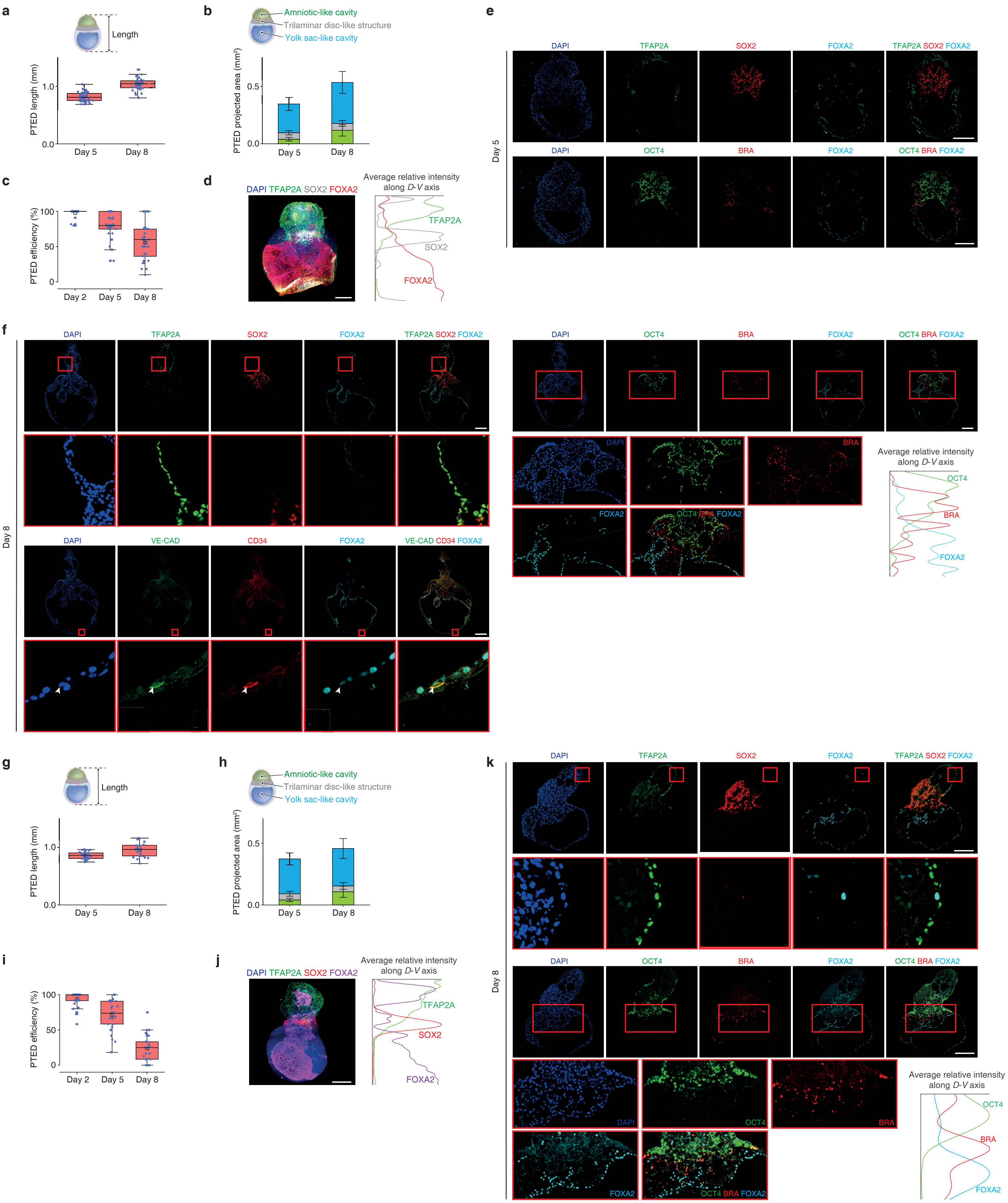

### Extended Data Figure 10

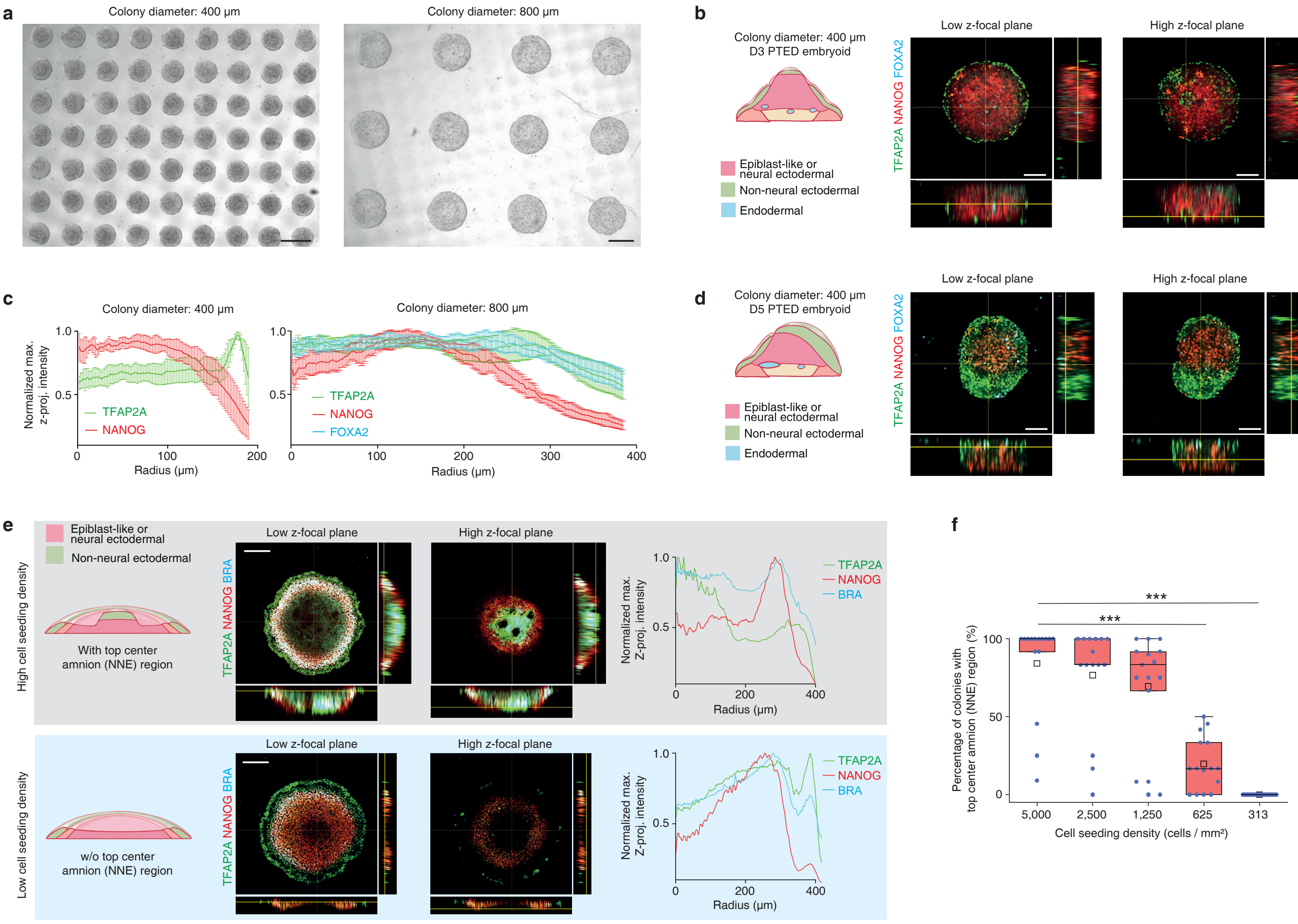

### Extended Data Figure 11

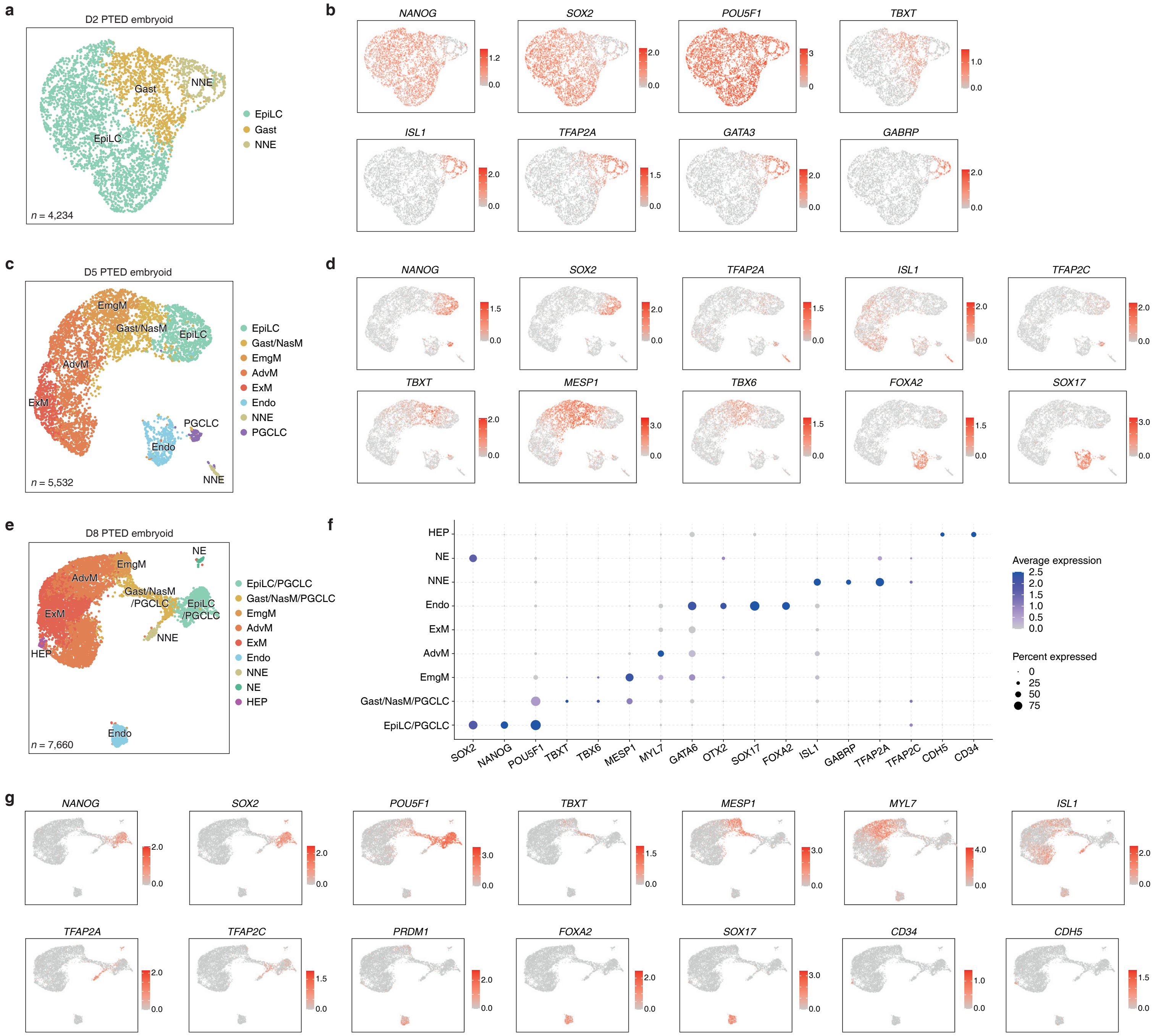

### Extended Data Figure 12

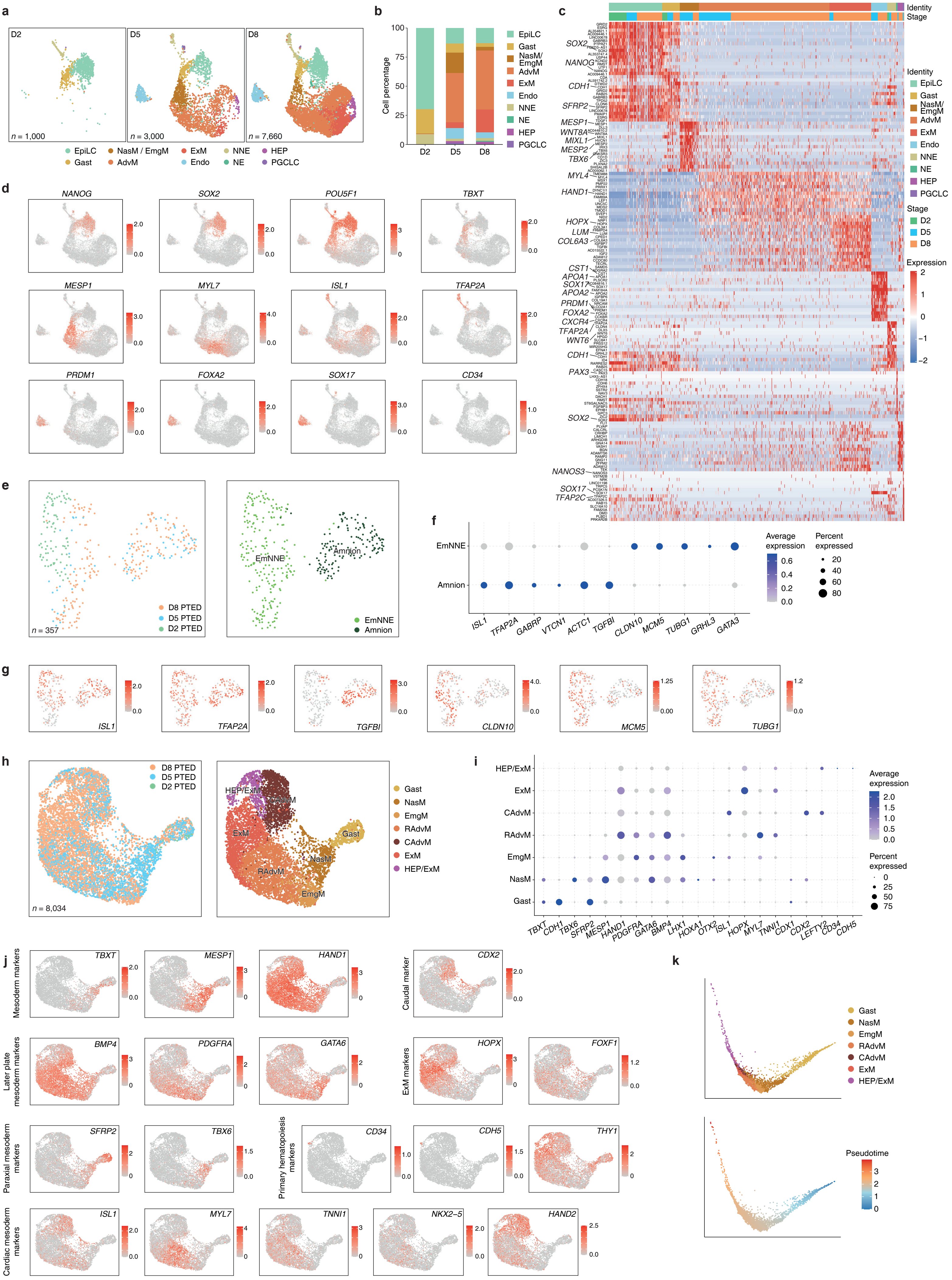

### Extended Data Figure 13

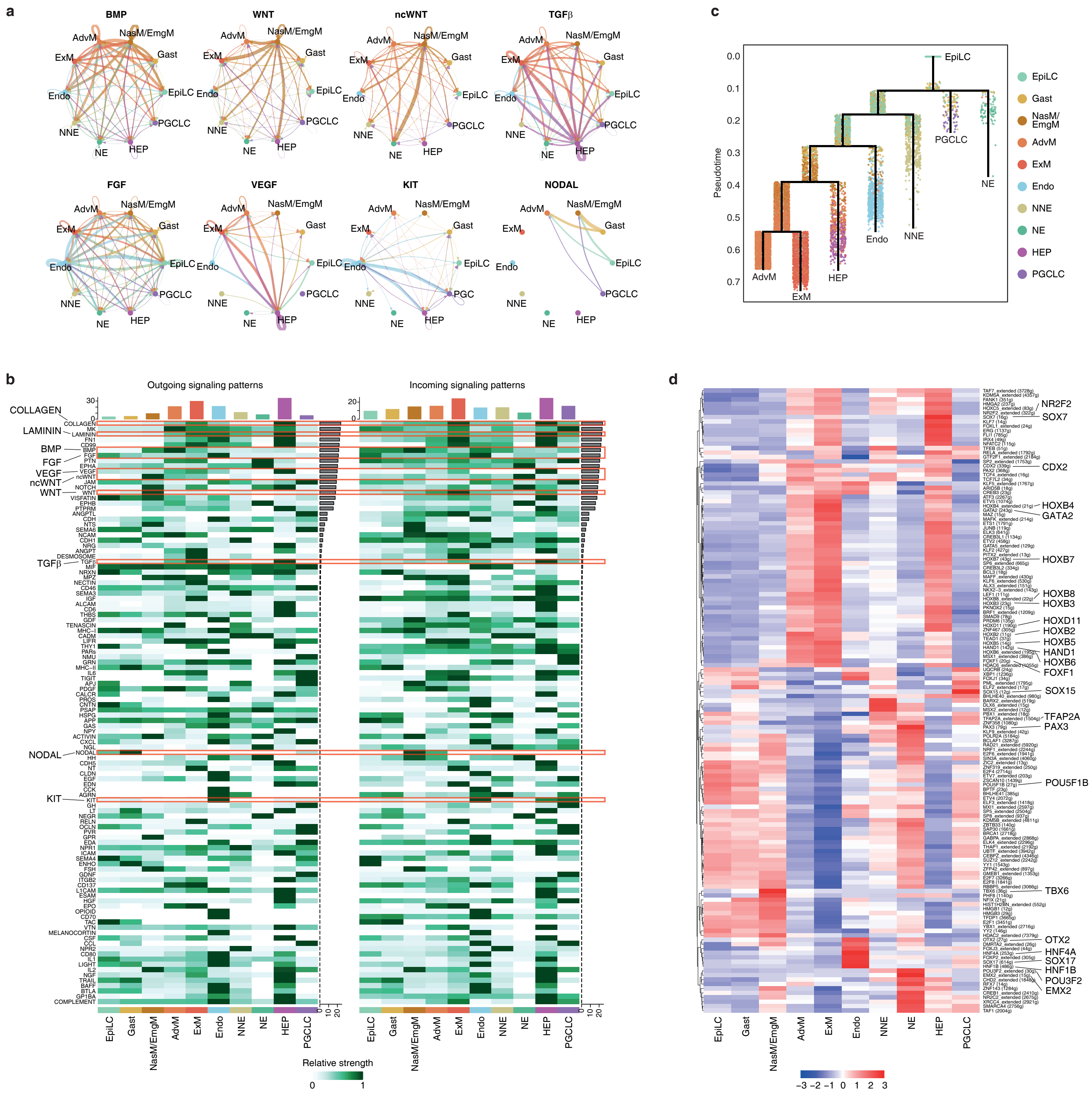

### Extended Data Figure 14

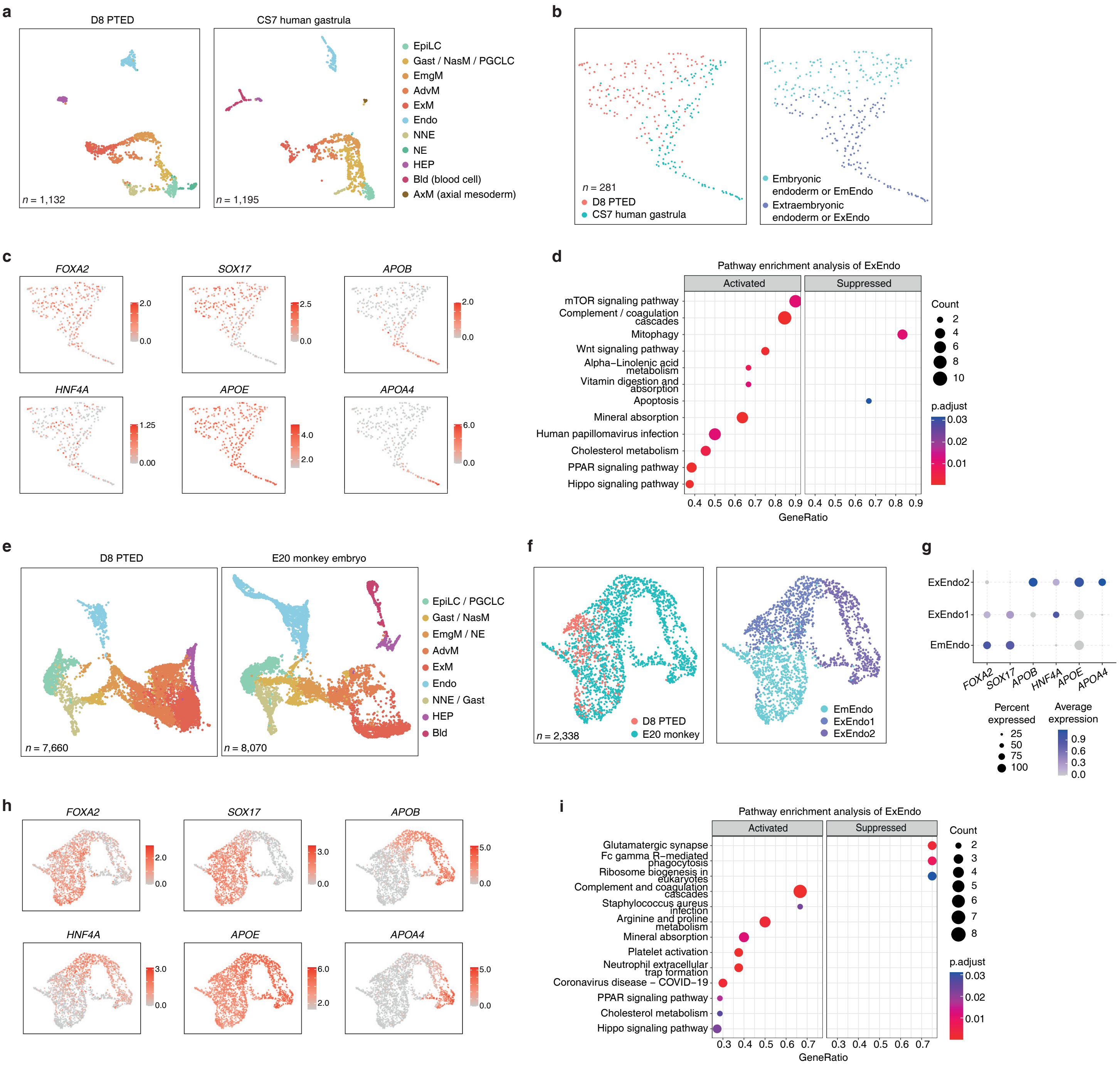

### Extended Data Figure 15

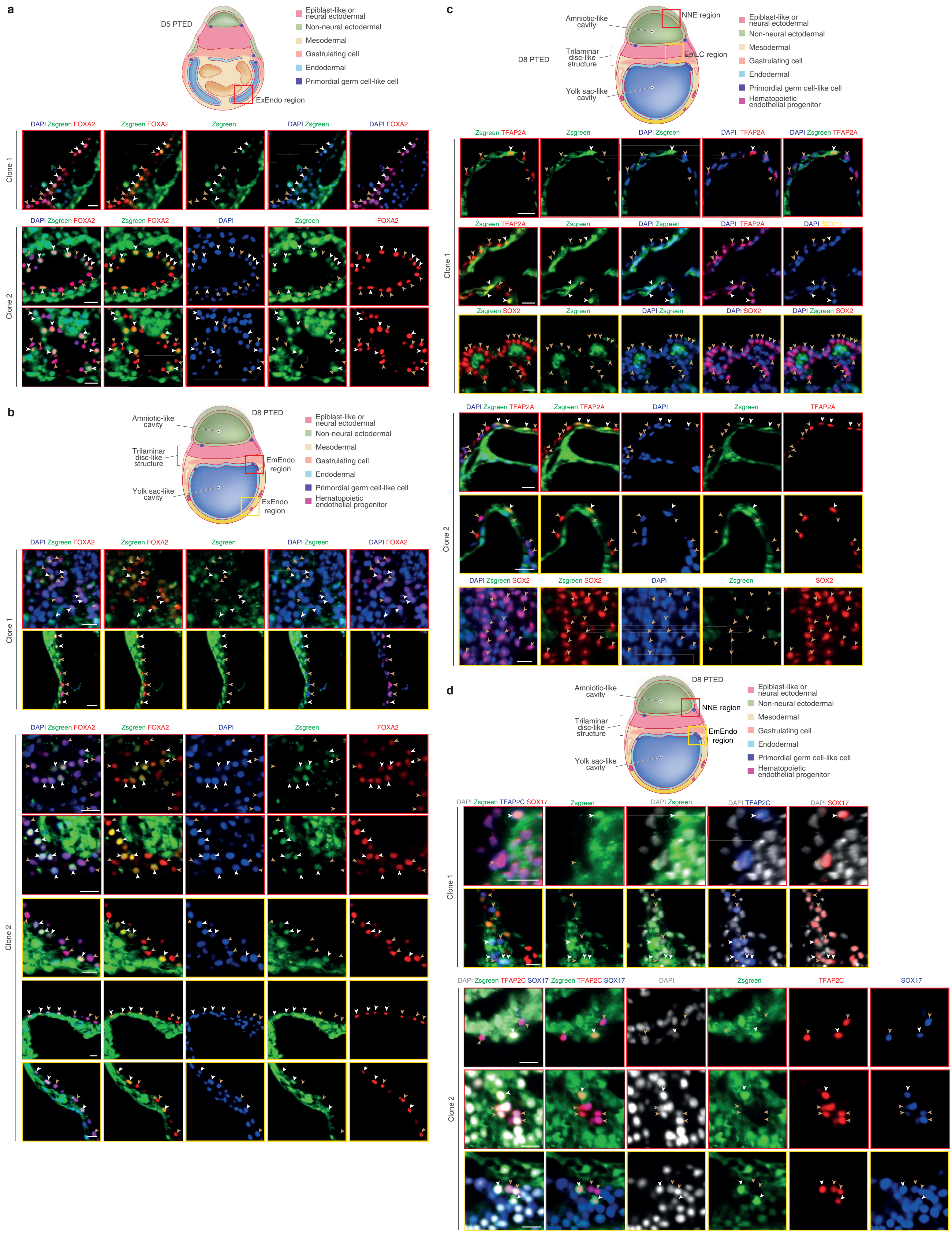

### Extended Data Figure 16

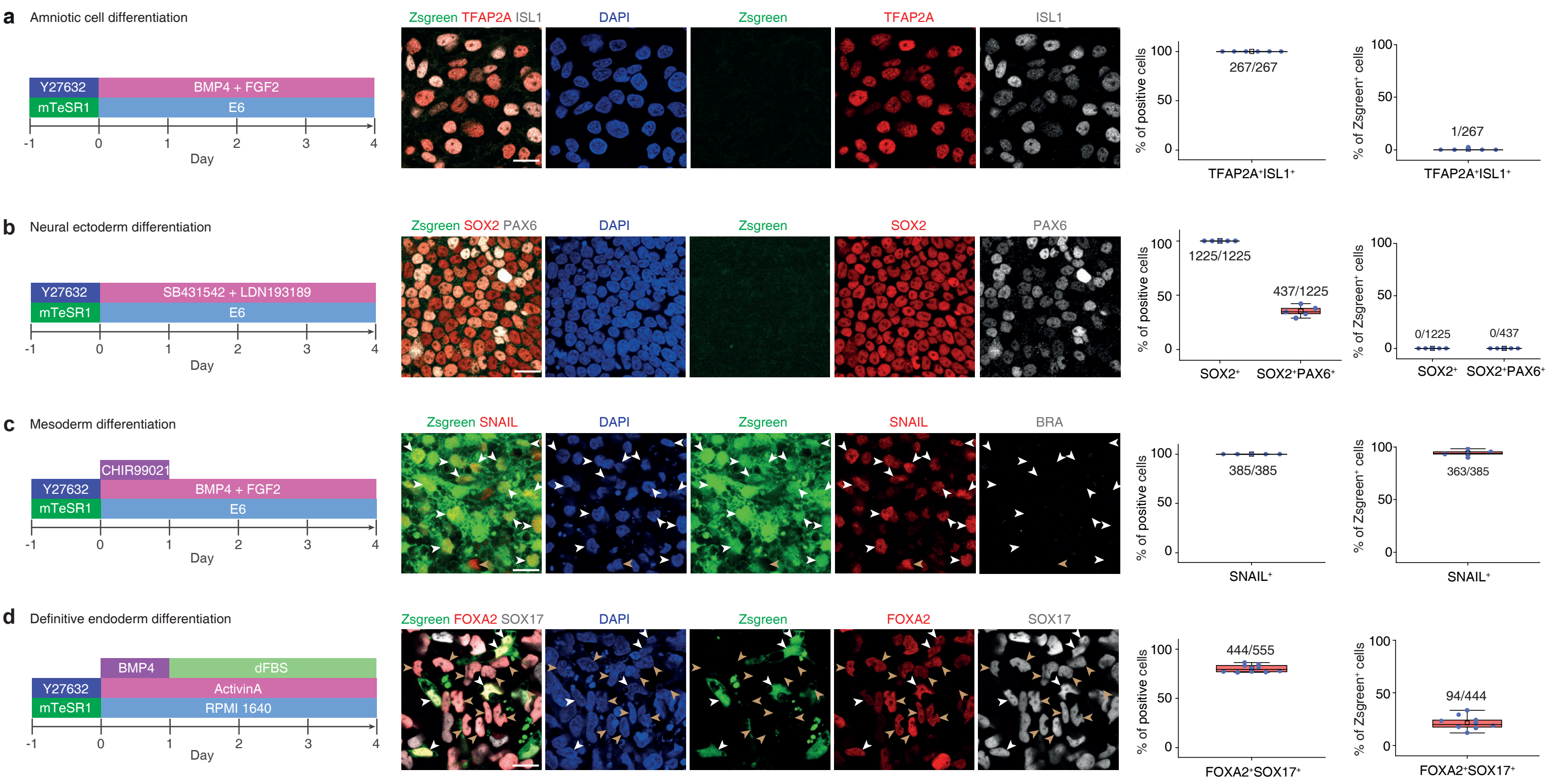

### Extended Data Figure 17

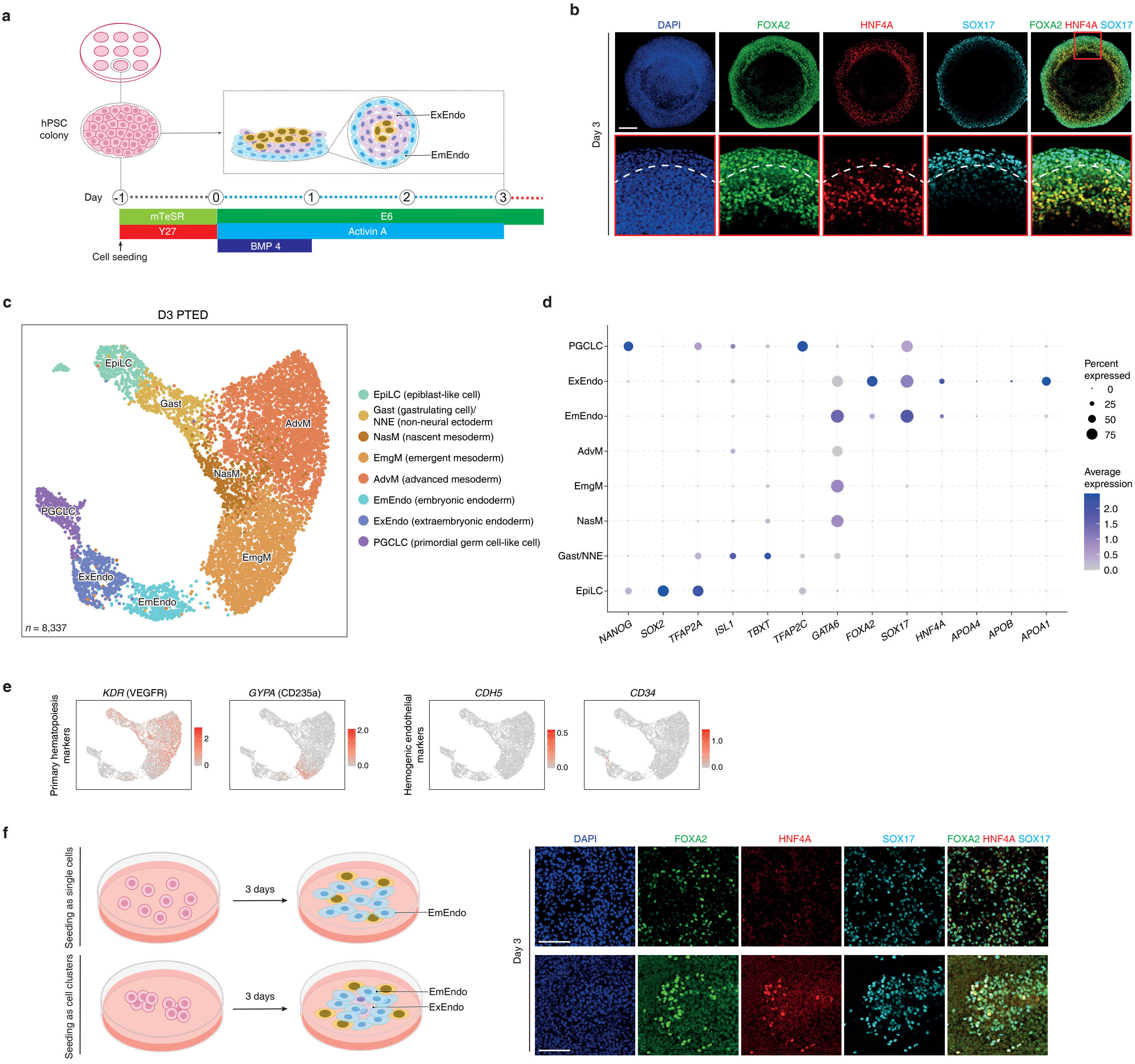

### Extended Data Figure 18

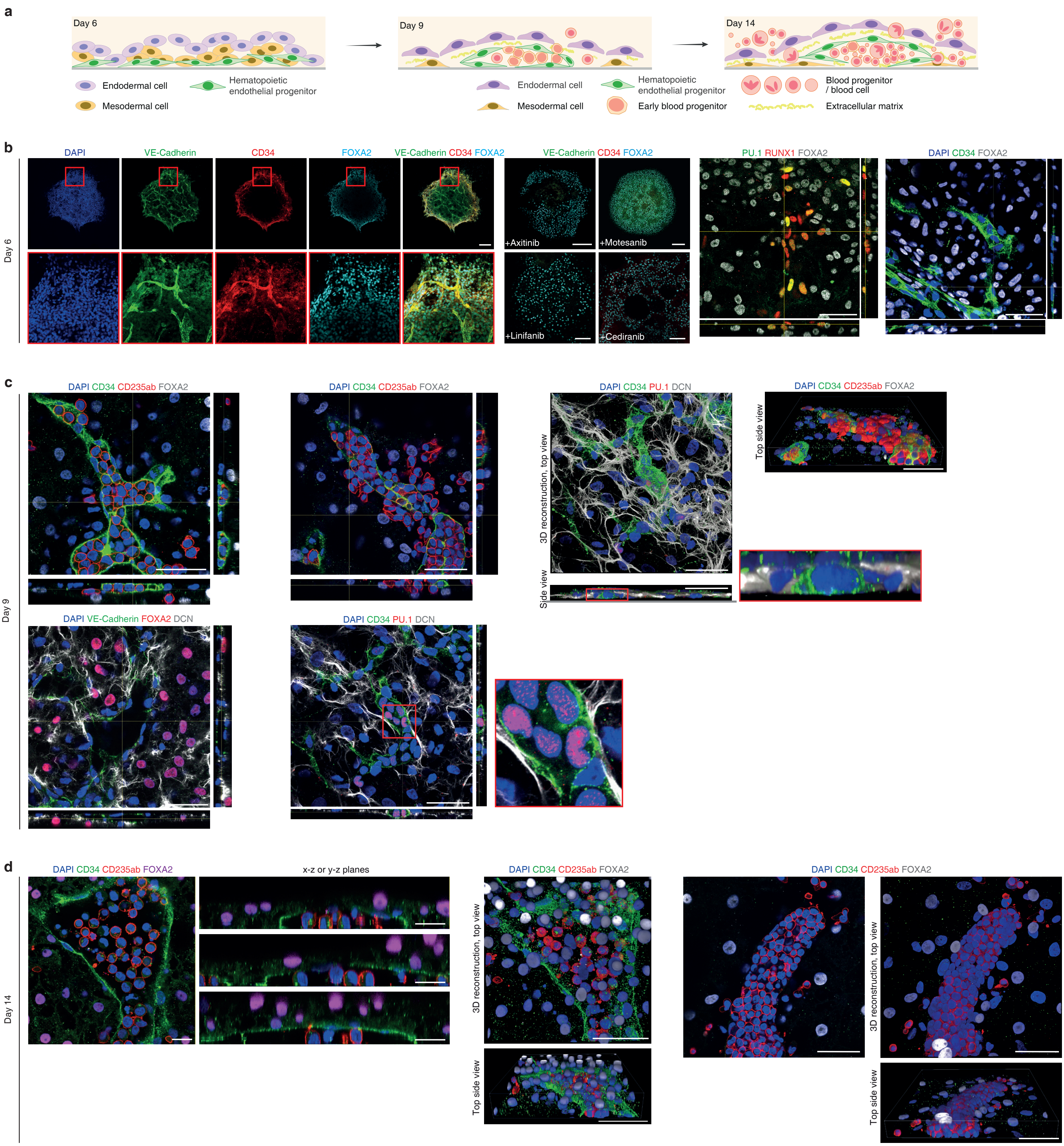

### Extended Data Figure 19

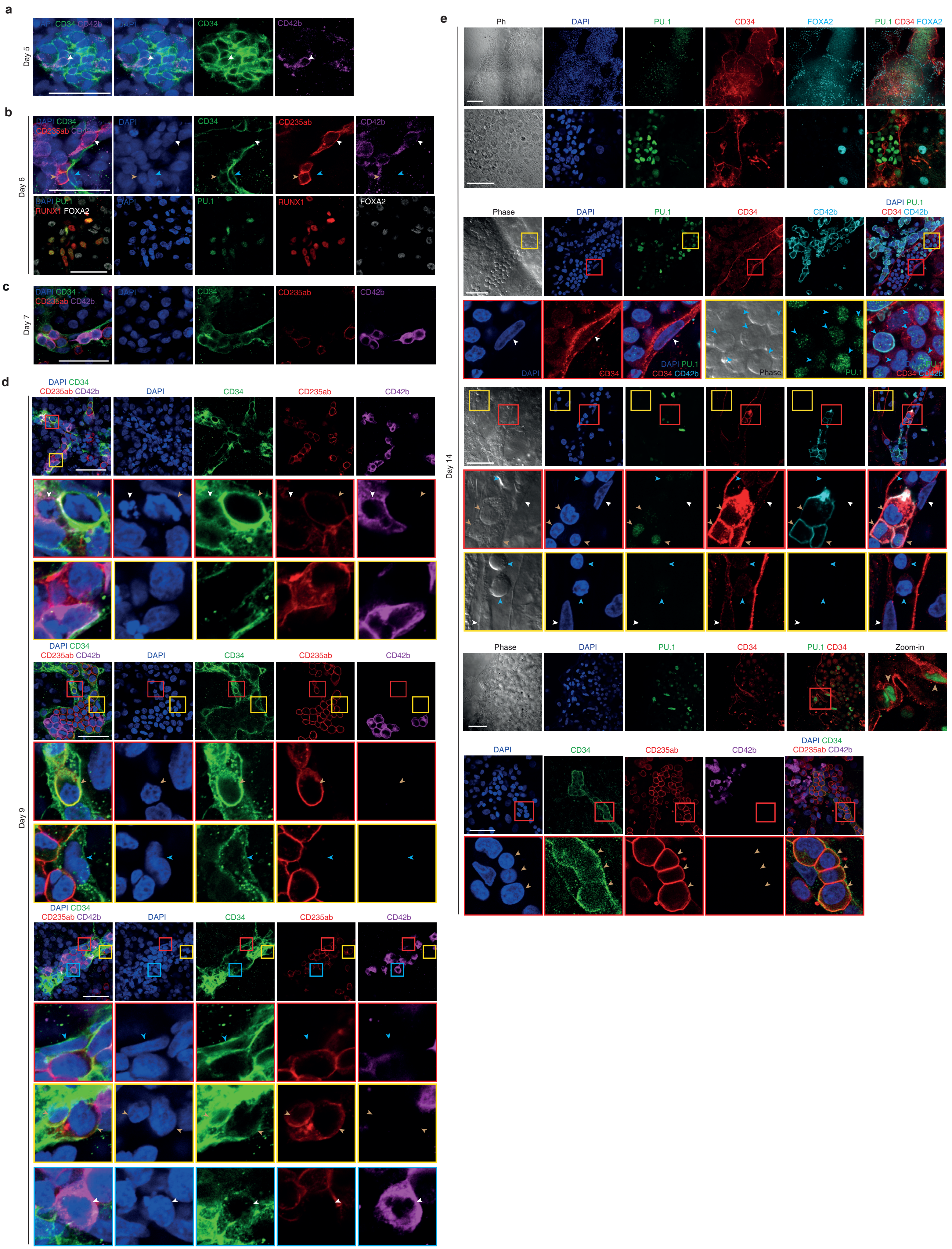

### Extended Data Figure 20

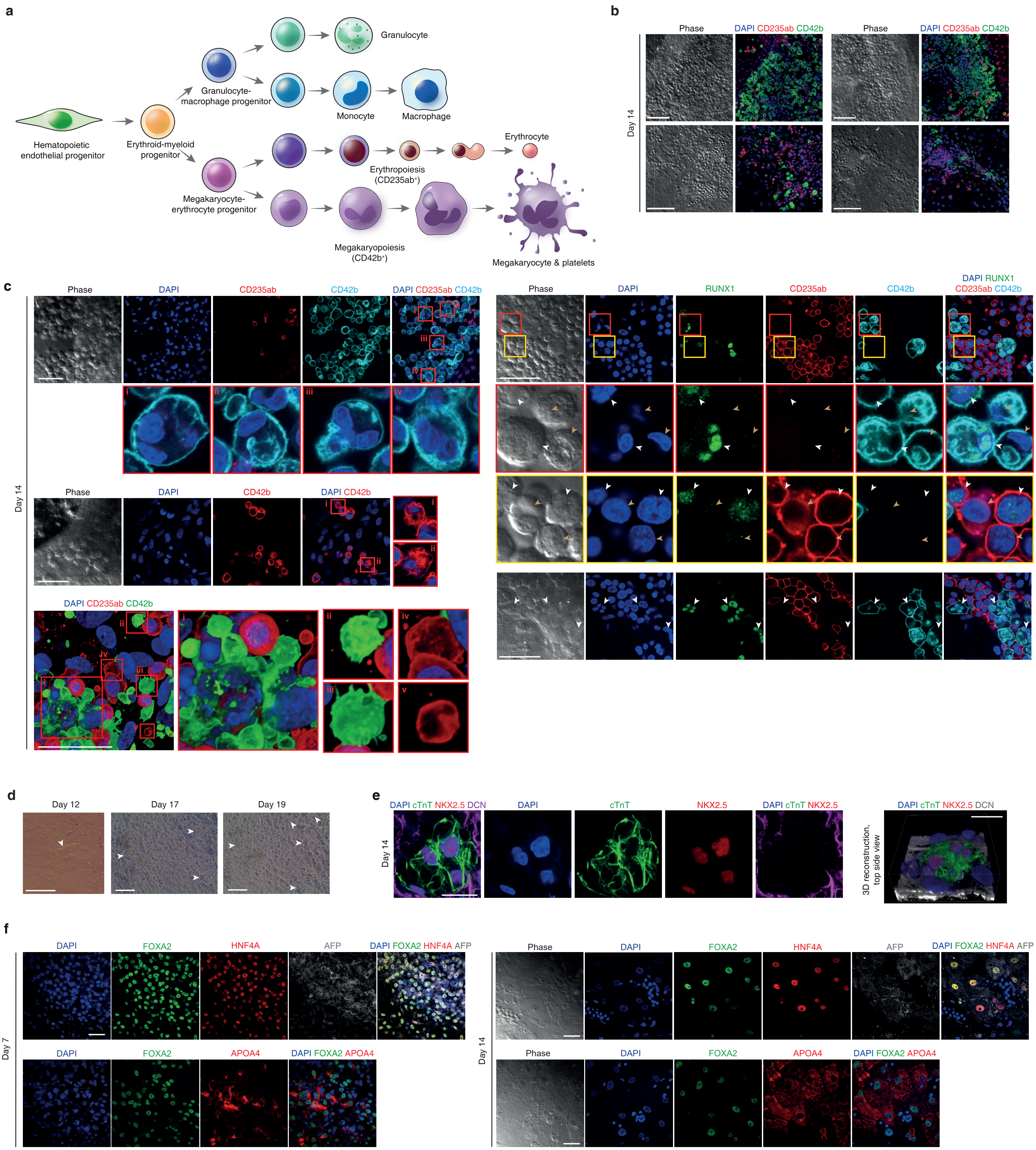

#### Extended Data Figure 21

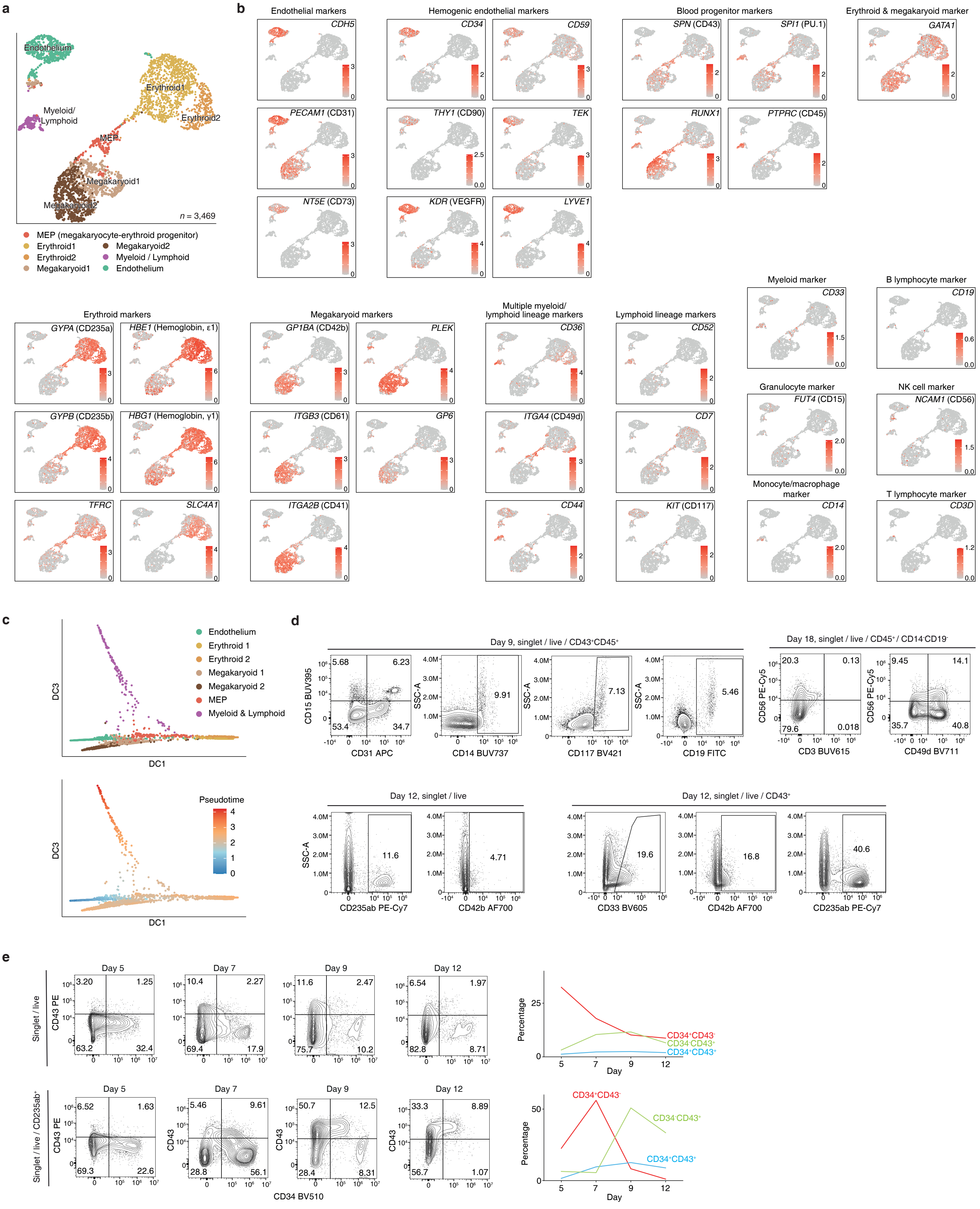

### Extended Data Figure 22

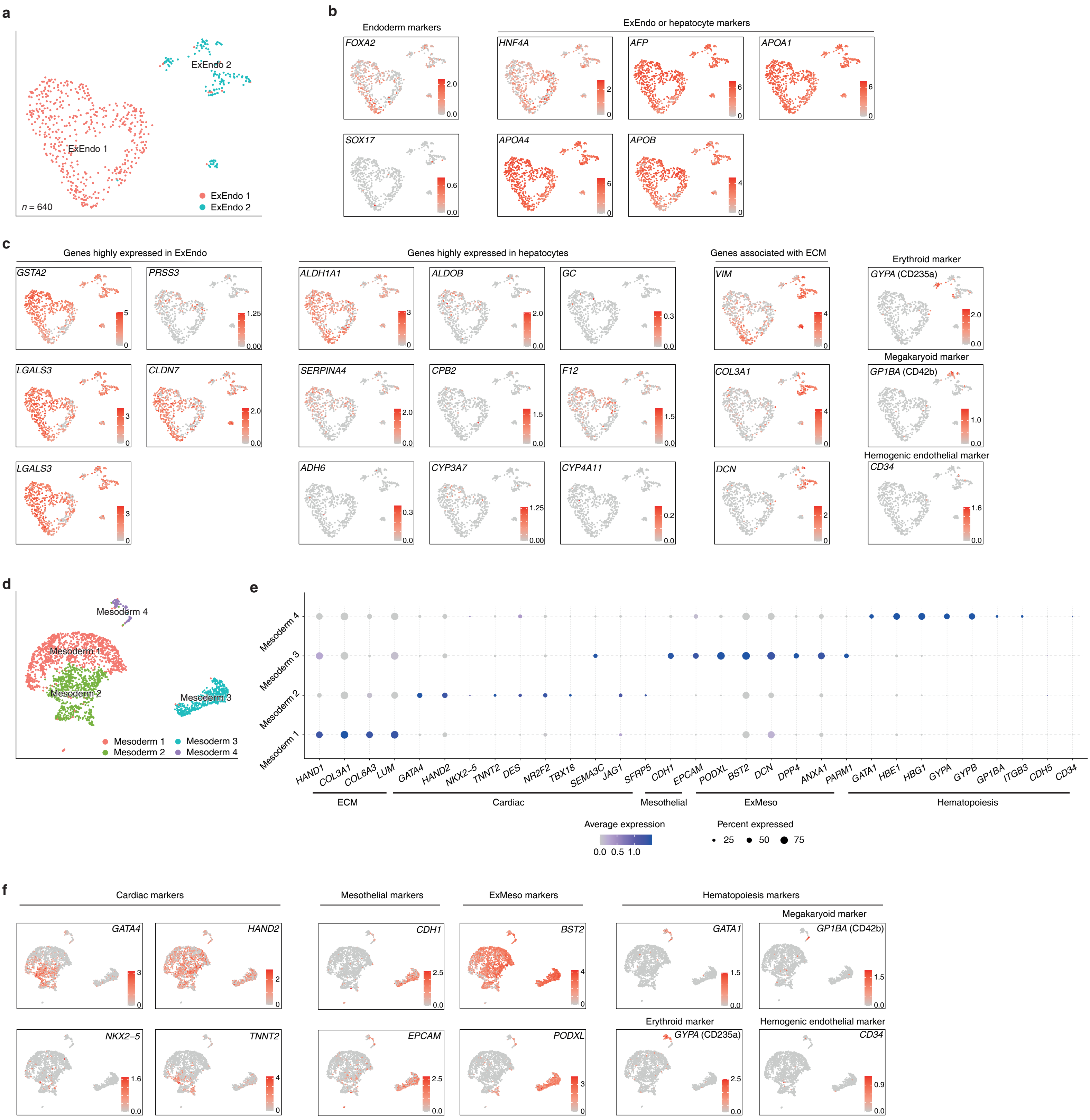

#### Extended Data Figure 23

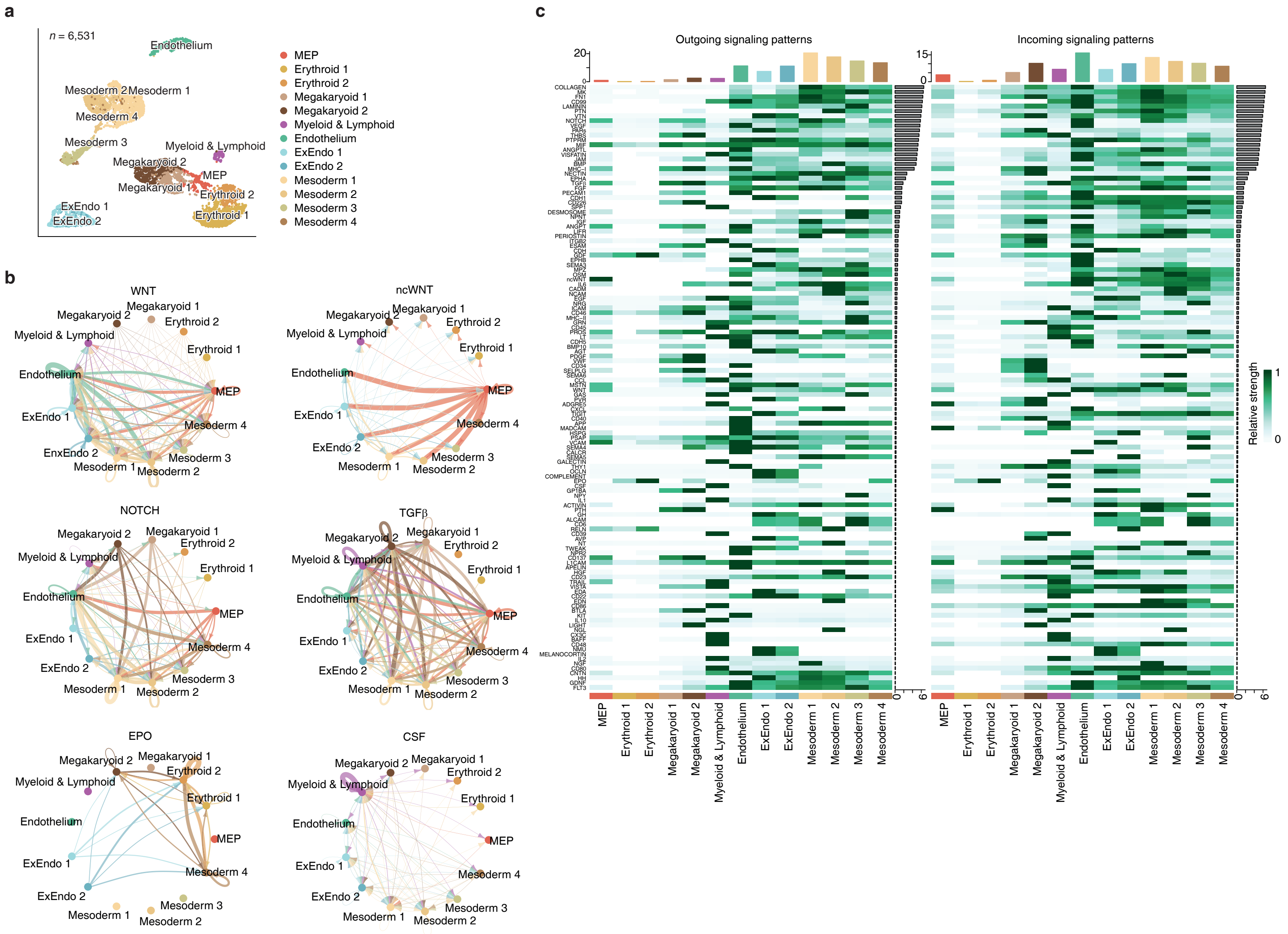
