## Supplementary figures and images for "A transgene-free, human peri-gastrulation embryo model with trilaminar embryonic disc-, amnion- and yolk sac-like structures"

### Flow cytometry gates.

#### Flow cytometry gating process

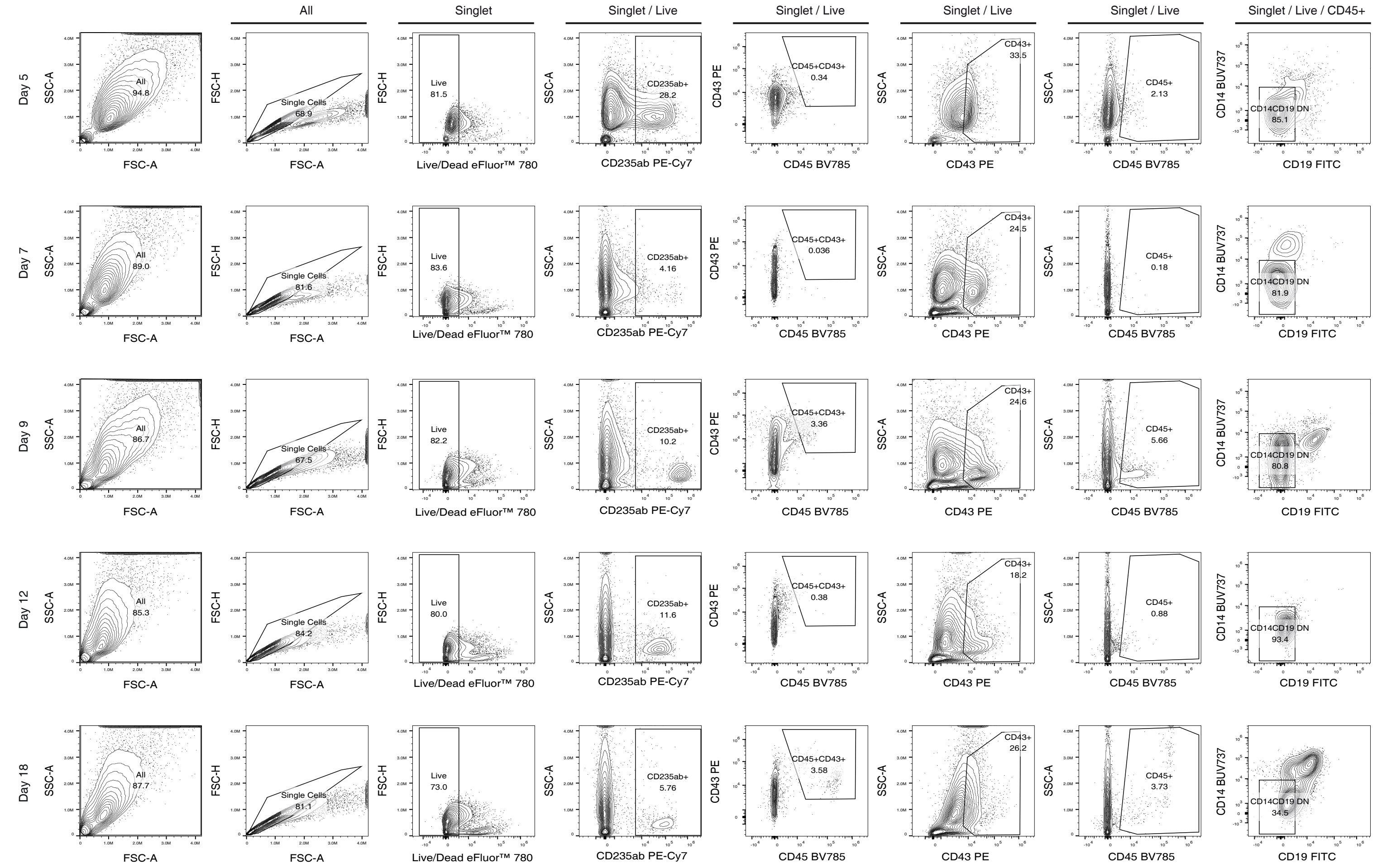
